# Resistance to chemical carcinogenesis induction via a dampened inflammatory response in naked mole-rats

**DOI:** 10.1101/2021.10.21.465383

**Authors:** Kaori Oka, Shusuke Fujioka, Yoshimi Kawamura, Yoshihiro Komohara, Takeshi Chujo, Koki Sekiguchi, Yuki Yamamura, Yuki Oiwa, Natsuko Omamiuda-Ishikawa, Shohei Komaki, Yoichi Sutoh, Satoko Sakurai, Kazuhito Tomizawa, Hidemasa Bono, Atsushi Shimizu, Kimi Araki, Takuya Yamamoto, Yasuhiro Yamada, Hiroyuki Oshiumi, Kyoko Miura

## Abstract

Naked mole-rats (NMRs) have a very low spontaneous carcinogenesis rate, which has prompted scientists to study their cancer resistance mechanisms in order to provide clues for human cancer prevention. Although cancer resistance in NMRs has been intensively investigated at the cellular level, it is still unknown how strongly resistant NMR individuals are to carcinogenesis and how NMR tissues respond to experimental carcinogenesis induction. Here, we show that NMRs exhibit extraordinary resistance against potent chemical carcinogenesis induction through a dampened inflammatory response. Although carcinogenic insults damaged skin cells of both NMRs and mice, NMR skin showed markedly lower immune cell infiltration and reduced induction of inflammatory genes. NMRs harbor loss-of-function mutations in receptor-interacting protein kinase 3 (*RIPK3*) and mixed lineage kinase domain-like (*MLKL*) genes, which are essential for necroptosis, a type of necrotic cell death that activates strong inflammation. A necroptosis-inducing stimulus did not increase death of NMR cells. After carcinogenic insults, leakage of the HMGB1, a marker of necrotic cell death, was not increased in NMR skin. In mice, inhibition or knockout of RIPK3 reduced immune cell infiltration and delayed the onset of chemical carcinogenesis. Therefore, necroptosis deficiency may serve as a cancer resistance mechanism via attenuating the inflammatory response in NMRs. Our study sheds light on the importance of a dampened inflammatory response as a non-cell-autonomous cancer resistance mechanism in NMRs. Further in vivo study of the unusual tissue immune system and carcinogenesis resistance of NMRs may lead to the development of new strategies to prevent carcinogenesis in humans.

**Significance Statement:** In contrast with intensive studies of cancer resistance mechanisms in naked mole-rats (NMRs) at the cellular level, little is known about how NMR individuals respond to carcinogenesis induction, despite the fact that cell-to-cell interactions in tissues regulate carcinogenesis in vivo. Here, we demonstrate that NMRs are remarkably resistant to chemical carcinogenesis induction and characteristically have attenuated tissue inflammatory responses to carcinogenic insults. NMRs have loss-of-function mutations in *RIPK3* and *MLKL* genes and thus cannot activate necroptosis, a type of inflammation-inducing cell death. RIPK3 inhibition in mice reduced immune cell infiltration in response to carcinogenic insults and delayed the onset of chemical-induced carcinogenesis. Our results highlight the importance of studies on dampened tissue inflammatory responses to understand cancer resistance of NMRs.

## Introduction

The naked mole-rat (NMR) is the longest-living rodent with a maximum lifespan of 37 years, despite being comparable size to the laboratory mice, and previous studies have reported that it is protected from age-associated declines in biological functions and aging-related disorders (1, 2). In particular, spontaneous carcinogenesis has rarely been observed in over 2,000 necropsies of captive NMR colonies (3, 4). This provides clear evidence of the cancer resistance properties of NMRs; however, to the best of our knowledge, it is the only evidence to date of their cancer resistance in vivo. On the other hand, several recent reports demonstrated that some NMRs spontaneously develop cancers (5–7). Therefore, it is unclear how strongly resistant NMR individuals are to carcinogenesis. Moreover, there is no report regarding the tissue response of NMRs to experimental induction of carcinogenesis in vivo.

Intracellular mechanisms that may contribute to cancer resistance in NMRs have been proposed (8–10); however, whether NMR cells have strong cell-autonomous cancer resistance is currently debatable. There are two reports that NMR cells, in contrast to mouse cells, do not transform upon introduction of HRasV12 and SV40 Large T antigen (9, 11), and conversely another report involving extensive experiments that they do (12). One limitation of previous studies is that the findings are based on in vitro experimental transformation of cultured fibroblasts and their xenografts in immunodeficient mice.

In vivo carcinogenesis includes an initiation stage, in which DNA damage results in the generation of mutant cells. This is followed by changes in the tissue microenvironment around the mutant cells, which comprises surrounding immune cells and stromal cells. Microenvironmental changes regulate various environmental factors and promote carcinogenesis in a promotion stage (13, 14). In particular, tissue inflammation induces further genetic and epigenetic alterations of mutant cells and strongly promotes carcinogenesis in a non-cell-autonomous manner (15–18). Therefore, previous studies on the cancer resistance of cultured NMR fibroblasts might pay little attention to the physiological context of in vivo carcinogenesis and might overlook relevant cancer resistance mechanisms in NMR tissues.

Here, we show that NMR individuals exhibit extraordinary resistance to carcinogenesis induction by chemical carcinogens in vivo. Notably, NMR skin tissues showed an unusual dampened inflammatory response to carcinogenic insults. NMRs harbor loss-of-function mutations in receptor-interacting protein kinase 3 (*RIPK3*) and mixed lineage kinase domain-like (*MLKL*), the regulators of necroptosis, a type of strong inflammation-activating cell death associated with various inflammatory diseases (19). Loss of necroptosis-inducing ability in NMRs may serve as a mechanism that attenuates inflammatory responses and suppresses carcinogenesis in vivo. This study highlights a dampened tissue inflammatory response as a non-cell-autonomous mechanism underlying carcinogenesis resistance in NMR individuals.

## Results

### NMRs show marked resistance to chemical carcinogenesis induction

The in vivo responses of NMR tissues to carcinogenic insults were examined using two types of chemical carcinogens, and the effects were compared with those in mice. First, mice and NMRs received intramuscular injections of 3-methylcholanthrene (3MC), a carcinogen in various rodent species (20–22) (Fig. 1A). After treatment, all mice developed fibrosarcomas within 24 weeks (9/9 tested animals; Fig. 1B, C). However, 3MC-treated NMRs did not develop tumors in a period of 114 weeks (0/9 tested animals; Fig. 1B, C, and Fig. S1A). Histopathological analysis (three animals) showed no obvious abnormalities (Fig. 1C). The remaining animals were kept alive, and no visible tumors were observed for 177 weeks. NMRs who received a subcutaneous injection of 3MC also did not develop visible tumors for 97 weeks (0/5 tested animals; Fig. 1D–F). No obvious histopathological abnormalities such as hyperplasia were detected although Ki67-positive cells tended to increase slightly (Fig. 1F and Fig. S1B). On the other hand, the mice developed severe skin ulcers and had to be euthanized within 10 weeks (Fig. S1C).

**Figure 1.**
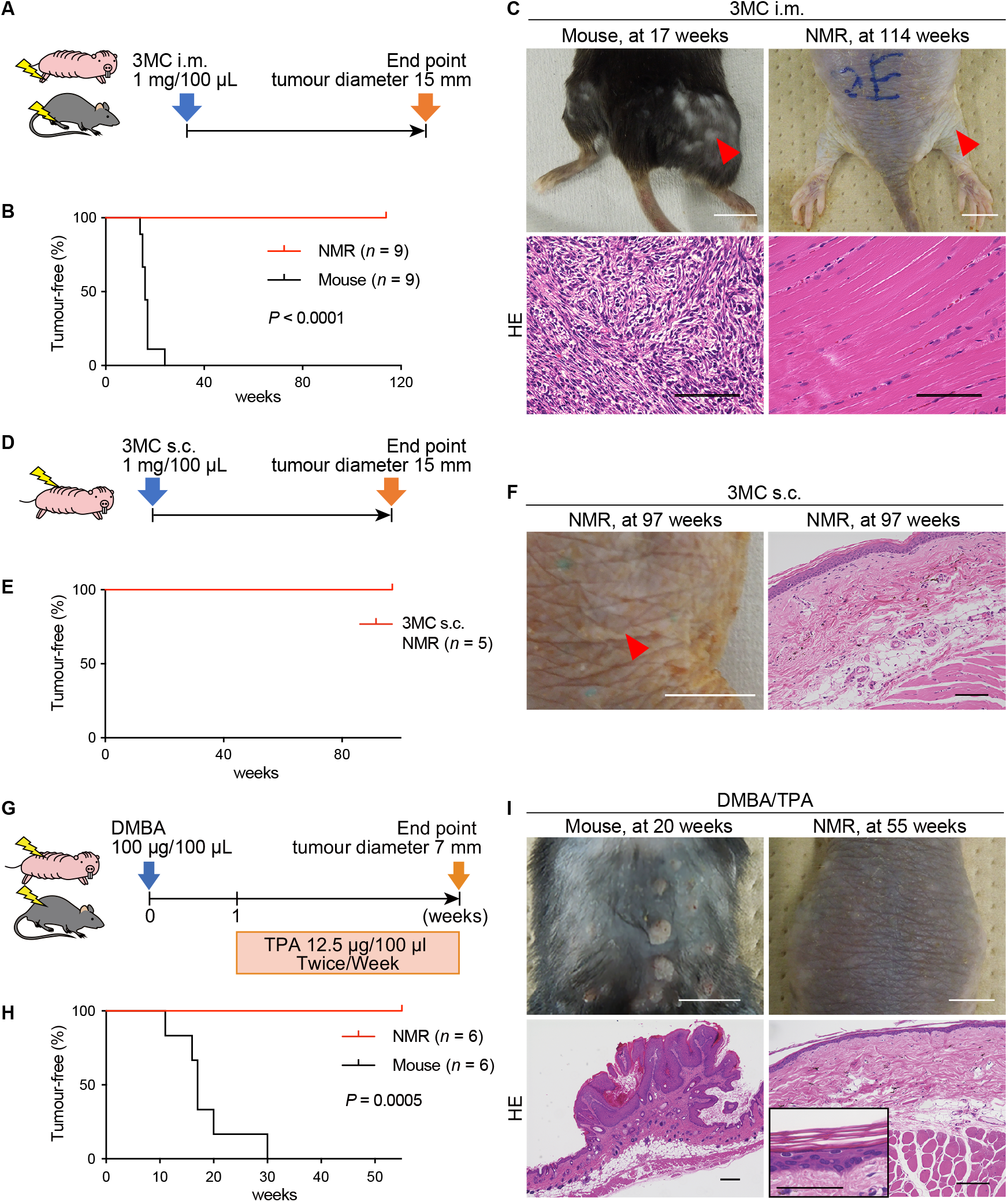
Naked mole-rats (NMRs) do not develop tumors in response to two types of chemical carcinogenesis induction. **A**, Schematic diagram for carcinogenesis induction by intramuscular (i.m.) injection of 1 mg 3-methylcholanthrene (3MC) into the hind limb. **B**, Kaplan–Meier curves of tumor-free mice and NMRs after i.m. injection of 1 mg 3MC. *n* = 9 animals per species. **C**, Gross appearance and hematoxylin and eosin (HE) staining of a mouse tumor at 17 weeks and NMR muscle at 114 weeks after i.m. 3MC injection. Red arrows indicate injection sites. Scale bars: 1 cm (upper) and 50 μm (lower). **D**, Schematic diagram for carcinogenesis induction by subcutaneous (s.c.) injection of 1 mg 3MC into the back skin. **E**, Kaplan–Meier curves of tumor-free NMRs after s.c. injection of 1 mg 3MC. *n* = 5 animals. **F**, Gross appearance and HE staining of NMR back skin at 97 weeks after s.c. injection of 1 mg 3MC. The red arrowhead indicates the injection site. Scale bars: 1 cm (gross) and 100 μm (HE). *n* = 5 animals. **G**, Schematic diagram for carcinogenesis induction by 7,12-dimethylbenz[a]anthracene (DMBA)/12-O-tetradecanoylphorbol-13-acetate (TPA) treatment on the back skin. **H**, Kaplan–Meier curves of tumor-free mice and NMRs after starting DMBA/TPA treatment. *n* = 6 animals per species. **I**, Gross appearance and HE staining of mouse papillomas at 20 weeks and NMR skin at 55 weeks after starting DMBA/TPA treatment. Scale bars: 1 cm (upper) and 100 μm (lower). Inset is a higher magnification of NMR skin (scale bar: 50 μm). Log-rank test for **B** and **H**.

Next, other carcinogens, namely, 7,12-dimethylbenz[a]anthracene (DMBA) and 12-*O*-tetradecanoylphorbol-13-acetate (TPA) (23), were administered to the back skin of mice and NMRs (Fig. 1G). All mice developed multiple papillomas within 30 weeks (6/6 tested animals; Fig. 1H, I). On the other hand, NMRs did not develop any visible tumors at 55 weeks, and histopathological analysis of skin biopsies showed no obvious abnormalities, although Ki67-positive cells increased slightly (0/6 tested animals; Fig. 1H, I, and Fig. S1D). These animals continued to receive TPA and did not develop tumors for 116 weeks.

### Carcinogens increase tissue damage in NMRs

To evaluate the early tissue responses to the carcinogen, 3MC was injected subcutaneously, and the effects were analyzed after 1 week (24) (Fig. 2A). NMR and mouse skin tissues showed increased phospho-histone H2A.X (pH2AX)-positive or 8-hydroxy-2′-deoxyguanosine (8-OHdG)- positive DNA-damaged cells in response to 3MC treatment, and TUNEL-positive dead cells were similarly increased (Fig. 2B, C and Fig. S2).

**Figure 2.**
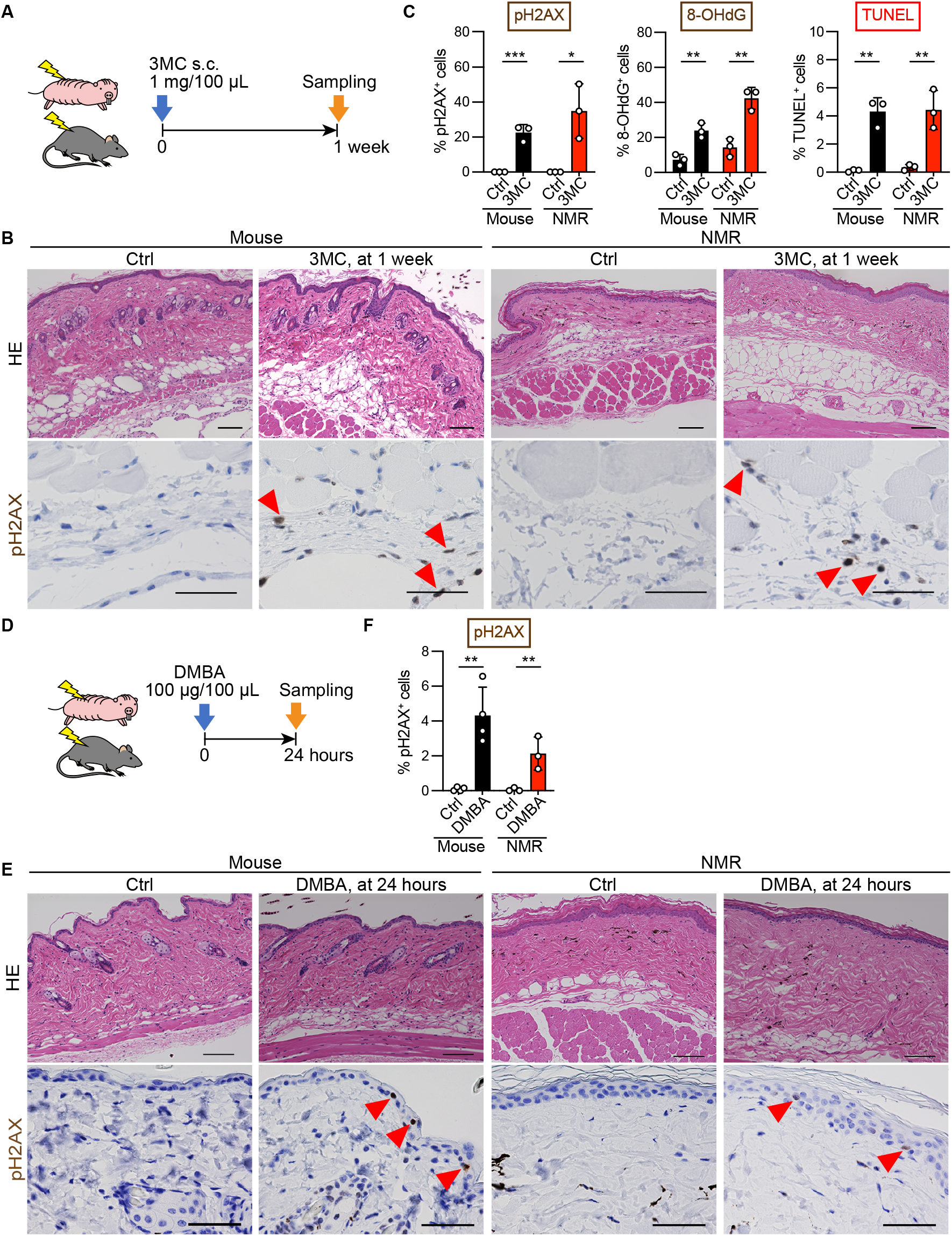
DNA damage and cell death increase in NMRs upon administration of carcinogens. **A**, Schematic diagram for investigating short-term responses to 3MC after subcutaneous (s.c.) injection into the back skin. **B**, HE staining and immunohistochemical staining for phospho-histone H2A.X (pH2AX, brown) of the skin of mice and NMRs at 1 week after s.c. injection of 3MC. Scale bars: 100 μm (HE) and 50 μm (pH2AX). Red arrowheads indicate positive cells. **C**, Quantification of pH2AX-, 8-hydroxy-2′-deoxyguanosine (8-OHdG)-, and TUNEL-positive cells after s.c. injection of 3MC. **D**, Schematic diagram for investigating short-term responses to DMBA treatment on the back skin. **E**, HE staining and immunohistochemical staining of pH2AX (brown) in the skin of mice and NMRs at 24 h after DMBA treatment. Scale bars: 100 μm (HE) and 50 μm (pH2AX). Red arrowheads indicate positive cells. **F**, Quantification of pH2AX-positive cells at 24 h after DMBA treatment. For quantification in **C** and **F**, data are presented as the mean ± SD of *n* = 3–4 animals. Unpaired *t*-test versus untreated control (Ctrl).

We then evaluated the tissue responses to DMBA at earlier stages (Fig. 2D). Similar to the effect of 3MC, DMBA treatment for 24 h significantly increased the number of pH2AX-positive DNA-damaged cells in both mouse and NMR skin (Fig. 2E, F). Taken together, these results demonstrate that treatment with carcinogenic agents increased tissue damage such as DNA damage and cell death in NMR tissues. Despite this increasing tissue damage, NMR individuals showed marked resistance against two types of chemical carcinogenesis induction.

### NMRs show dampened tissue inflammatory responses after carcinogenic insults

The effect of chemical carcinogens on immune cell infiltration was evaluated by immunostaining using pre-validated antibodies against CD45 (leukocytes), IBA1 (macrophages), myeloperoxidase (MPO, myeloid cells: neutrophils and macrophages), and CD3 (T cells) (Fig. S3A and Dataset S1). In mice, 3MC treatment significantly increased the number of CD45-, IBA1-, and CD3-positive immune cells at 1 and 3 weeks (Fig. 3A–C, Fig. S4, and Fig. S5A, B). These data reflect the infiltration of inflammatory immune cells after carcinogen treatment in mouse tissues as previously reported (16, 17). By contrast, 3MC-treated NMR skin showed very low levels of several types of immune cell infiltration at 1 and 3 weeks. Analysis of NMR skin at 97 weeks after 3MC treatment showed no significant increase in CD45-positive immune cells (Fig. 3A–C, Fig. S4, and Fig. S5). Similar to the results regarding the effects of 3MC, DMBA/TPA-treated NMR skin at 2 weeks showed a very small increase in the number of several immune cell types in contrast with mice (Fig. 3D–F and Fig. S6A, B). The accumulation of immune cells was markedly attenuated in NMR skin after 55 weeks of DMBA/TPA treatment, including after 108 rounds of treatment with TPA, a potent inflammatory agent (Fig. 3F and Fig. S6C). These results indicate that infiltration of inflammatory immune cells was much lower in NMRs than in mice after exposure to two types of chemical carcinogens.

**Figure 3.**
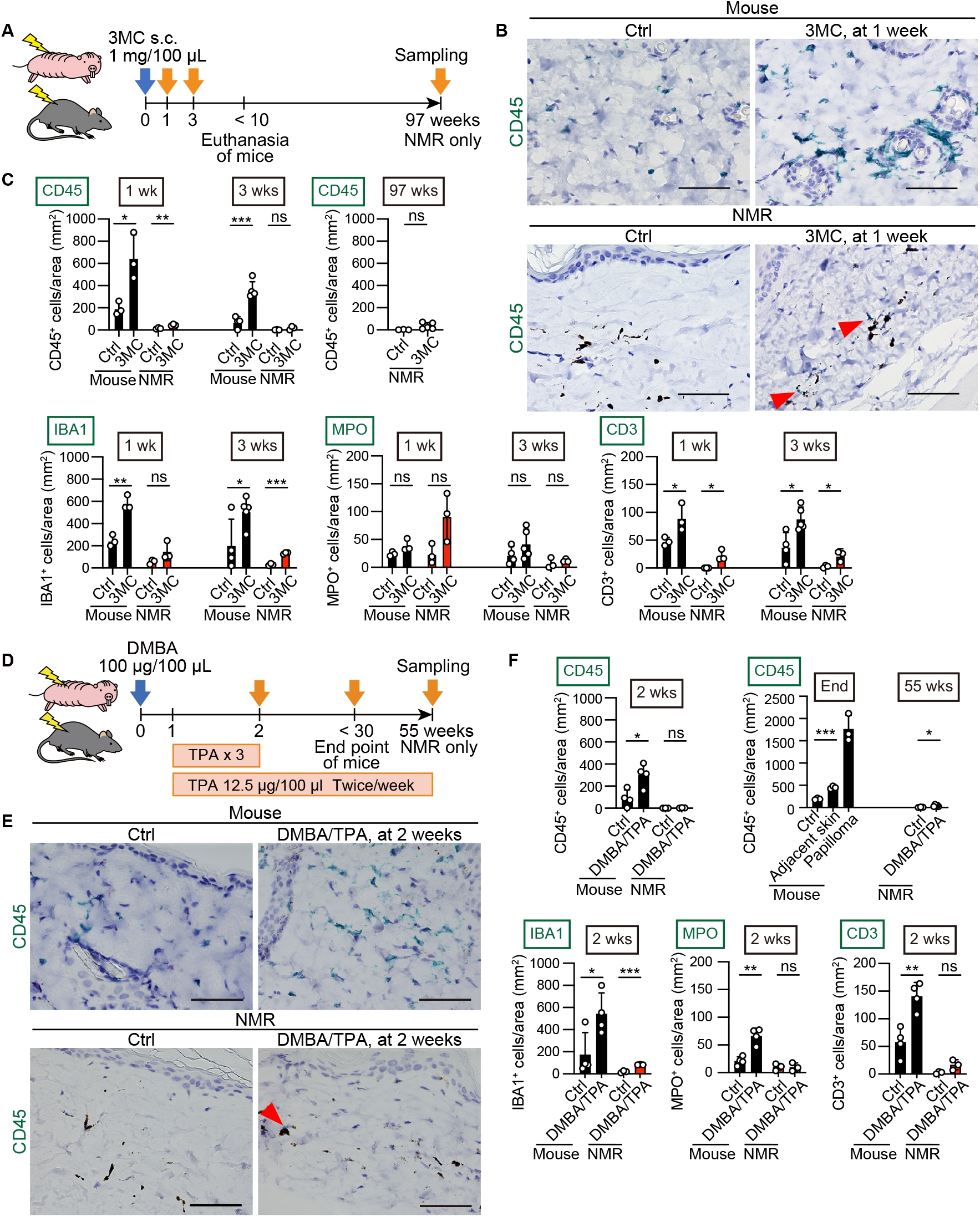
NMRs show attenuated infiltration of inflammatory immune cells into tissues upon administration of carcinogens. **A**, Schematic diagram for investigating immune cell infiltration into the skin after a subcutaneous (s.c.) injection of 3MC. **B**, Immunohistochemical detection of CD45 (green)-positive cells in skin sections 1 week after s.c. injection of 3MC. Black dots are melanin pigments in NMR dermis. Scale bar: 50 μm. Red arrowheads show positive cells in NMRs. **C**, Quantification of CD45-, IBA1-, MPO-, and CD3-positive cells per area in skin sections at 1, 3, and 97 (only for CD45) weeks after s.c. injection of 3MC. **D**, Schematic diagram for investigating immune cell infiltration into the skin after exposure to DMBA/TPA. **E**, Immunohistochemical detection of CD45 (green)-positive cells in skin sections at 2 weeks after exposure to DMBA/TPA. Black dots are melanin pigments in NMR dermis. Scale bar: 50 μm. The red arrowhead shows a positive cell in NMRs. **F**, Quantification of CD45-, IBA1-, MPO-, and CD3-positive cells per area in skin sections at 2 and 55 (only for CD45) weeks after exposure to DMBA/TPA. Mice were analyzed at the end point. “Adjacent skin” is the no-papilloma region from DMBA/TPA-treated mouse skin. For quantification in **C** and **F**, data are presented as the mean ± SD of *n* = 3–5 animals. Unpaired *t*-test versus untreated control (Ctrl).

Furthermore, we evaluated skin tissue responses to UV irradiation, which promotes carcinogenesis by inducing DNA damage and inflammation in mice (16, 25) (Fig. S7A). UVB irradiation significantly increased skin thickness, the number of pH2AX-positive DNA-damaged cells, TUNEL-positive dead cells, and cleaved caspase-3-positive apoptotic cells in both mouse and NMR skin, indicating that tissue damage increases in both species (Fig. S7B, C). In mouse skin, UV irradiation resulted in significant increases in the numbers of CD45-, IBA1-, and MPO-positive inflammatory immune cells, whereas in NMR skin, UV irradiation resulted in very small increases in the numbers of CD45- and CD3-positive immune cells, and no significant increase in IBA1- or MPO-positive immune cells (Fig. S8). These results indicate that the accumulation of inflammatory immune cells in response to various cancer-promoting stimuli is attenuated in NMRs, despite the induction of tissue damage such as DNA damage and cell death.

Next, we evaluated infection-associated inflammatory responses in NMRs (Fig. S9A). Subcutaneous injection of bacterial lipopolysaccharide (LPS), which reportedly activates NMR immune cells (26), increased interleukin-6 (*IL6*) expression, as well as the number of CD45- and MPO-positive immune cells in both mouse and NMR skin (Fig. S9B–D). Intraperitoneal LPS injection significantly increased the number of IBA1-positive cells in NMR livers (Fig. S10A). Thus, NMR immune cells can infiltrate into tissues in response to bacterial virulence factors. When co-cultured with dead NMR fibroblasts, NMR macrophages exhibited normal phagocytic activity of dead cells (Fig. S10B) despite showing reduced infiltration into carcinogen-treated damaged tissues.

During these experiments, we observed that the number of immune cells was lower in control NMR skin than in control mouse skin. To provide a context for this, we examined the number of tissue-resident immune cells in liver, skin, and intestine from mice, rats, guinea pigs, and NMRs. We found that the number of IBA1- or CD3-positive immune cells was lower in some NMR tissues, suggesting unique tissue immune homeostasis in NMRs (Fig. S11, S3).

To investigate the overall inflammatory responses to 3MC, UV, and LPS treatment, we performed global gene expression analysis using RNA-sequencing (RNA-seq). Changes in global gene expression or in selected ligand genes important for cell-to-cell communication, including many chemokines and cytokines (27), in response to the different treatments were greater in mouse skin than in NMR skin, and the upregulations were particularly large in the 3MC- and UV-treated mouse groups (Fig. S12A, B and Dataset S2). Cell type enrichment analysis using xCell (28) showed that all treatments significantly increased several immune cell enrichment scores in mouse skin (Fig. S12C and Dataset S3). By contrast, in NMR skin, 3MC and UV treatment did not significantly change the immune cell enrichment scores, whereas LPS did (Fig. S12C and Dataset S3). These results are consistent with those of immunohistochemical analyses of immune cell markers (Fig. 3 and Figs. S4, S5, S6, S8, S9), and confirm that inflammatory responses to cancer-promoting stimuli are attenuated in NMR tissues.

### Loss-of-function mutations in necroptosis regulators in NMRs may contribute to the attenuated tissue inflammatory response and carcinogenesis resistance

To examine the mechanisms underlying the different responses of NMR tissues to cancer-promoting stimuli, we analyzed differentially expressed genes (DEGs) in response to 3MC and UV treatment that differed from DEGs in response to LPS treatment between mice and NMRs. We selected genes that were species-specifically upregulated by >2 fold in NMRs or mice after both 3MC and UV treatment, but that were not commonly upregulated after LPS treatment (Fig 4A, blue-filled area, collectively termed 3MC-UV Mouse-DEGs and Fig. S13A, purple-filled area, collectively termed 3MC-UV NMR-DEGs). Enrichment analysis of the selected DEGs was performed using Metascape (29). Among 3MC-UV NMR-DEGs, genes related to the Kyoto Encyclopedia of Genes and Genomes (KEGG) pathway “p53 signaling pathway” were highly enriched, suggesting activation of the p53 pathway in 3MC- and UV-treated NMR skin (Fig. S13A and Dataset S4). This was consistent with the immunostaining data showing increased DNA damage and cell death in 3MC- and UV-treated NMR skin (Fig. 2B, C, and Fig. S2, S7). Among 3MC-UV Mouse-DEGs, genes related to the KEGG pathway “Cytokine-cytokine receptor interaction” and the gene ontology (GO) term “Leukocyte migration” were highly enriched, indicating the activation of inflammatory responses in 3MC- and UV-treated mouse skin (Fig. 4A and Dataset S4).

**Figure 4.**
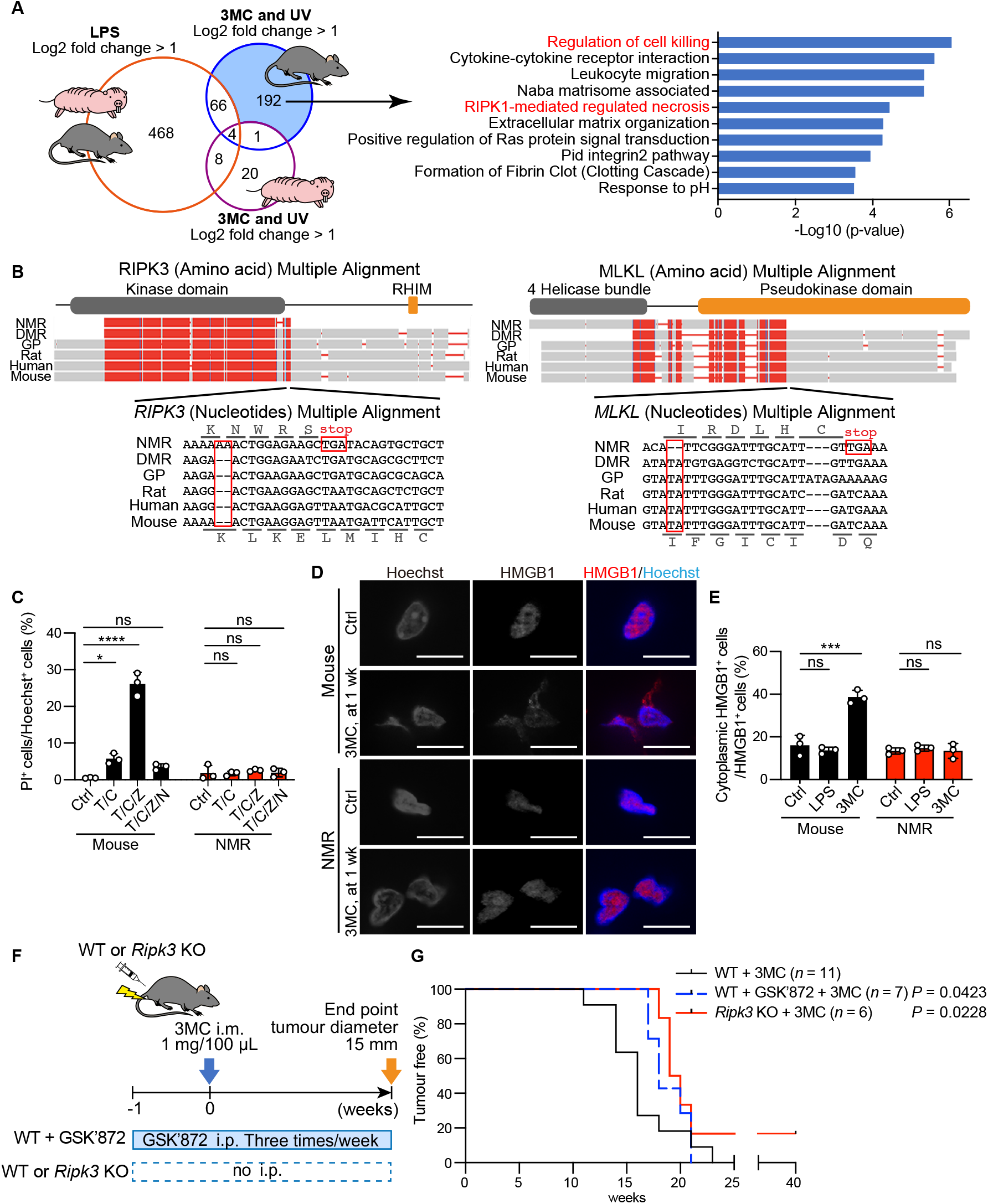
Loss of necroptosis regulators may contribute to an attenuated tissue inflammatory response and resistance to carcinogenesis in NMR individuals. **A**, Venn diagram showing the number of genes upregulated in both mouse and NMR skin upon lipopolysaccharide (LPS) treatment; genes upregulated specifically in mouse or NMR skin upon exposure to 3MC (1 week) and UV; and enriched pathways of 3MC-UV Mouse-DEGs. **B**, Multiple alignments of receptor-interacting kinase 3 (*RIPK3*) and mixed lineage kinase domain-like (*MLKL*) sequences from the NMR, Damaraland mole-rat (DMR), guinea pig (GP), rat, human, and mouse. Frame-shift mutations and premature stop codons in the NMR sequence are boxed. Reading frames for the NMR and mouse sequences are indicated. The functional domains are shown above the alignments. **C**, Cell death analysis in fibroblasts treated with a combination of TNF-α (T), cycloheximide (C), z-VAD-fmk (Z), or Nec-1 (N). Data are presented as the mean ± SD of *n* = 3 independent experiments. **D**, Immunofluorescence staining of high mobility group box-1 protein (HMGB1, red) in skin at 1 week after 3MC-injection. Nuclei: blue. Scale bar: 10 μm. **E**, Quantification of cytoplasmic HMGB1 in skin after each treatment. Data are presented as the mean ± SD of *n* = 3 animals for each species. One-way ANOVA with Tukey’s multiple comparison test for **C** and Dunnett’s multiple comparisons test versus untreated control (Ctrl) for **E**. **F**, Schematic diagram for carcinogenesis induction by intramuscular (i.m.) injection of 3MC with intraperitoneal (i.p.) injection of GSK’872 in mice or i.m. injection of 3MC in *Ripk3* knockout (KO) mice. **G**, Kaplan–Meier curves of tumor-free mice (*n* = 11 [for wild-type, WT], *n* = 7 [for GSK’872], or *n* = 6 [for *Ripk3* KO] animals). *P* = 0.0423 for GSK’872 and *P* = 0.0228 for *Ripk3* KO; Gehan–Breslow–Wilcoxon test.

Notably, among 3MC-UV Mouse-DEGs, genes related to the KEGG pathways “RIPK1-mediated regulated necrosis” (necroptosis) and “Regulation of cell killing” were the most significantly enriched (Fig. 4A and Dataset S4), which were not observed among 3MC-UV NMR-DEGs. Necroptosis, a type of programmed necrotic cell death, triggers inflammation, and promotes colon and pancreatic cancer development (14, 30–32). Thus, we hypothesized that inactivation of necroptosis in NMRs may underlie the attenuated inflammatory responses in NMRs, possibly leading to cancer resistance.

The RIPK1-RIPK3 complex induces necroptosis via the necroptosis effector, MLKL (33–35). We found that the NMR genome harbors a two-nucleotide insertion in the *RIPK3* gene and a two-nucleotide deletion in the *MLKL* gene, both of which cause frame-shift mutations and introduce premature stop codons (Fig. 4B). These alterations remove the RHIM domain in RIPK3 and the pseudokinase domain in MLKL, which are both functionally essential for necroptosis in other mammalian species (36). Because NMR *RIPK3* and *MLKL* genes have premature stop codons located before the final exon, the transcripts from these two genes are putative targets for nonsense-mediated mRNA decay (NMD) (37). As NMR *RIPK3* mRNA was expressed in the skin (Fig. S13B, C), we examined whether NMR *RIPK3* is degraded by NMD. RT-qPCR analysis of NMR fibroblasts treated with actinomycin D (ActD, a transcriptional inhibitor) and/or cycloheximide (CHX, a translational inhibitor that potently inhibits NMD) showed that NMR *RIPK3* transcripts exhibited relatively low steady-state levels after ActD treatment, whereas the *RIPK3* mRNA level increased upon CHX treatment (Fig. S13D). This result indicates that NMR *RIPK3* mRNA is degraded by NMD. NMR *MLKL* mRNA expression was not detected in the skin (Fig. S13E–G). Although previous studies have shown that only the N-terminal 4-alpha helical bundle domain of MLKL can cause spontaneous cell death depending on the cellular context (38–40), NMR MLKL could not induce spontaneous cell death (Fig. S14A, B). Thus, the genes essential for necroptosis induction are likely to be defective in NMRs.

To evaluate whether necroptosis is impaired in NMRs, we performed experimental necroptosis induction in vitro. In mouse fibroblasts, treatment with tumor necrosis factor-α (TNF-α), CHX, and z-VAD-fmk (caspase inhibitor) caused massive cell death, which was inhibited by necrostatin-1 (Nec1, RIPK1 inhibitor), as previously reported (41), indicating activation of necroptosis (Fig, 4C). In contrast to mouse fibroblasts, NMR fibroblasts did not show increased cell death in response to TNF-α + CHX or TNF-α + CHX + z-VAD-fmk, although TNF-α upregulated *IL6* (42) as observed in mice (Fig. 4C and Fig. S14C). These results suggest that NMR cells are incapable of inducing TNF-α-mediated necroptosis and apoptosis, although they are capable of inducing DNA damage-induced caspase-3-dependent apoptosis (Fig. S14D–F). In mice, RIPK3 is important for the induction of both necroptosis and TNF-induced apoptosis mediated by RIPK1 (43). Thus, the loss-of-function mutation of *RIPK3* in NMRs may contribute to their inability to undergo necroptosis and TNF-induced apoptosis mediated by RIPK1.

Generally, necroptosis triggers inflammation through the release of various cellular components such as high mobility group box-1 protein (HMGB1) (44), which can be observed during cancer progression (16, 17). 3MC, DMBA, and UV treatment did not significantly alter cytoplasmic HMGB1 translocation in NMR skin, in contrast to the significant increase observed in mouse skin (Fig. 4D, E and Fig. S15). These results further support the idea that the inability to induce necroptosis in NMRs may contribute to the dampened immune cell responses to carcinogenic stimuli.

Next, we assessed whether the necroptosis inhibitor (45) GSK’872, a RIPK3 inhibitor, or disruption of the *Ripk3* gene could suppress the 3MC-induced inflammatory response in mice and impede carcinogenesis as observed in NMRs (Fig. 4F and Fig. S16, S17A). The results of western blotting showed that 3MC treatment activated the MLKL protein (as indicated by MLKL phosphorylation), and MLKL activation was suppressed by GSK’872 treatment (Fig. S17B). GSK’872 and disruption of the *Ripk3* gene significantly suppressed cytoplasmic HMGB1 translocation (Fig. S17C), indicating that necroptosis was successfully suppressed. In addition, both manipulations reduced the infiltration of inflammatory immune cells in 3MC-treated mouse skin (Fig. S17D). Finally, we evaluated the effect of GSK’872 treatment or *Ripk3* knockout on 3MC-induced chemical carcinogenesis in mice (Fig. 4F). Continuous administration of GSK’872 or disruption of the *Ripk3* gene significantly delayed the onset of carcinogenesis in 3MC-treated mice (Fig. 4G; *P* = 0.0423 for GSK’872; *P* = 0.0228 for *Ripk3* KO mice; Gehan–Breslow–Wilcoxon test). Thus, in mice, the suppression of the necroptosis regulator attenuated immune cell infiltration and chemical carcinogenesis. This result is consistent with our assumption that the absence of necroptosis regulators in NMRs may contribute to the reduced inflammatory response and resistance to chemical carcinogenesis.

## Discussion

In this study, NMRs showed marked resistance to two types of chemical carcinogenesis induction in vivo. The distinctive feature of the NMR tissue response to carcinogenic insults was an unusual dampened inflammatory response, which may serve as a non-cell-autonomous cancer resistance mechanism in NMR individuals. Inhibition of RIPK3 in mice resulted in the reduced inflammatory response and the delayed onset of carcinogenesis (Fig. S18). Therefore, we propose that one of the mechanisms underlying the attenuated tissue inflammatory response and remarkable cancer resistance of NMRs may be specific loss-of-function mutations in the necroptosis regulators *RIPK3* and *MLKL*.

In a different cancer-resistant rodent, the blind mole-rat, the same dose of 3MC causes low frequency carcinogenesis (~9%) (22, 46). Thus, NMRs are especially resistant to chemical carcinogenesis. In contrast to NMRs, 3MC induces massive inflammation in blind mole-rats (22). This distinct difference in inflammatory responses between NMRs and blind mole-rats, both of which show spontaneous cancer resistance and a high DNA repair capacity (47–49), may contribute to the differences in resistance to in vivo carcinogenesis induction.

The attenuated cancer-promoting tissue inflammatory response may act as a gatekeeper to prevent carcinogenesis in NMRs. This is supported by previous reports showing that toll-like receptor 4 knockout mice exhibit carcinogenic resistance owing to dampened inflammatory responses (16, 17). Other mechanisms besides the deficiency in necroptosis, especially those related to immune cell characteristics, might also contribute to the unique inflammatory response in NMRs. Immune homeostasis in NMRs may be unusual because the resident immune cells, which contribute to the attenuated immune response and to cancer resistance in mice (50, 51), were less numerous in some tissues of NMRs than in those of other rodent species (Fig. S11). Moreover, a single cell RNA-seq study of immune cells revealed the unique immune system of NMRs, which is characterized by a lack of natural killer cells (26). Recent in silico and in vitro studies have shown that cancer-resistant bats lack certain immunity-related genes (52, 53). Future studies examining the immune system of cancer-resistant animals should improve our understanding of their cancer resistance mechanisms.

The type of cell death and its modulation play critical roles in the regulation of inflammation and homeostasis in vivo. In this study, caspase-3-dependent apoptosis occurred in NMRs, whereas necroptosis did not (Fig. 4C–E, Fig. S7B, C, Fig. S14D–F, and Fig. S15). Since the pro-inflammatory potential of necroptosis is markedly higher than that of apoptosis (44), the suppression of necroptosis may contribute substantially to the attenuation of the inflammatory response in NMR tissues as in RIPK3 inhibited/disrupted mice. Necroptosis is involved in a variety of inflammatory age-related diseases/disorders, such as ischemia-reperfusion injury, atherosclerosis, and neurodegenerative diseases. On the other hand, necroptosis also plays an important role in innate immunity during infectious diseases (19). Notably, NMRs are resistant not only to cancer, but also to aging-related declines in biological function, neurodegenerative disease, and ischemia-reperfusion injury, although they exhibit high susceptibility to herpes virus infection (3, 54–56). It is possible that deficiency in necroptosis induction may constitute an important part of the mechanisms responsible for the unusual characteristics of NMRs. It would also be interesting to study how other types of cell death, such as ferroptosis and pyroptosis, are regulated in NMRs.

Recent studies have shown that RIPK3 is involved not only in the induction of necroptosis and RIPK1-mediated apoptosis, but also in the activation of the NLRP3 inflammasome, maturation of IL-1β, and production of inflammatory cytokines, all of which are not directly activated via necroptosis (57). MLKL contributes to various biological functions, such as endosomal trafficking and extracellular vesicle formation, in addition to the induction of necroptosis and inflammatory cytokines (58). Therefore, the loss-of-function mutations of *RIPK3* and *MLKL* in NMRs may affect not only necroptosis, but also the attenuation of the tissue inflammatory response via suppression of the NLRP3 inflammasome and various other biological processes in vivo. This will require further analysis.

In addition to the role of cancer-promoting inflammation, the generation of mutant cells is also crucial for the initiation of carcinogenesis (59). Although carcinogen treatment damaged DNA and cells in NMR skin (Fig. 2 and Fig. S2, S7), it is possible that NMRs are protected against mutant cell generation or efficiently eliminate mutant cells. Possible explanations include 1) inhibition of mutant cell generation via several mechanisms, such as the previously reported efficient DNA double-strand break repair (48), or 2) elimination of mutant cells by unknown mechanisms, which may synergistically contribute to the in vivo cancer resistance of NMRs.

The present results demonstrate the importance of research into the unusual tissue immune response and extraordinary resistance to carcinogenesis of NMR individuals. Further insight into the tissue responses of the NMR to carcinogenic insults may lead to the development of new anticancer strategies for humans.

## Materials and Methods

### Animals

NMRs were maintained at Kumamoto University and Hokkaido University. All NMRs (8–31 months) used in this research were raised in rooms that were maintained at 30°C ± 0.5°C and 55% ± 5% humidity with 12 h light and 12 h dark cycles (10). The NMRs used in this study are listed in Dataset S5. Male C57BL/6N mice (8–10 weeks) were purchased from CLEA Japan, and *Ripk3* knockout (KO) mice were generated by deletion of the *Ripk3* gene. Wild-type mice and KO mice were kept in rooms that were maintained at 24.5°C ± 1.5°C and 50% ± 10% humidity with 12 h light and 12 h dark cycles. Male rats (Wistar, 6 months) and guinea pigs (Hartley, 6 months) were purchased from Japan SLC. The Ethics Committees of Kumamoto University (approval no. A30-043 and A2020-042) and Hokkaido University (14-0065) approved all procedures, which were in accordance with the Guide for the Care and Use of Laboratory Animals (United States National Institutes of Health, Bethesda, MD, USA).

### Generation of *Ripk3* knockout mice and genotyping

*Ripk3* KO mice were generated by introduction of the Cas9 protein (317–08441; NIPPON GENE), tracrRNA (GE-002; FASMAC), synthetic crRNA (FASMAC), and ssODN into C57BL/6N fertilized eggs by electroporation. For generating the *Ripk3* KO allele, the synthetic crRNAs were designed according to the sequence AAGAGAGACTGGCTATCGTG (GGG) of the 5′ upstream region of *Ripk3* and ACTAGGAGAGGATCCCACTG (AGG) in the *Ripk3* intron 9. The ssODN 5′- CGACTTTCTTTCGTTGTGTGACCTCAGttttatttGATAGCCAGTCTCTCTTGGACCCCTTAGCTCC ACC-3′ was used as a homologous recombination template.

The electroporation solution contained 10 μM tracrRNA, 10 μM synthetic crRNA, 0.1 mg/mL Cas9 protein, and 1 μg/μL ssODN in Opti-MEM I Reduced Serum Medium (31985062; Thermo Fisher Scientific). Electroporation was performed using the Super Electroporator NEPA 21 (NEPA GENE) on glass microslides with round wire electrodes (1.0 mm gap [45–0104; BTX]). Four steps of square pulses were applied (1], three times of 3 mS poring pulses with 97 mS intervals at 30 V; 2], three times of 3 mS polarity-changed poring pulses with 97 mS intervals at 30 V; 3], five times of 50 mS transfer pulses with 50 mS intervals at 4 V with 40% decay of voltage per pulse; 4], five times of 50 mS polarity-changed transfer pulses with 50 mS intervals at 4 V with 40% decay of voltage per pulse).

The targeted *Ripk3* KO allele in F0 mice was identified by genomic PCR using the following primers: Ripk3 KO F: 5′- AGCGACACCTTGTGATCTCC-3′ and Ripk3 KO R: 5′- CTGGCCCAAGACAACCCTTA -3′ for the knockout allele (396 bp); Ripk3 Wild F: 5′- GGAAAAGTCAGCCAATCCCG -3′ and Ripk3 Wild R: 5′- GCAAGACTAGAGCACACCCTC -3′ for the wild-type allele (375 bp).

### 3MC treatment

C57BL/6N mice (average body weight, 24.0 g), NMRs (average body weight, 26.6 g), and *Ripk3* KO mice were intramuscularly or subcutaneously injected with 3MC (Sigma-Aldrich; 1 mg dissolved in 100 μL corn oil) into the hindlimbs or back skin (60). Animals were observed weekly until tumors >15 mm in diameter developed at the injected sites, at which point the animals were sacrificed humanely using isoflurane anesthesia, and the tumors were used for further analysis. For NMRs, muscle samples (3MC-injected sites and opposite sites as controls) collected after 114 weeks and skin samples collected after 97 weeks were used for histopathological analysis.

To evaluate responses to short exposure to 3MC, 3MC (1 mg dissolved in 100 μL corn oil) was subcutaneously injected into the back skin of C57BL/6N mice or NMRs, and the site was examined at 1 or 3 weeks after injection. Injected sites (100 mm^2^) were collected and used for further analysis. To suppress RIPK3 activity, GSK’872 (31, 45) (SelleckBio; 1 mg/kg body weight dissolved in saline) was intraperitoneally injected three times a week from 1 week before 3MC treatment until the end point of the experiment.

### DMBA/TPA treatment

C57BL/6N mice and NMRs were treated with DMBA (Sigma-Aldrich; 100 μg in 100 μL acetone) on the back skin. One week after DMBA treatment, animals were treated twice a week with TPA (Cayman Chemical; 12.5 μg in 100 μL acetone) until tumor formation was observed (61). Animals were observed daily until tumors >7 mm in diameter developed on the skin, at which point the animals were sacrificed humanely by isoflurane anesthesia, and the tumors were used for further analysis. For NMRs, skin biopsies were performed under isoflurane anesthesia at 55 weeks, and samples were used for histopathological analysis. For histopathological analysis, one individual who had an external wound possibly due to fighting was excluded as previously described (23).

To evaluate responses to short exposure to DMBA, DMBA (100 μg in 100 μL acetone) was administered to the back skin, and skin biopsies were performed after 24 h. To evaluate responses to short exposure to DMBA/TPA, DMBA (100 μg in 100 μL acetone) was administered to the back skin, and 1 week after DMBA treatment, animals were subsequently treated three times with TPA (12.5 μg in 100 μL acetone). Skin biopsies were performed 1 week after starting TPA treatment.

### Hematoxylin and eosin (HE) staining and immunohistochemical analysis

Histological examination was performed at K.I. Stainer, Inc. (Kumamoto, Japan). Briefly, the samples were fixed in 4% paraformaldehyde in phosphate-buffered saline (PBS), embedded in paraffin, and cut into 4 μm sections; HE staining was routinely performed. The antibodies and protocols are listed in Dataset S1. Briefly, for immunostaining, the sections were deparaffinized using xylene and rehydrated with a graded series of ethanol. Antigen retrieval was performed by heat-induced epitope retrieval in citrate buffer or Tris buffer, or by enzymatic retrieval using proteinase K (62). The sections were incubated with 1% bovine serum albumin in Tris-buffered saline with 0.1% NaN_3_ for blocking, and stained with primary antibodies against CD45 (Abcam, ab10558), MPO (DAKO, A0398), IBA1 (FUJIFILM WAKO, 019-19741), CD3 (Nichirei, 413591), Ki67 (Abcam, ab16667), 8-OHdG (Santa Cruz Biotechnology, sc-393871), pH2AX (Cell Signaling Technology [CST], 9718), or HMGB1 (Abcam, ab79823). The sections were incubated with horseradish peroxidase (HRP)-conjugated anti-rabbit, anti-mouse, or anti-rat secondary antibodies (Nichirei) as a secondary antibody. Positive signals were visualized using HistoGreen substrate (Cosmo Bio) for staining of immune cells in the skin (because it is not easy to distinguish diaminobenzidine (DAB)-stained cells from dermal melanin pigments in NMR skin in limited-sized figures) or DAB (Nichirei). For HMGB1, Alexa Fluor 555 anti-rabbit IgG (CST, A21429) secondary antibody was used. Nuclei were counterstained with hematoxylin (for CD45, MPO, IBA1, CD3, Ki67, 8-OHdG, and pH2AX) or Hoechst 33258 (Sigma-Aldrich) for HMGB1. For cleaved caspase-3, 10 μm fresh-frozen sections were fixed with 4% PFA, washed with PBS, and blocked with 5% normal goat serum in 0.3% Triton X-100 (Nacalai Tesque) in PBS. The sections were incubated with primary antibodies against cleaved caspase-3 (CST; 9664; 1:400) in Can Get Signal Solution B (TOYOBO). The sections were stained with Alexa Fluor 555 anti-rabbit IgG (CST; A21429; 1:1000) as a secondary antibody, and nuclei were stained with 1 μg/mL Hoechst 33258 (Sigma-Aldrich).

The images were captured using a BZ-X 710 fluorescence microscope (KEYENCE) and analyzed using a BZ-X image analyzer (KEYENCE).

### TUNEL staining

For TUNEL staining, 4 μm paraffin sections were deparaffinized and rehydrated as described above. The sections were stained using the TUNEL Assay Kit BrdU-Red (Abcam) according to the manufacturer’s instructions. Nuclei were counterstained with Hoechst 33258. The images were captured using a BZ-X 710 fluorescence microscope and analyzed using a BZ-X image analyzer (KEYENCE).

### Morphometric analyses of skin inflammatory responses

Epidermal thickness was quantified by calculating the mean length of skin surface to the epidermal junction by five hand-drawn line segments per field (four fields were analyzed per animal) using ImageJ. Positive cells identified by immunostaining and TUNEL staining were quantified by counting the mean number of cells in each of the four images of one section from more than three animals per experiment, and were normalized to the total number of cells (for Ki67, TUNEL, pH2AX, 8OHdG, cleaved caspase-3, CD45, IBA1, MPO, and CD3) or to the tissue area (for CD45, IBA1, MPO, and CD3). The quantification was performed by three independent investigators including a pathologist (Y. Komohara). Total cells, either Hoechst or hematoxylin-positive nuclei (at least 350 cells per animal) and tissue area (bright field), were measured using a Hybrid Cell Count application (KEYENCE) in a BZ-X image analyzer.

Cytoplasmic HMGB1-positive cells were quantified by counting the mean number of cells in highly magnified sections from more than three animals per experiment, and normalized to the number of total HMGB1-positive cells (at least four images from one section per animal, >100 cells). The quantification was performed by two independent investigators.

### UV irradiation

UV irradiation on the back skin of C57BL/6N mice and NMRs was performed every other day for 12 days with a dose of 1,000 J/m^2^ using a UV lamp (UVP UVM-28; Analytic Jena) for the times indicated in Fig. S7A (25). Prior to irradiation, the back skin of C57BL/6N mice was shaved. At 24 h after final irradiation, the animals were sacrificed humanely using isoflurane anesthesia, and the skin samples were used for further analysis.

### LPS treatment

C57BL/6N mice and NMRs were subcutaneously or intraperitoneally injected with LPS (Sigma-Aldrich; 10 mg/kg body weight dissolved in saline). After 24 h of treatment, the animals were sacrificed humanely using isoflurane anesthesia, and skin or liver samples were used for further analysis.

### RNA isolation and quantification of gene expression

Total RNA was extracted using the RNeasy Plus Mini Kit (Qiagen, for cells) or TRIzol (Thermo Fisher Scientific, for tissues) according to the manufacturer’s protocol. The gDNA Eliminator Spin Column (Qiagen) or TURBO DNA-free™ Kit (Invitrogen) was used for genomic DNA elimination according to the manufacturer’s protocol. Reverse transcription reactions were performed with ReverTra Ace qPCR RT Master Mix (TOYOBO) using 300 ng total RNA as a template. The resulting cDNA was used for reverse transcription polymerase chain reaction (RT-PCR) and quantitative reverse transcription PCR (RT-qPCR). For RT-PCR, 24 cycles (for *actin beta* [*ACTB*]) or 35 cycles (for *MLKL*) of amplification were performed under the following conditions using PrimeSTAR Max DNA Polymerase (Takara): denaturing at 98°C for 10 s, annealing at 55°C for 30 s, and extension at 72°C for 30 s. The DNA fragments were electrophoresed in 2% agarose gels. RT-qPCR analysis was performed using Thunderbird SYBR qPCR Mix (TOYOBO) or PowerUp SYBR Green Master Mix (Thermo Fisher Scientific) on a CFX384 Touch Real-Time PCR Detection System (Bio-Rad) with the primers listed in Dataset S6 (63).

### Cell culture

Primary NMR or mouse skin fibroblasts were obtained from the back skin of 1–2-year-old NMRs or 6–8-week-old C57BL/6N mice (10). The cells were cultured in Dulbecco’s modified Eagle’s medium (Sigma-Aldrich) supplemented with 15% fetal bovine serum (FBS) (for NMR fibroblasts) or 10% FBS (for mouse fibroblasts) (Gibco), 1% penicillin/streptomycin (FUJIFILM WAKO), 2 mM L-glutamine (FUJIFILM WAKO), and 0.1 mM non-essential amino acids (FUJIFILM WAKO) at 32°C in a humidified atmosphere containing 5% O_2_ and 5% CO_2_. We used the fibroblasts within five passages. The medium was replaced every 2 days. For investigation of NMD, NMR fibroblasts were incubated with 5 μg/mL ActD (Sigma-Aldrich) and/or 30 μg/mL CHX (FUJIFILM WAKO) for 4 h. NMR fibroblasts treated with DMSO served as the control. After treatment, total RNA was isolated and used for RT-qPCR as described above.

### Lentiviral overexpression of NMR-MLKL

Because NMR MLKL mRNA was not expressed in NMR skin, the coding sequence of NMR-MLKL was artificially synthesized based on the NCBI sequence information and our genomic sequencing results (XM_021256495.1 and Fig. 4B) (Eurofins Genomics) and inserted into the lentiviral vector pCSII-EF-RFA-hyg (kindly provided by H. Naka-Kaneda). Then, the pCSII-EF-NMR-MLKL plasmid and packaging vectors (pCMV-VSV-G-RSV-Rev and pCAG-HIVgp) (64) were used to transfect 293T cells using a polyethylenimine MAX transfection reagent (CosmoBio) according to the manufacturer’s instructions. The conditioned medium containing viral particles was concentrated by ultracentrifugation and used for viral transduction into NMR SV40ER cells, an NMR skin fibroblast cell line expressing simian virus 40 early region (65). The infected cells were passaged and subjected to propidium iodide (PI) staining as follows.

### Necroptosis assay

Primary mouse or NMR fibroblasts were seeded at 1 × 10^4^ cells/well onto 24-well plates and stimulated with TNF-α (PeproTech; 50 ng/mL), z-VAD-fmk (Abcam; 20 μM), CHX (1 μg/mL), and Nec-1 (Sigma-Aldrich; 20 μM). After 24 h, cells were stained with Hoechst 33342 (DOJINDO; 1 μg/mL) for 10 min at 32°C. Then, the cells were stained with PI (FUJIFILM Wako; 10 μg/mL) for 5 min at 32°C. Images were captured using a BZ-X 710 fluorescence microscope (KEYENCE), and the number of cells positive for PI or Hoechst 33342 was counted (at least 100 cells per treatment) using a BZ-X image analyzer (KEYENCE). PI and Hoechst 33342 double-positive cells were regarded as dead cells.

### Etoposide treatment

NMR fibroblasts were exposed to etoposide at 200 μM for 4 days. Etoposide-containing medium was added to subconfluent fibroblasts. After 2 days, the medium was replaced by freshly prepared etoposide-containing medium for an additional 2 days. Then, the cells were collected for Annexin V/PI analysis and western blotting.

### Flow cytometry analysis for apoptosis detection

The FITC Annexin V Apoptosis Detection Kit (BD Biosciences or BioLegend) was used for the detection of apoptosis. Primary NMR skin fibroblasts were stained according to the manufacturer’s protocols and analyzed on a FACSVerse (BD Biosciences) flow cytometer.

### Phagocytosis assay

NMRs and mice were sacrificed humanely using isoflurane anesthesia, and limbs were isolated. After removing muscles and cartilage tissue, the bones were crushed and suspended in PBS. The cell suspension was filtered through a 70 μm cell strainer (Falcon) and suspended in hypo-osmotic solution to remove red blood cells. The remaining cells after hemolysis were processed into a single cell suspension and cultured in RPMI-1640 (FUJIFILM WAKO) supplemented with 15% FBS, 1% penicillin/streptomycin, 2 mM L-glutamine, 0.1 mM non-essential amino acids, and 20 ng/mL mouse macrophage colony stimulating factor (M-CSF) (BioLegend) for 8 days (66). Dead cells were prepared by 200 J/m^2^ of UVC irradiation to fibroblasts using a UV crosslinker (Analytic Jena). After UV irradiation, cells were cultured for 24 h, and dead cells were collected and stained using pHrodo (Thermo Fisher Scientific) according to the manufacturer’s protocol. The same amounts of pHrodo-labeled dead cells (5 × 10^5^ cells) were co-incubated with NMR or mouse bone marrow macrophage culture. After 2 h, phagocytosis was evaluated by measuring pH-sensitive fluorescence of pHrodo using the BZ-X image analyzer (KEYENCE).

### RNA-seq analysis

Total RNA was extracted from mouse and NMR skin tissues using TRIzol (Thermo Fisher Scientific) and purified using the RNeasy Plus Mini Kit (Qiagen). Any contamination with genomic DNA was removed from total RNA using the RNase-Free DNase Set (Qiagen) according to the manufacturer’s protocol. cDNA libraries were generated from 200 ng total RNA using a TruSeq stranded mRNA library preparation kit (Illumina). The resultant libraries were sequenced on NextSeq550 (Illumina) in single-ended mode. Low-quality bases and the adapters in the sequenced reads were trimmed using Cutadapt (ver.1.14) (67) with Python 2.7.6. The trimmed reads were mapped to either the mouse (mm10) or NMR (HetGla_female_1.0) reference genome, with the UCSC refGene gtf for mouse and the Ensembl HetGla gtf and previously published gff (68) for NMR, using STAR (ver.2.4.1d) (69). For identification of DEGs, the uniquely mapped reads were counted and normalized to calculate fold changes and false discovery rate (FDR) using HTSeq (ver.0.11.2) (70) and edgeR (ver.3.18.1) (71), with the UCSC refGene gtf for mouse and the Ensembl HetGla gtf and previously published gff (68) for NMR. Enrichment of genes in specific cellular functions (GO terms, Reactome, and KEGG pathways) was analyzed using Metascape (29). The gene expression levels were calculated as transcripts per million (TPM) using deepTools (ver.2.1.0) (72), and mapping was visualized using the Integrative Genomics Viewer. The immune enrichment score was analyzed using xCell (28).

### Western blotting

The skin or cell samples were lysed in cell-lysis buffer (125 mM Tris-HCl, pH 6.8, 4% sodium dodecyl sulphate [SDS], and 10% sucrose) and boiled for 10 min. Protein concentration was measured using the BCA Protein Assay Kit (Takara Bio). The protein samples were subjected to SDS-polyacrylamide gel electrophoresis and transferred to polyvinylidene fluoride membranes using the Trans-Blot Turbo Transfer System (Bio-Rad). Membranes were probed with antibodies against MLKL (Abcam, ab184718; 1:1000), pMLKL (Abcam, ab196436; 1:1000), cleaved caspase-3 (CST, 9664; 1:1000), β-actin (CST, 4970; 1:2000), or vinculin (Sigma-Aldrich, V9131; 1:1000). The membranes were incubated with HRP-conjugated anti-rabbit (CST, 7074; 1:1000) or HRP-conjugated anti-mouse (CST, 7076; 1:1000) IgG secondary antibodies and visualized using ECL Prime Western Blotting Detection Reagent (GE Healthcare) and ImageQuant LAS 4000 Mini (FUJIFILM). The experiments were performed in biological duplicates or triplicates.

### Statistical analysis

We used GraphPad Prism (GraphPad ver.8) for statistical analysis. The two groups were analyzed using the two-tailed unpaired *t*-test. For multiple comparisons, the data were analyzed using one-way analysis of variance (ANOVA), followed by Tukey’s multiple comparisons test for multiple comparisons or by Dunnett’s multiple comparisons test. Time to tumor progression was estimated using Kaplan–Meier curves and was statistically analyzed using the log-rank Mantel– Cox test or the Gehan–Breslow–Wilcoxon test. Each data point represents the mean ± standard deviation (SD) derived from at least three animals or biological replicates. *P*-values <0.05 were considered statistically significant.

## Acknowledgments

We thank Drs. H. Niwa, K. Yamagata, K. Seino, and H. Wada for scientific discussions and administrative support; Ms. Takana Motoyoshi (K.I. Stainer, Kumamoto, Japan) for technical assistance for immunohistochemistry; M. Kobe, Y. Tanabe, and Y. Fujimura for help with animal maintenance; and all members of the K.M. Laboratory for technical assistance and scientific discussions. We thank H. Miyoshi for lentiviral vectors and H. Naka-Kaneda for the pCSII-EF-RfA-TK-hyg vector. We thank Jane Doe of the Liaison Laboratory Research Promotion Center for technical support. This work was supported in part by AMED under Grant Numbers JP21bm0704040 and JP21gm5010001; Grants-in-Aid for Scientific Research from the Japanese Society for the Promotion of Science from the Ministry of Education, Culture, Sports, Science and Technology (MEXT) to K.M., Y. Kawamura, and K.O.; JSPS KAKENHI Grant Number JP 16H06276 (AdAMS) to K.O.; and the Tenure-Track Grant of Kumamoto University to Y. Kawamura. K.M. was supported by the Takeda Science Foundation, KOSÉ Cosmetology Research Foundation, Kanzawa Medical Research Foundation, Nakatomi Foundation, Naito Foundation, Foundation for Promotion of Cancer Research, Kato Memorial Bioscience Foundation, MSD Life Science Foundation, Inamori Foundation, and Frontier Salon Foundation. The ASHBi is supported by the World Premier International Research Center Initiative (WPI), MEXT, Japan.

## References

1. B. P. Lee, M. Smith, R. Buffenstein, L. W. Harries, Negligible senescence in naked mole rats may be a consequence of well-maintained splicing regulation. GeroScience 42, 633–651 (2020).

2. J. G. Ruby, M. Smith, R. Buffenstein, Naked mole-rat mortality rates defy Gompertzian laws by not increasing with age. Elife 7, 1–18 (2018).

3. R. Buffenstein, Negligible senescence in the longest living rodent, the naked mole-rat: insights from a successfully aging species. J. Comp. Physiol. B. 178, 439–45 (2008).

4. M. A. Delaney, L. Nagy, M. J. Kinsel, P. M. Treuting, Spontaneous Histologic Lesions of the Adult Naked Mole Rat (*Heterocephalus glaber*). Vet. Pathol. 50, 607–621 (2013).

5. M. A. Delaney, et al., Initial Case Reports of Cancer in Naked Mole-rats (*Heterocephalus glaber*). Vet. Pathol. 53, 691–6 (2016).

6. K. R. Taylor, N. A. Milone, C. E. Rodriguez, Four cases of spontaneous neoplasia in the naked mole-rat (*Heterocephalus glaber*), a putative cancer-resistant species. Journals Gerontol. – Ser. A Biol. Sci. Med. Sci. 72, 38–43 (2017).

7. J. E. Cole, J. C. Steeil, S. J. Sarro, K. L. Kerns, A. Cartoceti, Chordoma of the sacrum of an adult naked mole-rat. J. Vet. Diagnostic Investig. 32, 132–135 (2020).

8. A. Seluanov, et al., Hypersensitivity to contact inhibition provides a clue to cancer resistance of naked mole-rat. Proc. Natl. Acad. Sci. U. S. A. 106, 19352–19357 (2009).

9. X. Tian, et al., High-molecular-mass hyaluronan mediates the cancer resistance of the naked mole rat. Nature 499, 346–349 (2013).

10. S. Miyawaki, et al., Tumour resistance in induced pluripotent stem cells derived from naked mole-rats. Nat. Commun. 7, 11471 (2016).

11. S. Liang, J. Mele, Y. Wu, R. Buffenstein, P. J. Hornsby, Resistance to experimental tumorigenesis in cells of a long-lived mammal, the naked mole-rat (Heterocephalus glaber). Aging Cell 9, 626–635 (2010).

12. F. Hadi, et al., Transformation of naked mole-rat cells. Nature 583, E1–E7 (2020).

13. J. DiGiovanni, Multistage carcinogenesis in mouse skin. Pharmacol. Ther. 54, 63–128 (1992).

14. D. Hanahan, R. A. Weinberg, Hallmarks of cancer: The next generation. Cell 144, 646–674 (2011).

15. E. Hoste, et al., Innate sensing of microbial products promotes wound-induced skin cancer. Nat. Commun. 6, 5932 (2015).

16. T. Bald, et al., Ultraviolet-radiation-induced inflammation promotes angiotropism and metastasis in melanoma. Nature 507, 109–113 (2014).

17. D. Mittal, et al., TLR4-mediated skin carcinogenesis is dependent on immune and radioresistant cells. EMBO J. 29, 2242–2252 (2010).

18. H. Oshima, M. Oshima, The inflammatory network in the gastrointestinal tumor microenvironment: Lessons from mouse models. J. Gastroenterol. 47, 97–106 (2012).

19. M. E. Choi, D. R. Price, S. W. Ryter, A. M. K. Choi, Necroptosis: a crucial pathogenic mediator of human disease. JCI Insight 4 (2019).

20. T. H. Schreiber, E. R. Podack, A critical analysis of the tumour immunosurveillance controversy for 3-MCA-induced sarcomas. Br. J. Cancer 101, 381–386 (2009).

21. M. B. Shimkin, G. B. Mider, Induction Of Tumors In Guinea Pigs With Subcutaneously Injected Methylcholanthrene. JNCI J. Natl. Cancer Inst. 1, 707–725 (1941).

22. I. Manov, et al., Pronounced cancer resistance in a subterranean rodent, the blind mole-rat, Spalax: in vivo and in vitro evidence. BMC Biol. 11, 91 (2013).

23. E. L. Abel, J. M. Angel, K. Kiguchi, J. DiGiovanni, Multi-stage chemical carcinogenesis in mouse skin: Fundamentals and applications. Nat. Protoc. 4, 1350–1362 (2009).

24. J. Zhang, et al., Fibroblast-specific protein 1/S100A4-positive cells prevent carcinoma through collagen production and encapsulation of carcinogens. Cancer Res. 73, 2770–2781 (2013).

25. K. W. Chung, et al., Molecular insights into SIRT1 protection against UVB-induced skin fibroblast senescence by suppression of oxidative stress and p53 acetylation. Journals Gerontol. – Ser. A Biol. Sci. Med. Sci. 70, 959–968 (2015).

26. H. G. Hilton, et al., Single-cell transcriptomics of the naked mole-rat reveals unexpected features of mammalian immunity. PLOS Biol. 17, e3000528 (2019).

27. J. A. Ramilowski, et al., A draft network of ligand–receptor-mediated multicellular signalling in human. Nat. Commun. 6, 7866 (2015).

28. D. Aran, Z. Hu, A. J. Butte, xCell: Digitally portraying the tissue cellular heterogeneity landscape. Genome Biol. 18, 1–14 (2017).

29. Y. Zhou, et al., Metascape provides a biologist-oriented resource for the analysis of systems-level datasets. Nat. Commun. 10, 1523 (2019).

30. A. Linkermann, D. R. Green, Necroptosis. N. Engl. J. Med. 370, 455–465 (2014).

31. S. B. Lee, et al., The AMPK–Parkin axis negatively regulates necroptosis and tumorigenesis by inhibiting the necrosome. Nat. Cell Biol. 21, 940–951 (2019).

32. L. Seifert, et al., The necrosome promotes pancreatic oncogenesis via CXCL1 and Mincle-induced immune suppression. Nature 532, 245–249 (2016).

33. Y. S. Cho, et al., Phosphorylation-Driven Assembly of the RIP1-RIP3 Complex Regulates Programmed Necrosis and Virus-Induced Inflammation. Cell 137, 1112–1123 (2009).

34. S. He, et al., Receptor Interacting Protein Kinase-3 Determines Cellular Necrotic Response to TNF-α. Cell 137, 1100–1111 (2009).

35. J. A. Rickard, et al., RIPK1 regulates RIPK3-MLKL-driven systemic inflammation and emergency hematopoiesis. Cell 157, 1175–1188 (2014).

36. Y. Dondelinger, P. Hulpiau, Y. Saeys, M. J. M. Bertrand, P. Vandenabeele, An evolutionary perspective on the necroptotic pathway. Trends Cell Biol. 26, 721–732 (2016).

37. F. Lejeune, L. E. Maquat, Mechanistic links between nonsense-mediated mRNA decay and pre-mRNA splicing in mammalian cells. Curr. Opin. Cell Biol. 17, 309–315 (2005).

38. J. M. Hildebrand, et al., Activation of the pseudokinase MLKL unleashes the four-helix bundle domain to induce membrane localization and necroptotic cell death. Proc. Natl. Acad. Sci. U. S. A. 111, 15072–15077 (2014).

39. M. C. Tanzer, et al., Evolutionary divergence of the necroptosis effector MLKL. Cell Death Differ. 23, 1185–1197 (2016).

40. X. Chen, et al., Translocation of mixed lineage kinase domain-like protein to plasma membrane leads to necrotic cell death. Cell Res. 24, 105–121 (2014).

41. A. Degterev, W. Zhou, J. L. Maki, J. Yuan, “Assays for Necroptosis and Activity of RIP Kinases” in Regulated Cell Death Part B, A. Ashkenazi, J. A. Wells, J. B. T.-M. in E. Yuan, Eds. (Academic Press, 2014), pp. 1–33.

42. T. H. Lee, et al., The Death Domain Kinase RIP1 Is Essential for Tumor Necrosis Factor Alpha Signaling to p38 Mitogen-Activated Protein Kinase. Mol. Cell. Biol. 23, 8377–8385 (2003).

43. Y. Dondelinger, et al., RIPK3 contributes to TNFR1-mediated RIPK1 kinase-dependent apoptosis in conditions of cIAP1/2 depletion or TAK1 kinase inhibition. Cell Death Differ. 20, 1381–1392 (2013).

44. M. Pasparakis, P. Vandenabeele, Necroptosis and its role in inflammation. Nature 517, 311–320 (2015).

45. P. Mandal, et al., RIP3 Induces Apoptosis Independent of Pronecrotic Kinase Activity. Mol. Cell 56, 481–495 (2014).

46. R. Altwasser, et al., The transcriptome landscape of the carcinogenic treatment response in the blind mole rat: Insights into cancer resistance mechanisms. BMC Genomics 20, 1–15 (2019).

47. V. Domankevich, H. Eddini, A. Odeh, I. Shams, Resistance to DNA damage and enhanced DNA repair capacity in the hypoxia-tolerant blind mole rat, Spalax. J. Exp. Biol. 221 (2018).

48. X. Tian, et al., SIRT6 Is Responsible for More Efficient DNA Double-Strand Break Repair in Long-Lived Species. Cell 177, 622–638.e22 (2019).

49. A. Evdokimov, et al., Naked mole rat cells display more efficient excision repair than mouse cells. Aging (Albany. NY). 10, 1454–1473 (2018).

50. S. Dadi, et al., Cancer Immunosurveillance by Tissue-Resident Innate Lymphoid Cells and Innate-like T Cells. Cell 164, 365–377 (2016).

51. L. A. Kalekar, et al., Regulatory T cells in skin are uniquely poised to suppress profibrotic immune responses. Sci. Immunol. 4, 1–14 (2019).

52. J. Xie, et al., Dampened STING-Dependent Interferon Activation in Bats. Cell Host Microbe 23, 297–301.e4 (2018).

53. D. Jebb, et al., Six reference-quality genomes reveal evolution of bat adaptations. Nature 583, 578–584 (2020).

54. T. J. Park, et al., Fructose-driven glycolysis supports anoxia resistance in the naked mole-rat. Science. 356, 307–311 (2017).

55. J. Artwohl, et al., Extreme susceptibility of african naked mole rats (*Heterocephalus glaber*) to experimental infection with herpes simplex virus type 1. Comp. Med. 59, 83–90 (2009).

56. Y. H. Edrey, et al., Amyloid beta and the longest-lived rodent: the naked mole-rat as a model for natural protection from Alzheimer’s disease. Neurobiol. Aging 34, 2352–2360 (2013).

57. S. He, X. Wang, RIP kinases as modulators of inflammation and immunity. Nat. Immunol. 19, 912–922 (2018).

58. S. Martens, J. Bridelance, R. Roelandt, P. Vandenabeele, N. Takahashi, MLKL in cancer: more than a necroptosis regulator. Cell Death Differ. 28, 1757–1772 (2021).

59. A. Avgustinova, et al., Loss of G9a preserves mutation patterns but increases chromatin accessibility, genomic instability and aggressiveness in skin tumours. Nat. Cell Biol. 20, 1400–1409 (2018).

60. I. García-Cao, et al., “Super p53” mice exhibit enhanced DNA damage response, are tumor resistant and age normally. EMBO J. 21, 6225–6235 (2002).

61. K. Yamakoshi, et al., Real-time in vivo imaging of p16Ink4a reveals cross talk with p53. J. Cell Biol. 186, 393–407 (2009).

62. T. Nakagawa, et al., Optimum immunohistochemical procedures for analysis of macrophages in human and mouse formalin fixed paraffin-embedded tissue samples. J. Clin. Exp. Hematop. 57, 31–36 (2017).

63. J. Cheng, et al., Comparative study of macrophages in naked mole rats and ICR mice. Oncotarget 8, 96924–96934 (2017).

64. H. Miyoshi, U. Blömer, M. Takahashi, F. H. Gage, I. M. Verma, Development of a Self-Inactivating Lentivirus Vector. J. Virol. 72, 8150–8157 (1998).

65. S. Yamaguchi, et al., Characterization of an active LINE-1 in the naked mole-rat genome. Sci. Rep. 11, 1–8 (2021).

66. H. Wada, et al., Flow cytometric identification and cell-line establishment of macrophages in naked mole-rats. Sci. Rep. 9, 1–12 (2019).

67. M. Martin, Cutadapt removes adapter sequences from high-throughput sequencing reads. EMBnet.journal 17, 10 (2011).

68. M. Bens, et al., Naked mole-rat transcriptome signatures of socially suppressed sexual maturation and links of reproduction to aging. BMC Biol. 16, 77 (2018).

69. A. Dobin, et al., STAR: Ultrafast universal RNA-seq aligner. Bioinformatics 29, 15–21 (2013).

70. S. Anders, P. T. Pyl, W. Huber, HTSeq-A Python framework to work with high-throughput sequencing data. Bioinformatics 31, 166–169 (2015).

71. M. D. Robinson, D. J. McCarthy, G. K. Smyth, edgeR: A Bioconductor package for differential expression analysis of digital gene expression data. Bioinformatics 26, 139–140 (2009).

72. F. Ramírez, F. Dündar, S. Diehl, B. A. Grüning, T. Manke, DeepTools: A flexible platform for exploring deep-sequencing data. Nucleic Acids Res. 42, 187–191 (2014).

