## Supplementary Material for "Resistance to chemical carcinogenesis induction via a dampened inflammatory response in naked mole-rats"

#### Supplementary Material for
### Resistance to chemical carcinogenesis induction via a dampened inflammatory response in naked mole-rats

Kaori Oka[a,b,1], Shusuke Fujioka[a,b,1], Yoshimi Kawamura[a,b,1], Yoshihiro Komohara[c], Takeshi Chujo[d], Koki Sekiguchi[a], Yuki Yamamura[a], Yuki Oiwa[a,b], Natsuko Omamiuda-Ishikawa[a], Shohei Komaki[e], Yoichi Sutoh[e], Satoko Sakurai[f], Kazuhito Tomizawa[d,g], Hidemasa Bono[h], Atsushi Shimizu[e,i], Kimi Araki[g,j], Takuya Yamamoto[f,k,l,m], Yasuhiro Yamada[m,n], Hiroyuki Oshiumi[o], and Kyoko Miura[a,b,g*]

*Corresponding author: Kyoko Miura


**This PDF file includes:**

Figures S1 to S18
Datasets S1 to S6

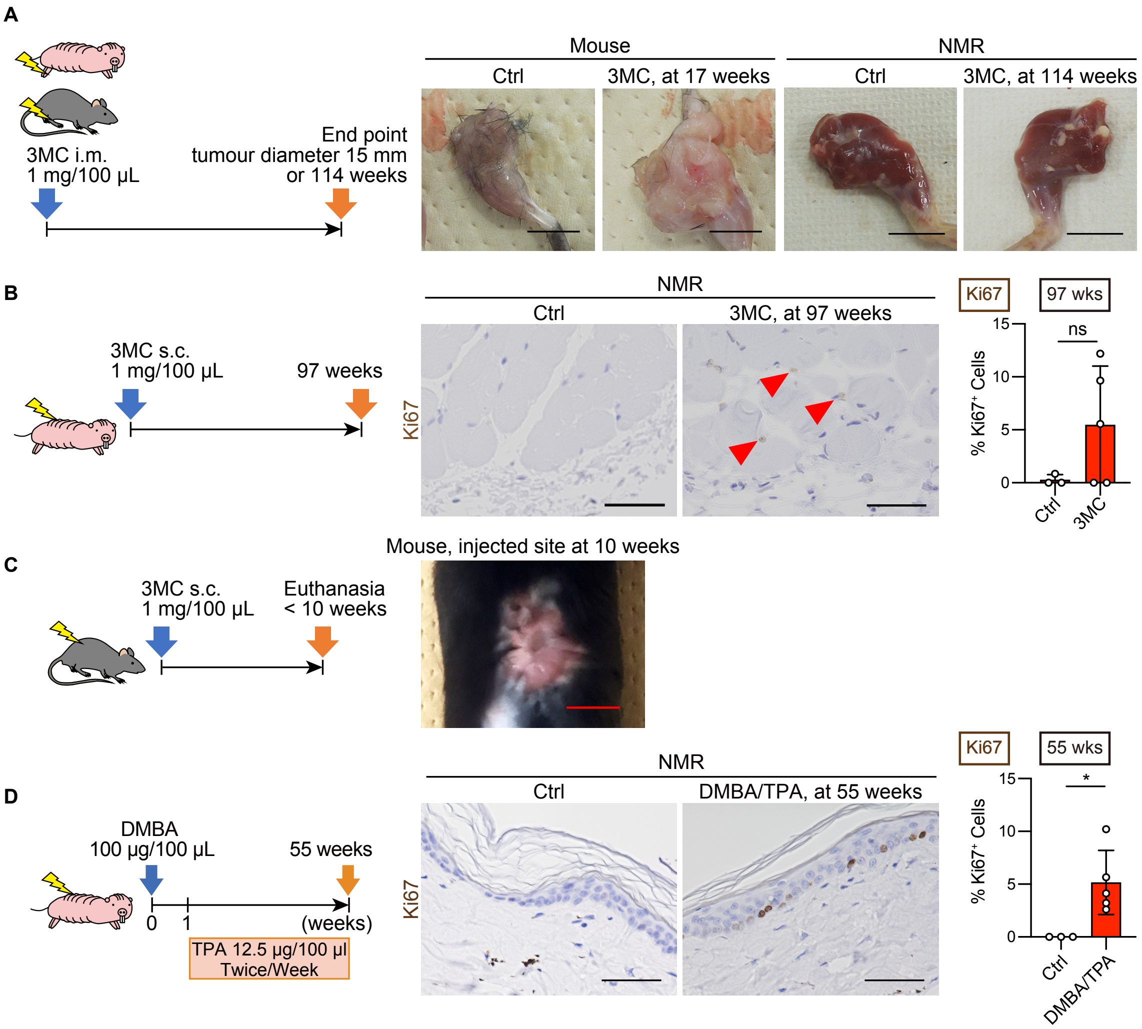


**Fig. S1. Response of mouse and naked mole-rat (NMR) skin to two types of chemical carcinogenesis induction.**

**A**, Gross appearance of the mouse limbs at 17 weeks and NMR limbs at 114 weeks after intramuscular (i.m.) injection of 1 mg 3-methylcholanthrene (3MC). The contralateral legs that were not injected with 3MC served as the control (Ctrl). Scale bar: 1 cm. **B**, Immunohistochemical staining and quantification of Ki67 (brown)-positive cells in NMR skin at 97 weeks after subcutaneous (s.c.) injection of 1mg 3MC. Red arrowheads show positive cells. Scale bar: 50 µm. **C**, Gross appearance of mouse back skin at 10 weeks after s.c. injection of 1 mg 3MC. Scale bar: 1 cm. We tested five animals. **D**, Immunohistochemical staining and quantification of Ki67 (brown)-positive cells in NMR skin at 55 weeks after starting 7,12-dimethylbenz[a]anthracene (DMBA)/12-O-tetradecanoylphorbol-13-acetate (TPA) treatment. Scale bar: 50 µm. For quantification in **B** and **D**, data are presented as the mean ± SD of $n$ = 3 (for control) or $n$ = 5 (for 3MC and DMBA/TPA) animals. Unpaired $t$-test versus untreated control (Ctrl).

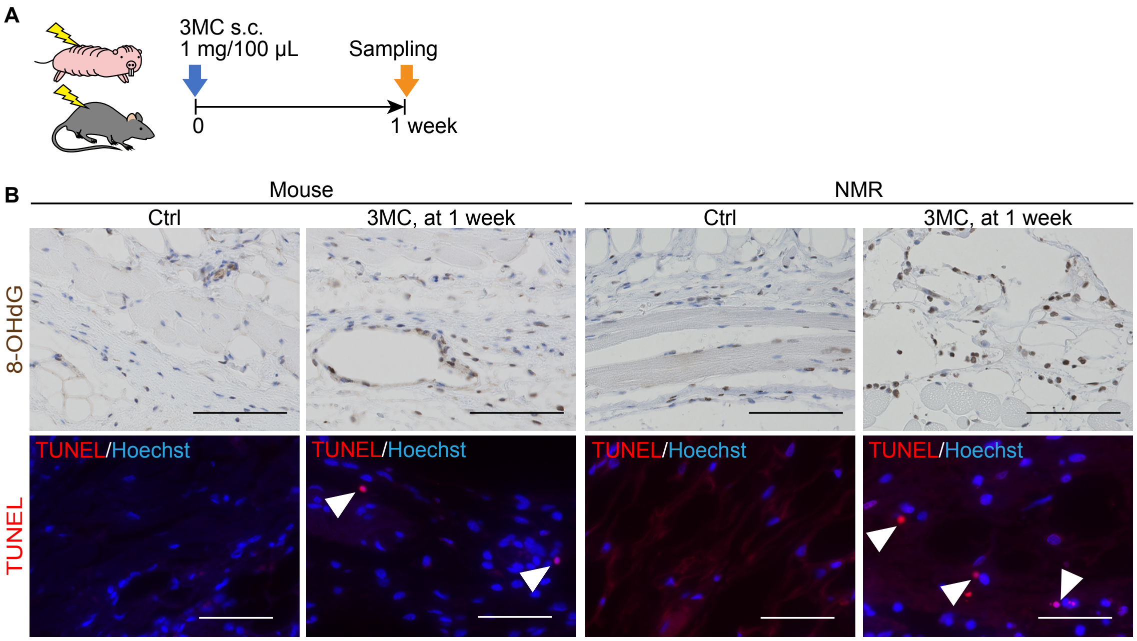


**Fig. S2. Tissue damage responses of mouse and NMR skin to 3MC treatment at 1 week.**
**A**, Schematic diagram for investigating short-term responses to 3MC after subcutaneous (s.c.) injection into the back skin. **B**, Immunohistochemical staining for 8-hydroxy-2′-deoxyguanosine (8-OHdG, brown)- and TUNEL (red) staining of the skin of mice and NMRs at 1 week after s.c. injection of 3MC. Scale bars: 100 μm (8-OHdG) and 50 μm (TUNEL). White arrowheads indicate TUNEL-positive cells.

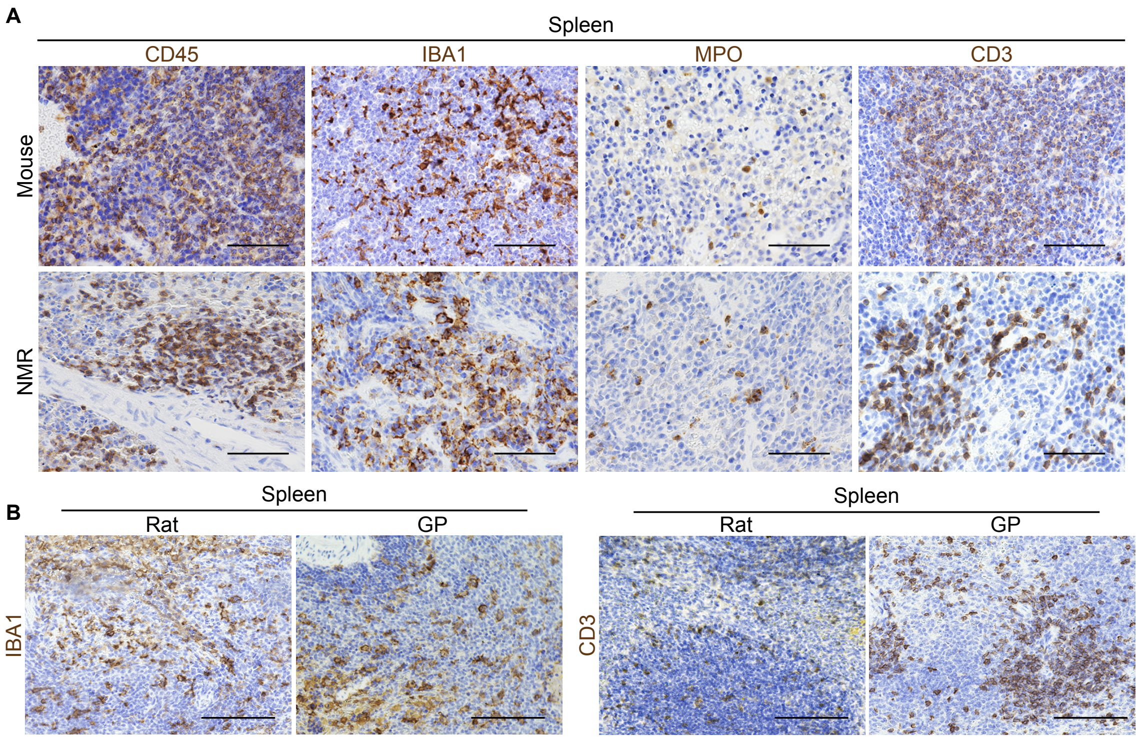


**Fig. S3. Validation of antibodies for immune cell markers by immunohistochemical staining in the mouse, NMR, rat and guinea pig (GP) spleen.**

**A**, Immunohistochemical staining (brown) for CD45, IBA1, myeloperoxidase (MPO), and CD3 in mouse and NMR spleens. Scale bar: 50 μm. **B**, Immunohistochemical staining (brown) for IBA1 and CD3 in rat and GP spleens. Scale bar: 100 μm.

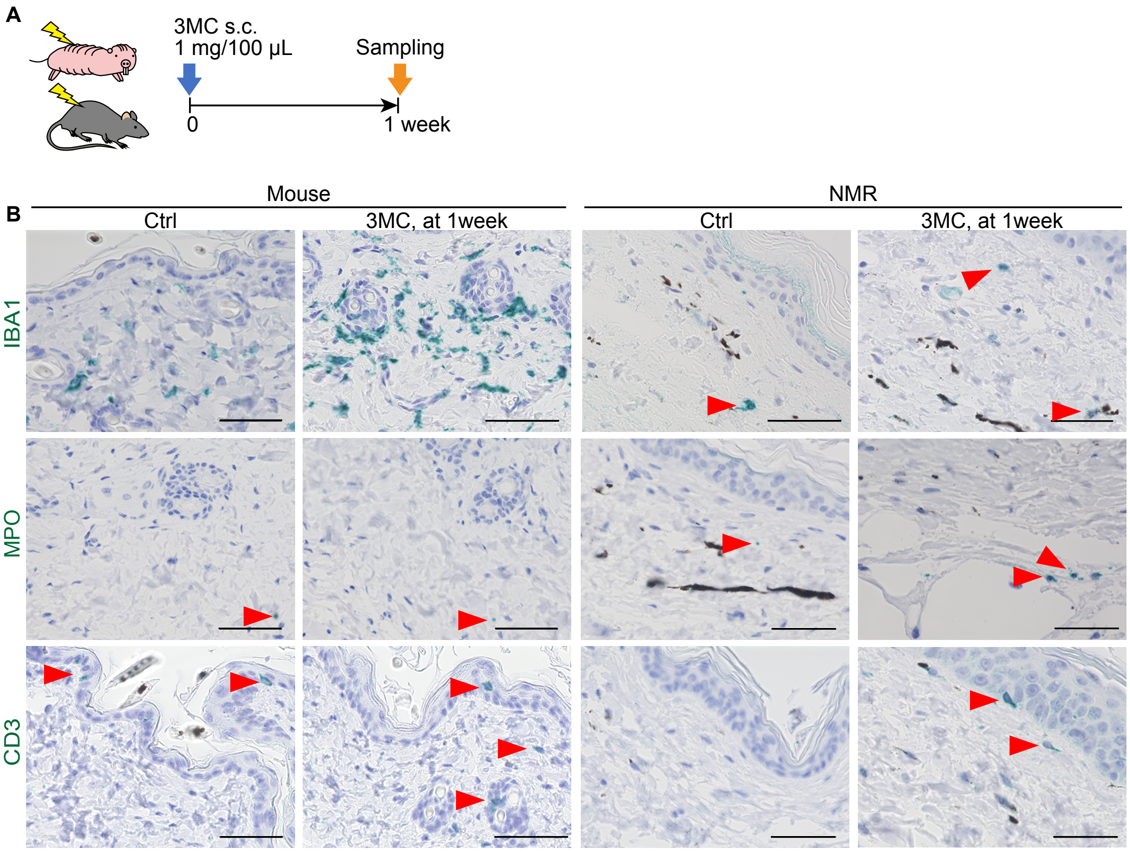


**Fig. S4. Immune cell responses of mouse and NMR skin to 3MC treatment at 1 week.**
**A**, Schematic diagram for investigating short-term responses to 3MC after subcutaneous (s.c.) injection into the back skin. **B**, Immunohistochemical detection (green) of IBA1-, MPO-, and CD3-positive cells in the skin of mice and NMRs at 1 week after 3MC injection. Scale bar: 50 μm. Red arrowheads show positive cells that are hard to see.

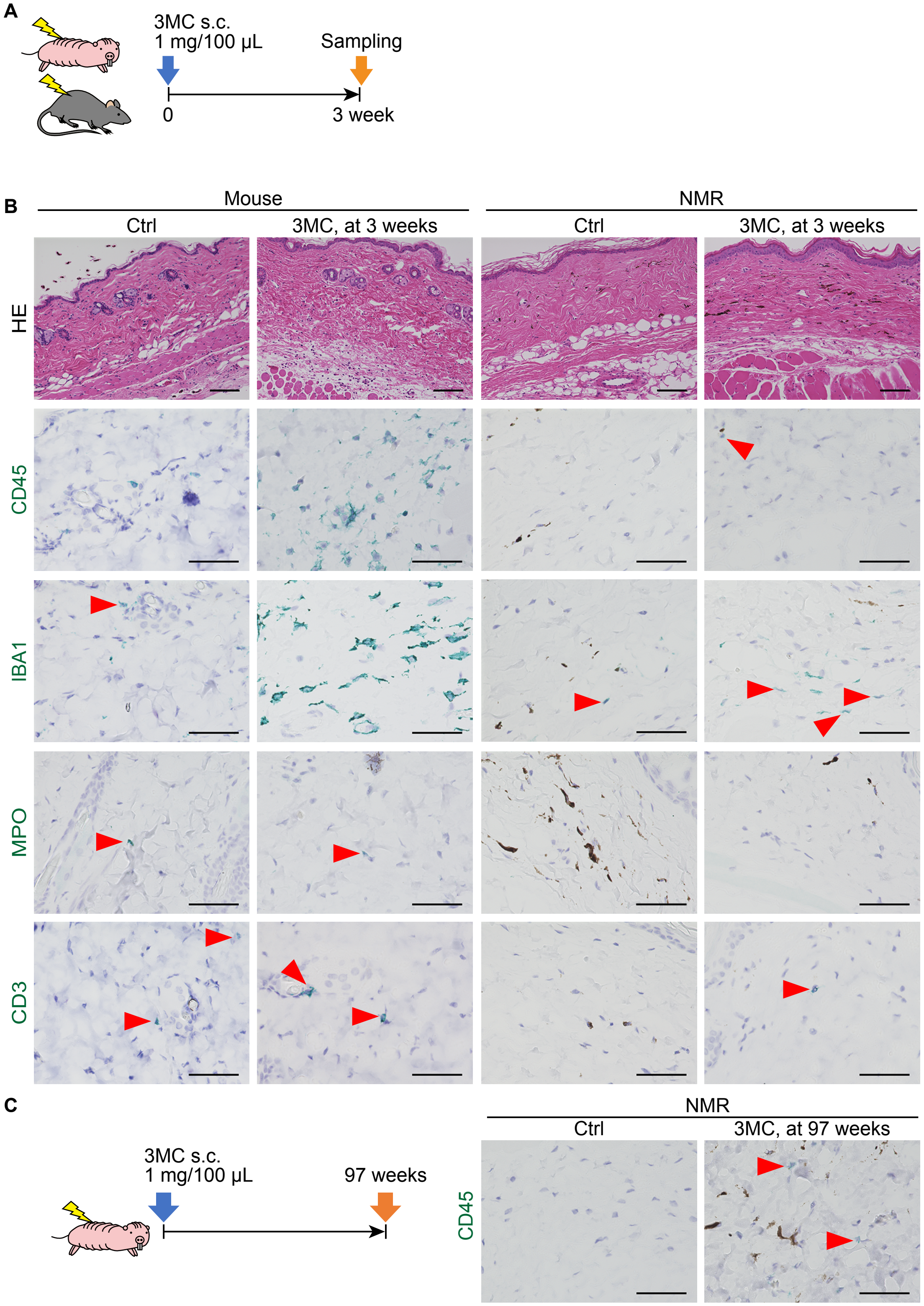


**Fig. S5. Immune cell responses of mouse and NMR skin to 3MC treatment at 3 and 97 weeks.**

**A**, Schematic diagram for investigating short-term responses to 3MC after subcutaneous (s.c.) injection into the back skin. **B**, Hematoxylin and eosin (HE) staining and immunohistochemical detection (green) of CD45-, IBA1-, MPO-, and CD3-positive cells in the skin of mice and NMRs at

3 weeks after s.c. injection of 3MC. Scale bars: 100 μm (HE) and 50 μm (others). Red arrowheads show positive cells that are hard to see. **C**, Immunohistochemical detection of CD45 (green)-positive cells in NMR skin at 97 weeks after s.c. injection of 3MC. Scale bar: 50 μm. Red arrowheads show positive cells.

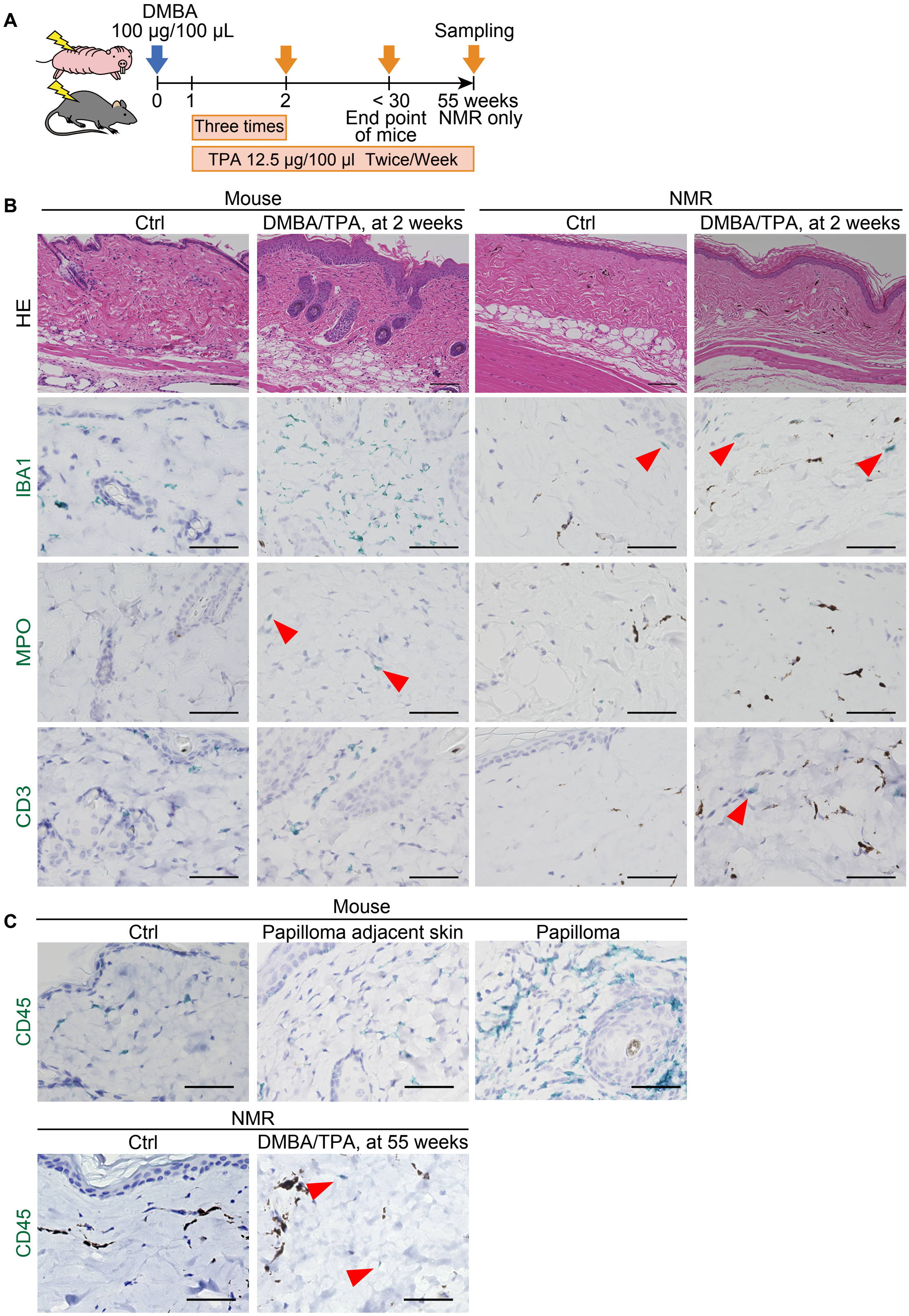


**Fig. S6. Immune cell responses of mouse and NMR skin to DMBA/TPA treatment.**
**A**, Schematic diagram for investigating responses after exposure to DMBA/TPA in the skin. **B**, HE staining and immunohistochemical detection (green) of IBA1-, MPO-, and CD3-positive cells in

the skin of mice and NMRs at 2 weeks after exposure to DMBA/TPA. Scale bars: 100 μm (HE) and 50 μm (others). Red arrowheads show positive cells that are hard to see. **C**, Immunohistochemical detection of CD45 (green)-positive cells in mouse skin sections at the end point and in NMRs at 55 weeks after exposure to DMBA/TPA. “Papilloma adjacent skin” is the no-papilloma region from DMBA/TPA-treated mouse skin. Scale bar: 50 μm. Red arrowheads show positive cells in NMRs.

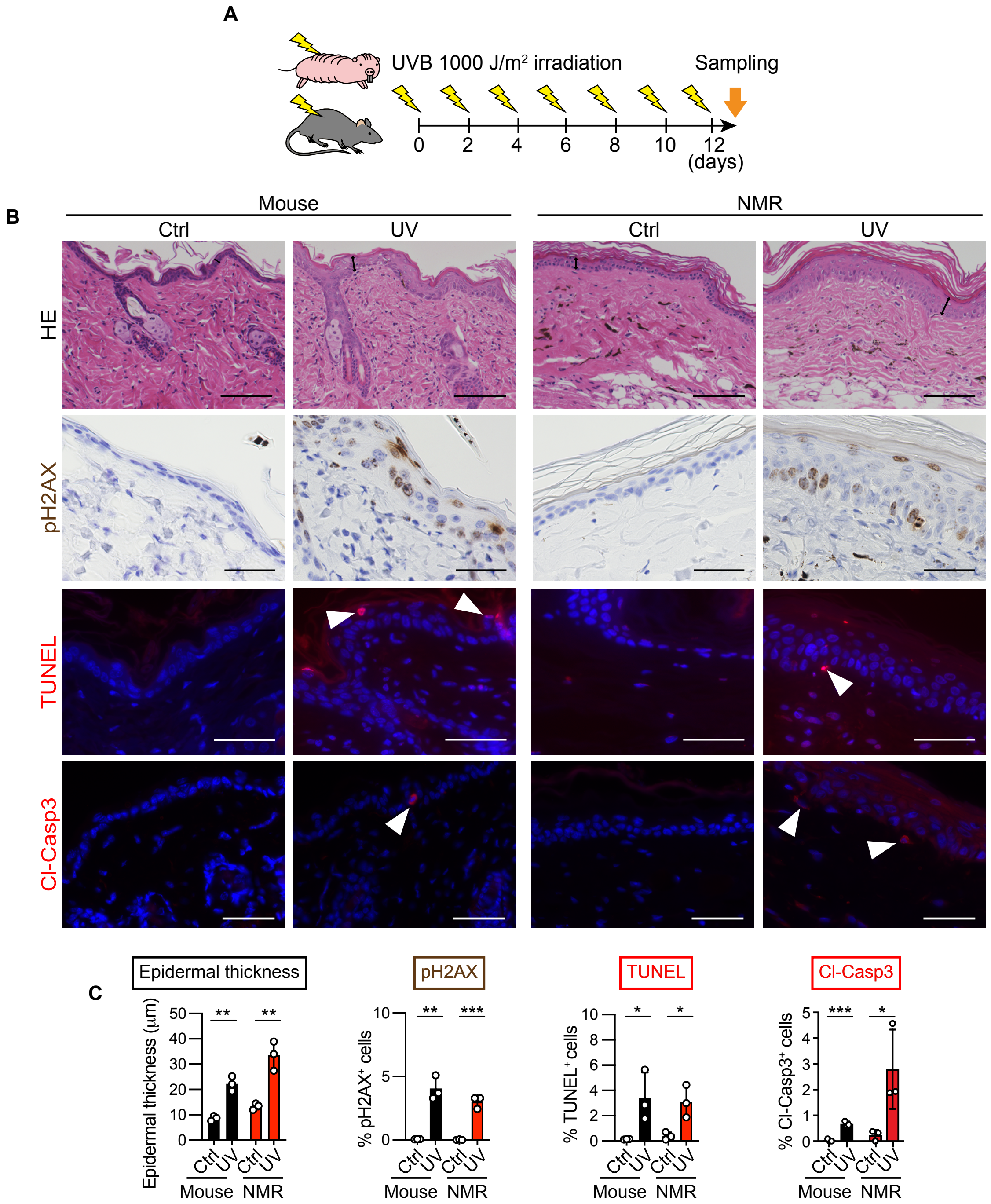


**Fig. S7. Tissue damage responses of mouse and NMR skin to UV irradiation.**
**A**, Schematic diagram for investigating responses to UVB irradiation. **B**, HE staining and immunohistochemical staining for phospho-histone H2A.X (pH2AX, brown), TUNEL (red), and cleaved caspase-3 (red) in the skin of mice and NMRs after UVB irradiation. Scale bars: 100 μm (HE) and 50 μm (others). Double-headed arrows show epidermal thickness, and white arrowheads show positive cells. **C**, Quantification of epidermal thickness from HE staining images and pH2AX-, TUNEL-, and cleaved caspase-3-positive cells per total cells after UVB irradiation. Data are presented as the mean ± SD of $n$ = 3 animals. Unpaired $t$-test versus untreated control (Ctrl).

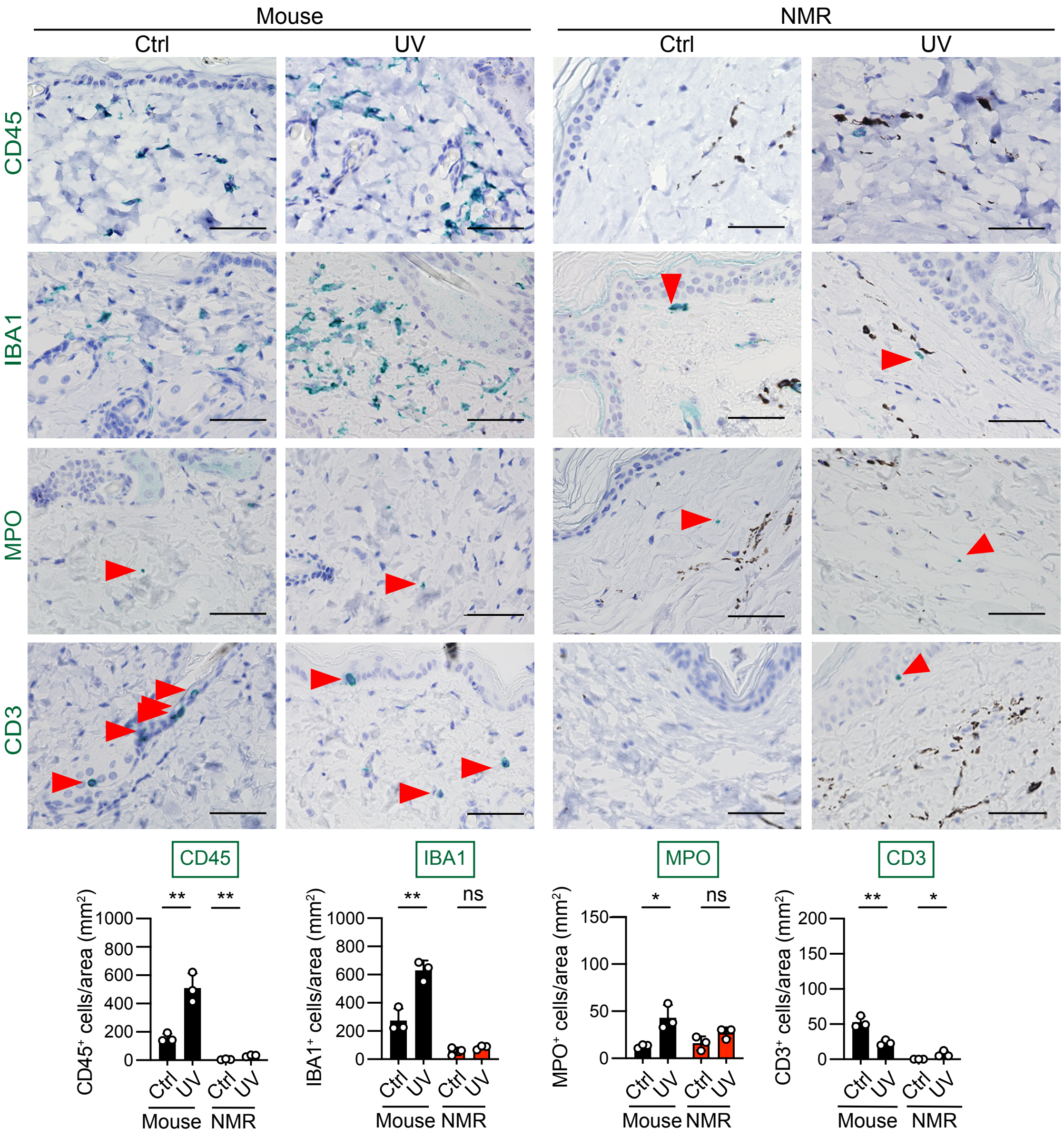


**Fig. S8. Immune cell responses of mouse and NMR skin after UV irradiation.**
Immunohistochemical staining (green) for CD45, IBA1, MPO, and CD3 and quantification of positive cells per area in the skin after UVB irradiation. Scale bar: 50 μm. Red arrowheads show positive cells that are hard to see. Data are presented as the mean ± SD of $n$ = 3 animals. Unpaired $t$-test versus untreated control (Ctrl).

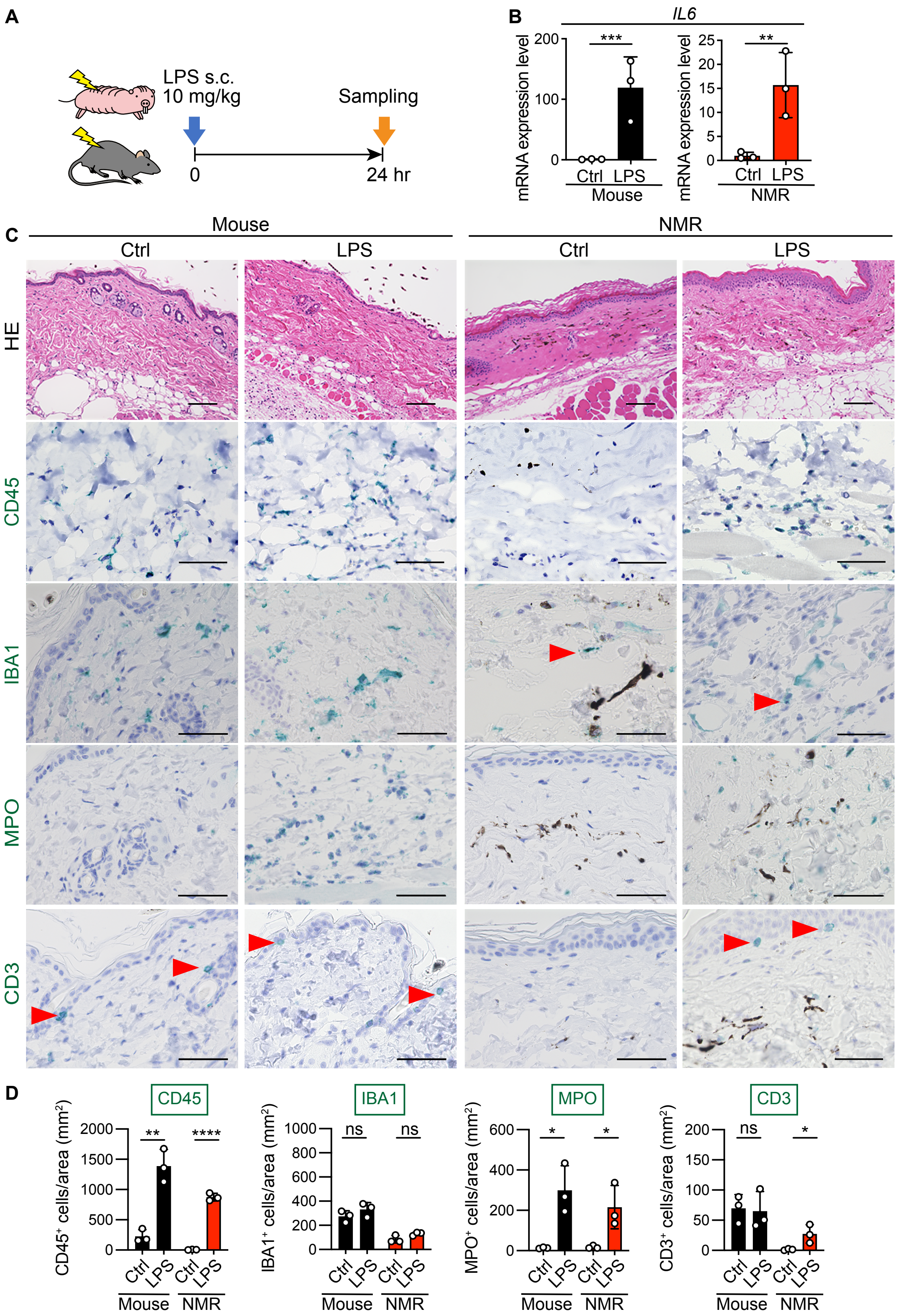


**Fig. S9. Responses of mouse and NMR skin to lipopolysaccharide (LPS) treatment.**
**A**, Schematic diagram for investigating responses to LPS subcutaneous (s.c.) injection. **B**, Relative interleukin-6 (*IL6*) mRNA levels in the skin of mice and NMRs after LPS injection

quantified by RT-qPCR and normalized to actin beta (*ACTB*) mRNA. The values shown are the average fold change relative to that of the untreated control. Primers are listed in Dataset S6. **C**, HE staining and immunohistochemical staining (green) for CD45, IBA1, MPO, and CD3 in the skin after s.c. injection of LPS. Scale bars: 100 μm (HE) and 50 μm (others). Red arrowheads show positive cells that are hard to see. **D**, Quantification of CD45-, IBA1-, MPO-, and CD3-positive cells per area at 24 h after LPS injection. For B and D, data are presented as the mean ± SD of *n* = 3 animals. Unpaired *t*-test versus untreated control (Ctrl).

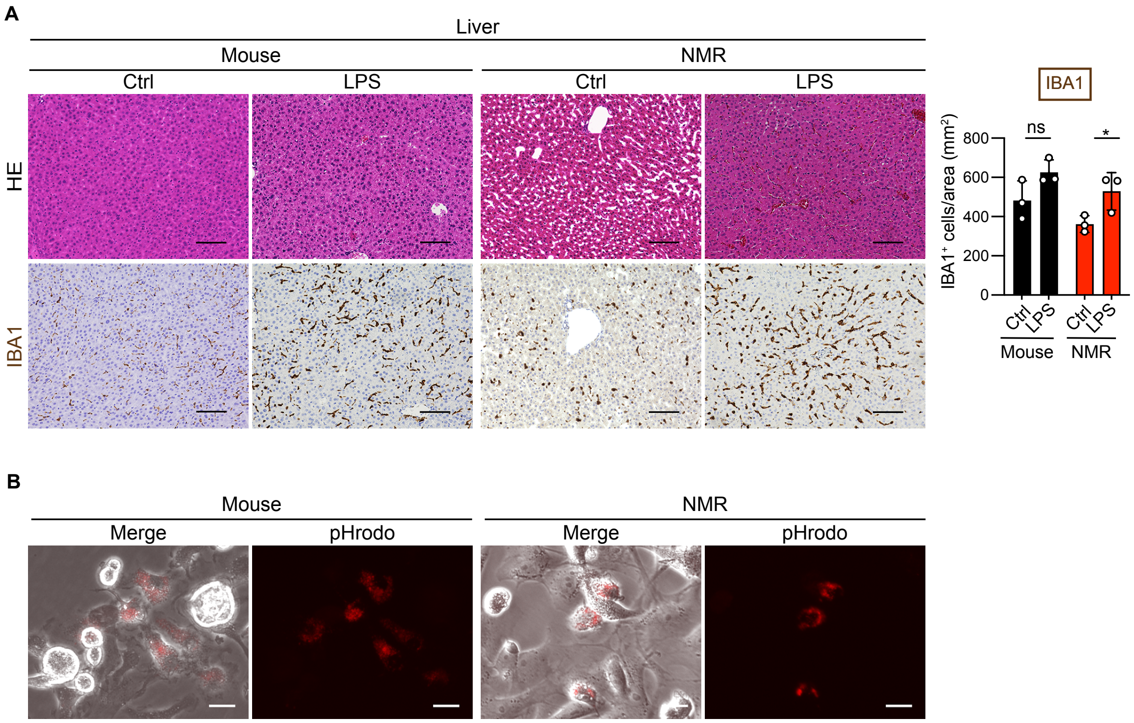


**Fig. S10. Response of livers to LPS and phagocytotic activity of bone marrow-derived macrophages in mice and NMRs.**

**A**, HE staining, immunohistochemical staining, and quantification of IBA1 (brown)-positive cells in the livers of mice and NMRs at 24 h after intraperitoneal LPS injection. Scale bar: 100 μm. Data are presented as the mean ± SD of $n$ = 3 animals. Unpaired $t$-test versus untreated control (Ctrl). **B**, Phagocytotic activity analysis of mouse and NMR bone marrow-derived macrophages. Scale bar: 20 μm. Only phagocytosed pHrodo-labeled dead cells show red fluorescence.

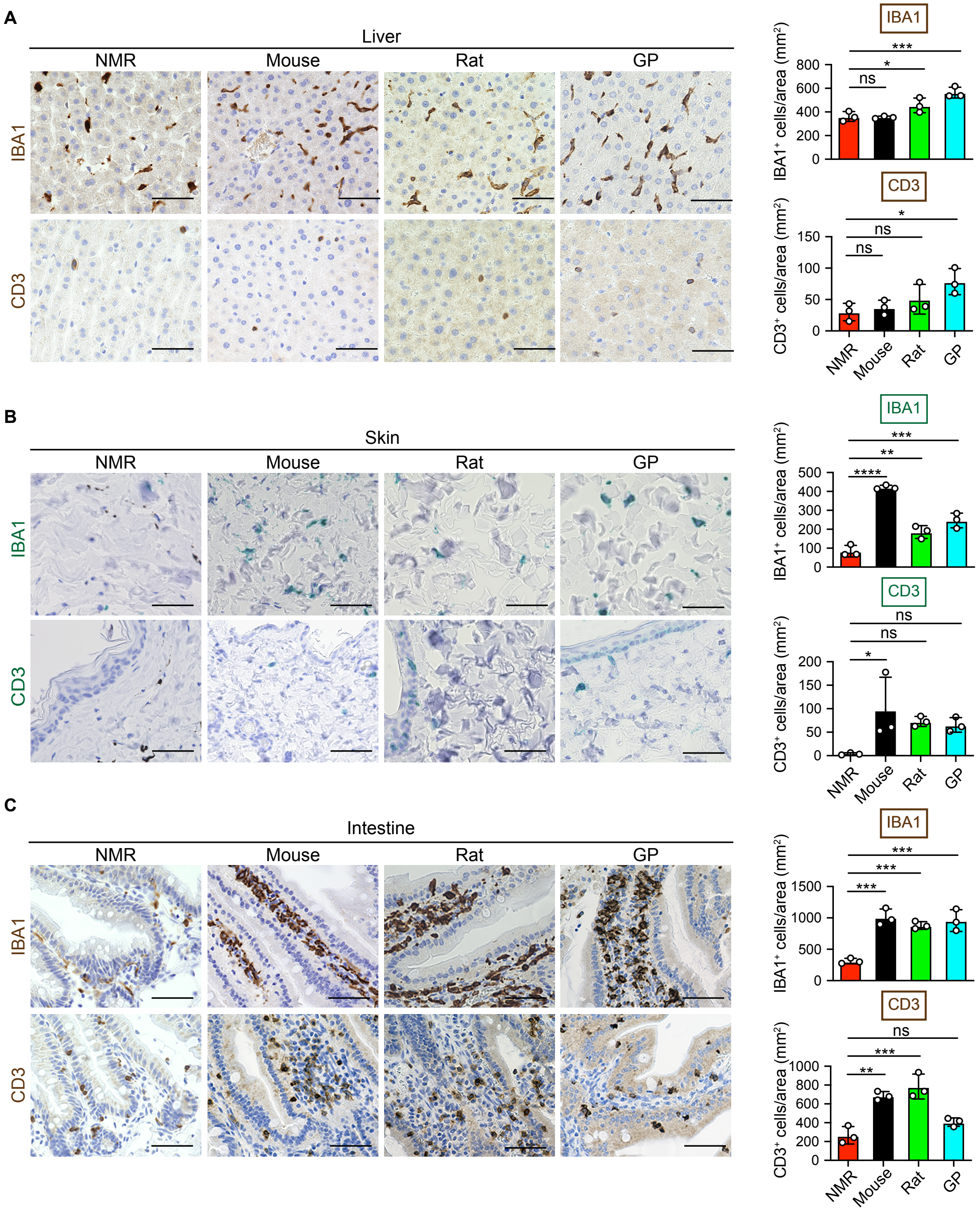


**Fig. S11. Immunohistochemical analysis of tissue-resident immune cells in several rodent species.**

**A**–**C**, Immunohistochemical staining and quantification of IBA1- and CD3-positive cells per area in liver (**A**, brown), skin (**B**, green), or small intestine (**C**, brown) sections of NMRs, mice, rats, and GPs. Scale bar: 50 μm. Data are presented as the mean ± SD of *n* = 3 animals. One-way ANOVA with Dunnett's multiple comparisons test versus NMR.

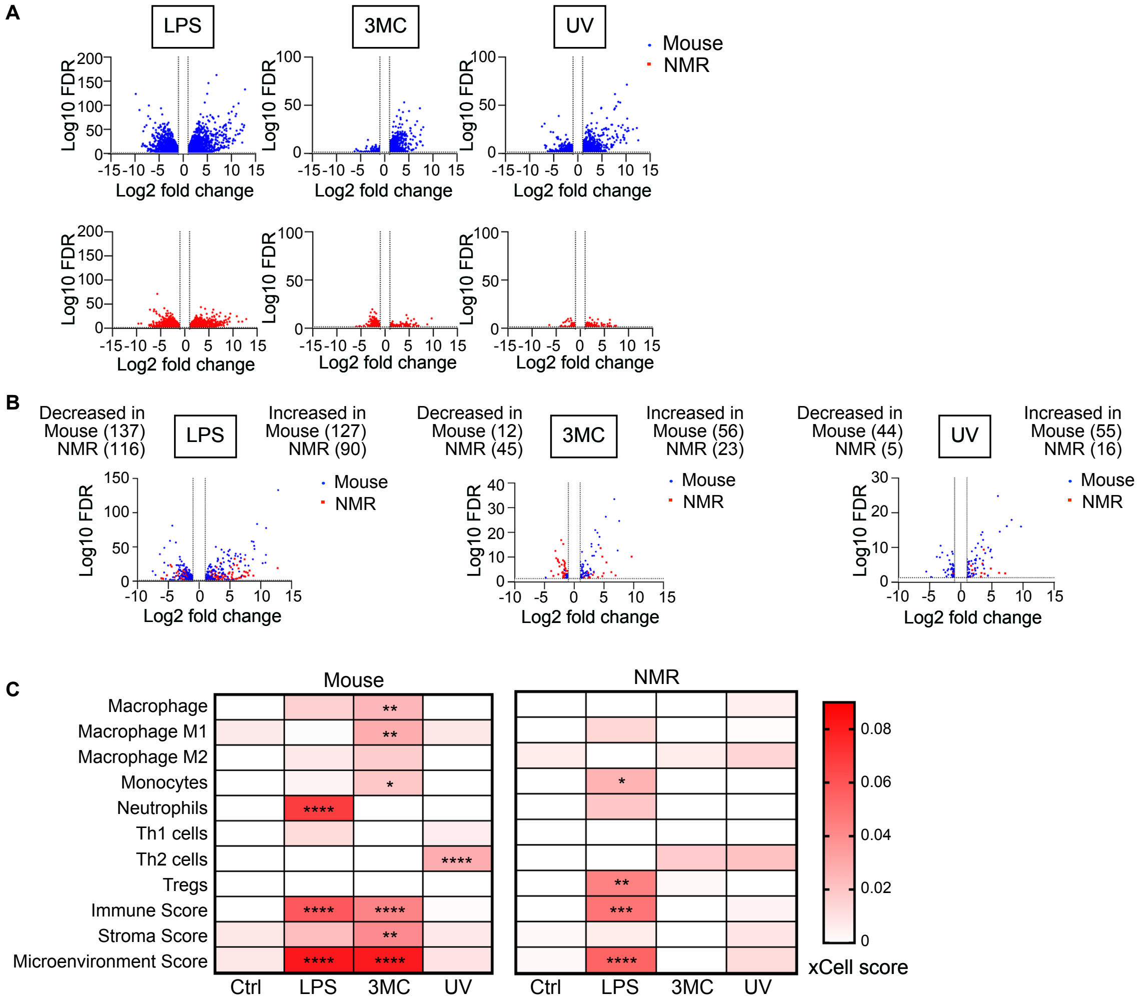


**Fig. S12. RNA-seq analyses of mouse and NMR skin after exposure to LPS, 3MC, or UV.**
**A** and **B**, Volcano plots of expression differences in all genes (**A**) and 467 selected ligands (**B**, Supplementary Table 2) between mouse skin (blue) and NMR skin (red) after exposure to LPS, 3MC (1 week), or UV. Each point indicates the gene with FDR-adjusted $P$ < 0.05 and |log2 fold change| > 1. $n$ = 3 animals per treatment. **C**, Heatmap of average scores calculated using xCell. The scores of representative immune cell types are shown. Only significant $P$-values are shown. For **C**, data are expressed as the mean of $n$ = 3 animals. One-way ANOVA with Dunnett's multiple comparisons test versus untreated control (Ctrl).

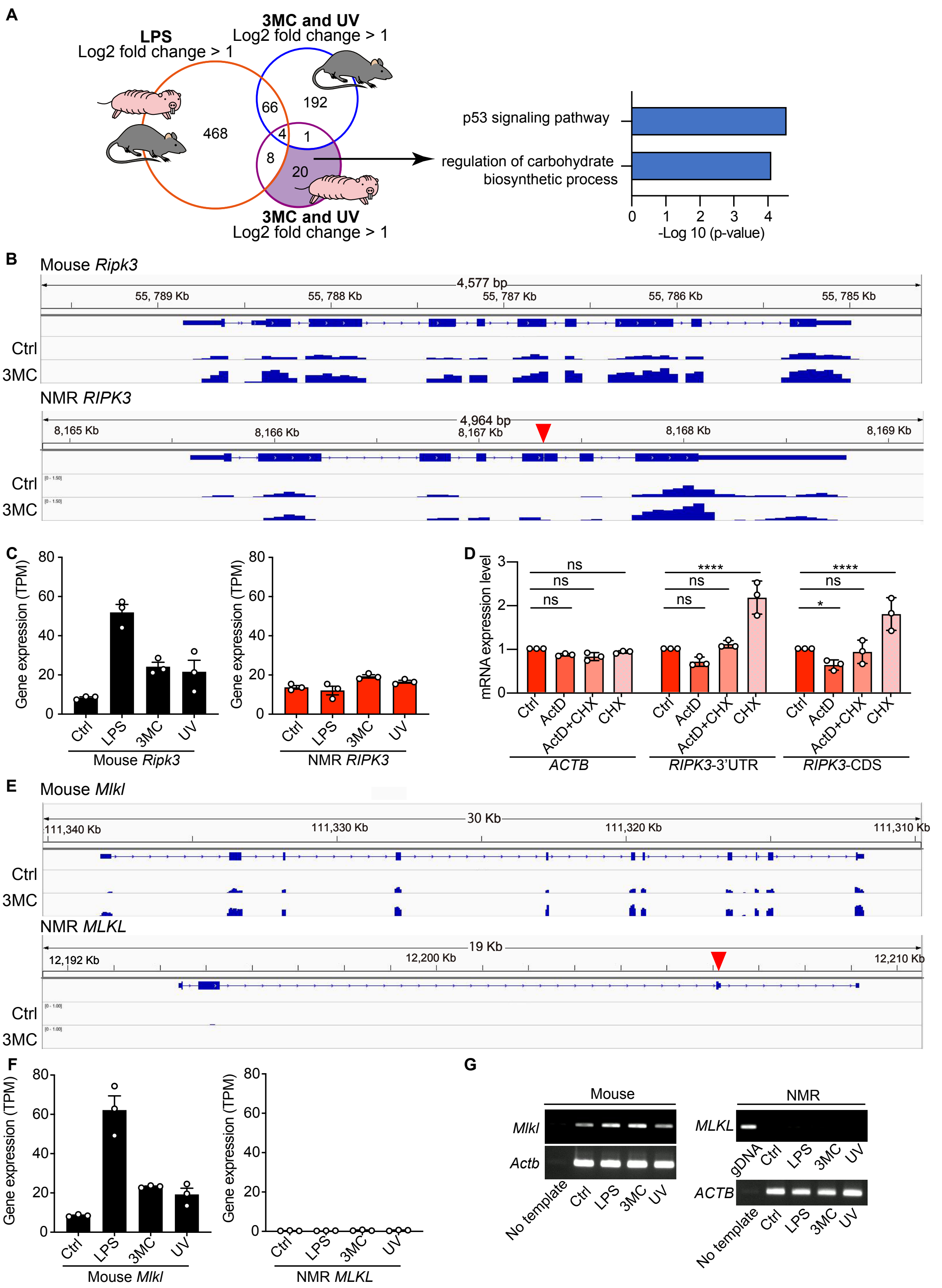


**Fig. S13. Gene expression analysis of receptor-interacting protein kinase 3 (*RIPK3*) and mixed lineage kinase domain-like (*MLKL*).**

**A**, RNA-seq analysis. Venn diagram showing the number of commonly upregulated genes in mouse and NMR skin upon LPS treatment (orange circle), and upregulated genes in mouse (blue circle) or NMR (purple circle) skin upon both 3MC (1 week) and UV treatment; enriched gene ontology (GO) terms and Kyoto Encyclopedia of Genes and Genomes (KEGG) pathways of 3MC-UV NMR-DEGs are shown (purple-filled area, 20 genes). **B**, Visualization of RNA-seq data at the *RIPK3* gene locus in the control (Ctrl) or 3MC-treated skin, using the Integrative Genomics Viewer (IGV). A red arrowhead indicates a frame-shift mutation in NMR *RIPK3*. **C**, *RIPK3* expression level (transcripts per million, TPM) in Ctrl-, LPS-, 3MC-, or UV-treated skin ($n$ = 3 animals in each treatment). **D**, RT-qPCR analysis of NMR fibroblasts incubated with actinomycin D (ActD) and/or cycloheximide (CHX). The values shown are the average fold change relative to that of untreated cells (Ctrl). Data are presented as the mean ± SD of $n$ = 3 biological replicates. The mRNA expression was normalized to *GAPDH* expression. Actin beta (*ACTB*) was used as an example of nonsense-mediated mRNA decay non-target mRNA. Two-way ANOVA followed by Dunnett's post hoc test versus untreated control. **E**, Visualization of RNA-seq data at the *MLKL* gene locus in Ctrl or 3MC-treated skin using IGV. A red arrowhead indicates a frame-shift mutation in NMR-*MLKL*. **F**, *MLKL* expression levels (TPM) in Ctrl-, LPS-, 3MC-, or UV-treated skin ($n$ = 3 animals per treatment). **G**, Semi-quantitative RT-PCR analysis of the expression of *MLKL* and *ACTB* in skin after each treatment. gDNA: tail-tip genomic DNA used as a PCR positive control.

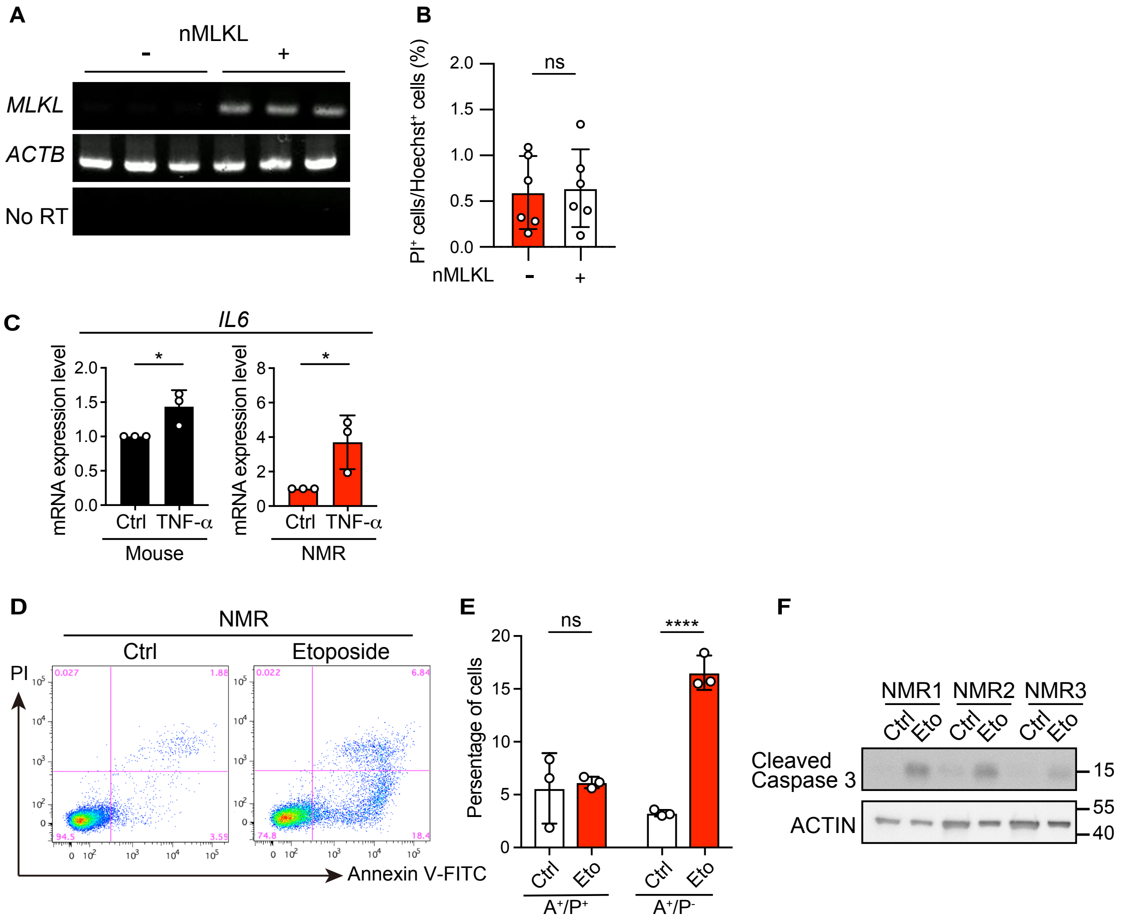


**Fig. S14. Responses to necroptotic or apoptotic stimuli in skin fibroblasts.**
**A**, Semi-quantitative RT-PCR analysis of the expression of *MLKL* and *ACTB* in NMR SV40ER cells with (+) or without (-) NMR-*MLKL* overexpression. Template without reverse transcriptase (No RT) was used as a negative control. **B**, Propidium iodide (PI) staining of NMR SV40ER cells overexpressing NMR-*MLKL*. **C**, Relative expression level of *IL6* mRNA in NMR and mouse fibroblasts treated with TNF-α for 24 h (normalized to *ACTB* mRNA). The values shown are the average fold change relative to that of the untreated control. **D**, Annexin V/PI staining of NMR fibroblasts treated with etoposide. **E**, Quantification of Annexin V- and/or PI-positive cells (%). $A^+/P^+$, Annexin and PI double-positive cells. $A^+/P^-$, Annexin single positive cells. Eto: etoposide. **F**, Western blot detection of cleaved caspase-3 and ACTIN in NMR fibroblasts treated with etoposide. $n$ = 3 biological replicates. Data are presented as the mean ± SD of $n$ = 3 biological replicates (for C, E) or $n$ = 6 independent experiments (for B). Unpaired $t$-test versus untreated control.

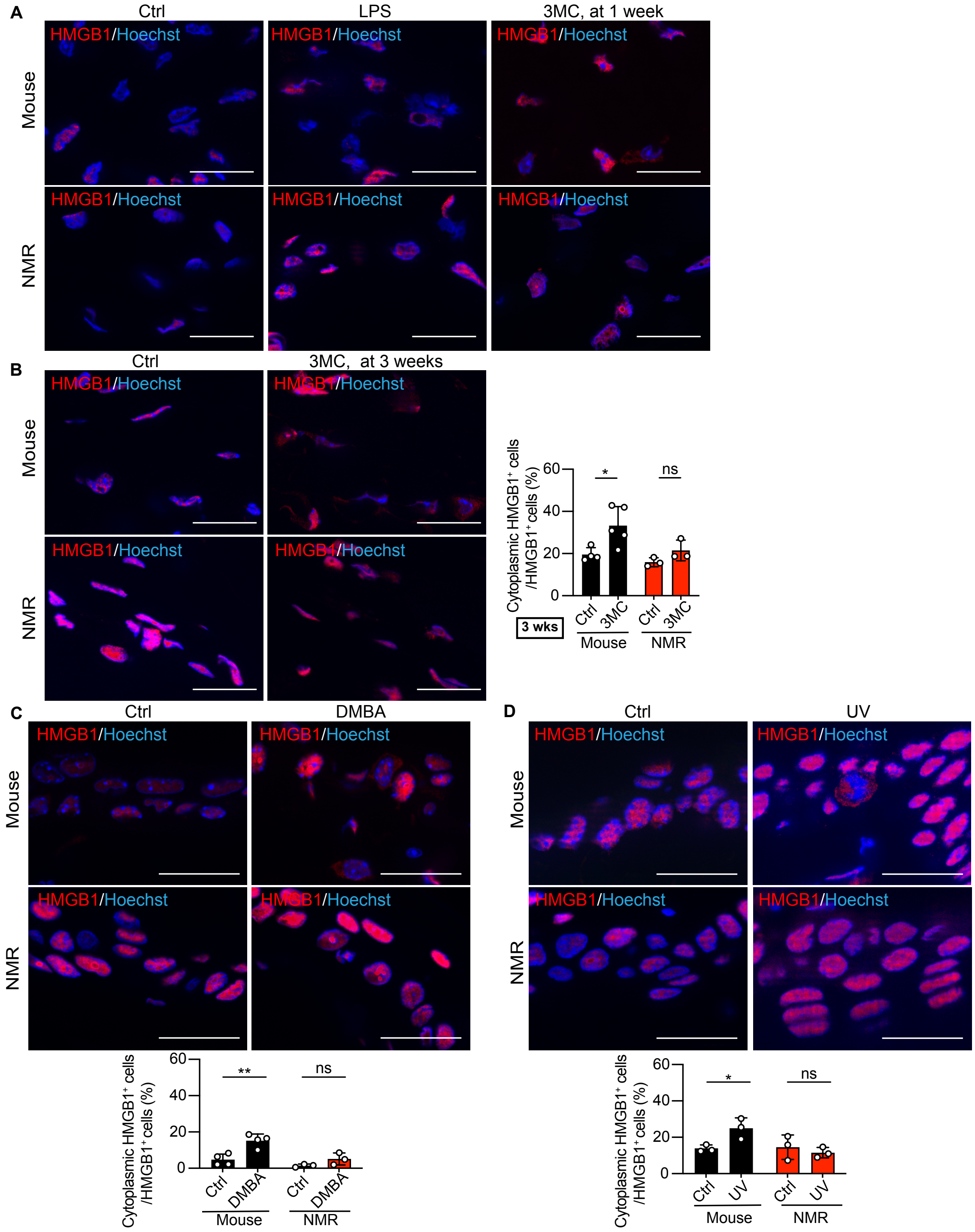


**Fig. S15. Immunofluorescence staining for high mobility group box-1 protein (HMGB1) in the skin after exposure to LPS, 3MC, DMBA, or UV.**

**A**, Immunofluorescence staining for HMGB1 (red) in the skin after exposure to 3MC (1 week) and LPS. Scale bar: 20 μm. **B**, Immunofluorescence staining and quantification of cytoplasmic HMGB1 (red) in the skin at 3 weeks after 3MC treatment. Scale bar: 20 μm. **C**, Immunofluorescence staining and quantification of cytoplasmic HMGB1 (red) in the skin at 24 h after DMBA treatment. Scale bar: 20 μm. **D**, Immunofluorescence staining and quantification of cytoplasmic HMGB1 (red) in the skin after UV treatment. Scale bar: 20 μm. Data are presented as the mean ± SD of *n* = 3–5 animals. Unpaired *t*-test versus untreated control (Ctrl).

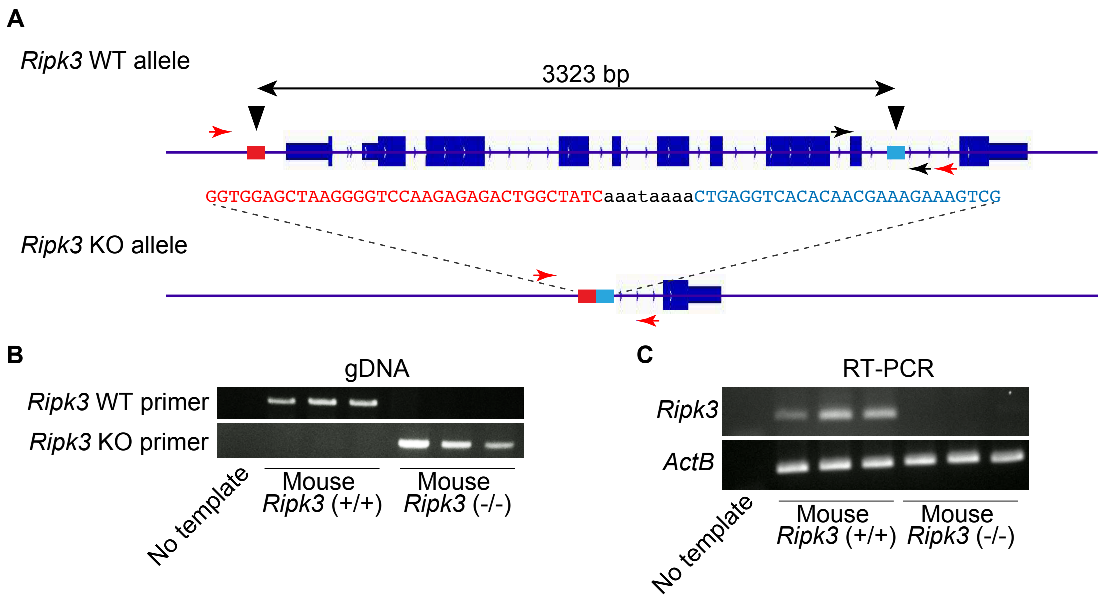


**Fig. S16. Knockout of mouse *Ripk3*.**
**A**, Schematic diagram of the *Ripk3* wild-type (WT) allele and the knockout (KO) allele. The targeted alleles of *Ripk3* were generated by introduction of Cas9, the synthetic crRNAs designed to target the 5′ upstream region and intron 9 (arrowheads), tracrRNA, and ssODN into C57BL/6N fertilized eggs. The red and black arrows indicate the primer sets used for detection of KO and WT alleles, respectively. **B**, PCR genotyping of the *Ripk3* alleles. **C**, Semi-quantitative RT-PCR analysis of the skin.

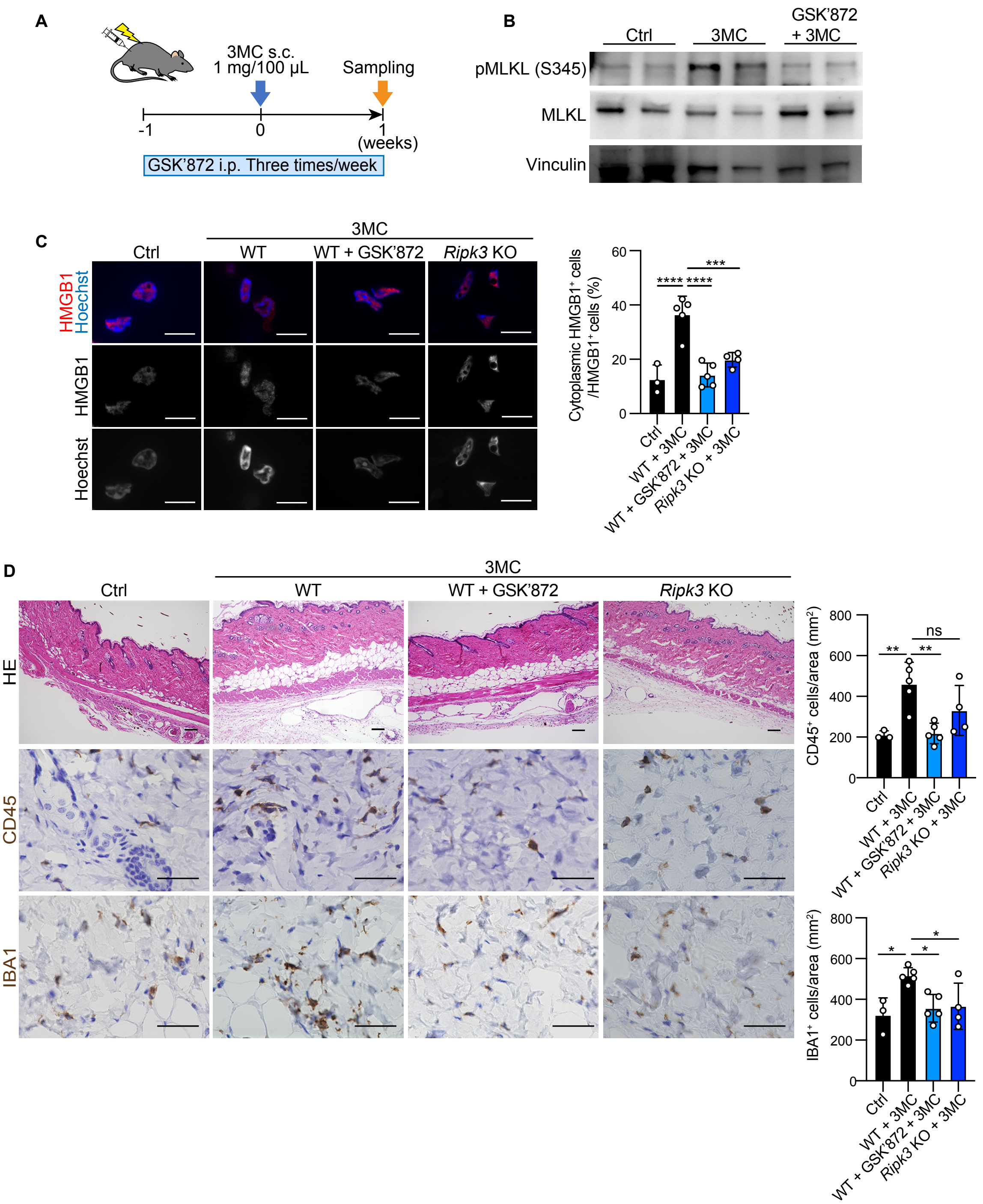


**Fig. S17. Inhibition of RIPK3 suppresses necroptosis and inflammatory immune cell responses in mouse skin.**

**A**, Schematic diagram for investigating responses to subcutaneous (s.c.) 3MC injection after suppression of necroptosis by GSK'872 in mouse skin. GSK'872 was intraperitoneally (i.p.) injected three times a week starting at 1 week before 3MC injection. **B**, Western blot detection of phospho-mixed lineage kinase domain-like (pMLKL [S345]), MLKL, and vinculin in mouse skin after exposure to 3MC with or without GSK'872. *n* = 2 animals per group. **C**, Immunofluorescence staining and quantification of cytoplasmic HMGB1 (red) in mouse skin after exposure to 3MC with or without GSK'872 in WT or *Ripk3* KO mice. Scale bar: 10 μm. **D**, HE staining, immunohistochemical staining (brown), and quantification of CD45- and IBA1-positive cells per area in mouse skin after exposure to 3MC with or without GSK'872 in WT or *Ripk3* KO mice. Scale bars: 100 μm (HE) and 50 μm (others). For **C** and **D**, data are presented as the mean ± SD of $n$ = 3 (control; Ctrl), 4 (*Ripk3* KO + 3MC), or 5 (WT + 3MC or WT + GSK'872 + 3MC) animals. One-way ANOVA with Dunnett's multiple comparison test versus 3MC.

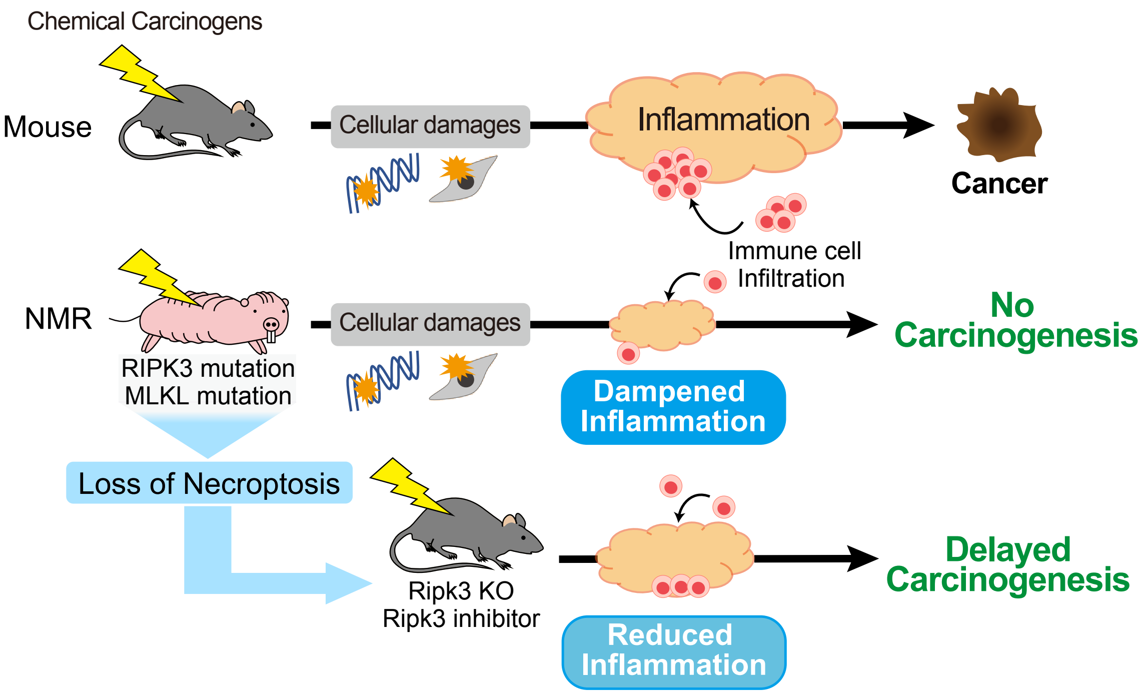


**Fig. S18. Graphical illustration which shows the main finding from this paper.**
After carcinogenic treatments, tissue damage such as DNA damage and cell death is induced in both NMRs and mice; however, only NMRs show attenuated inflammatory responses and do not develop tumors. Inhibition of RIPK3 in mice resulted in the reduced inflammatory response and the delayed onset of chemical carcinogenesis. Thus, in NMRs, loss-of-function mutations in genes essential for necroptosis induction may attenuate inflammatory responses and serve as an in vivo cancer resistance mechanism.

**Dataset S1: Antibodies used in this study.**

| Antibody | Code No. | Results | Antigen retrieval and concentration | Epitope | Percent homology (%) to the epitope |
|---|---|---|---|---|---|
| CD3 (SP7) | Nichirei (413591) | Positive | Antigen retrieval buffer pH9 (Nichirei), 1:500 | Human CD3, aa 150 to the C-terminus | NMR 86.2%, Mouse 91.4% |
| Iba1 | WAKO (019-19741) | Positive | EDTA pH8 buffer, 1:4000 | Rat Iba1, C-terminus | NMR 84.4%, Mouse 93.9% |
| Myeloperoxidase | DAKO (A0398) | Positive | EDTA pH8 buffer, 1:4000 | Human MPO | NMR 87.7%, Mouse 86.4% |
| CD45 | Abcam (ab10558) | Positive | Antigen retrieval buffer pH9 (Nichirei), 1:2000 | Human CD45, aa 900-1000 | NMR 87.1%, Mouse 91.1% |
| CD4 (EPR6855) | Abcam (ab133616) | Negative | | | |
| CD4 (GHH4) | dianova (DIA-404) | Negative | | | |
| CD8 (GHH8) | dianova (DIA-808) | Negative | | | |
| CD11b (M1/70) | Biolegend (101201) | Negative | | | |
| CD11b (EP1345Y) | Abcam (ab52478) | Negative | | | |
| CD34 (EP373Y) | Abcam (ab81289) | Negative | | | |
| CD45R(RA3-6B2) | Abcam (ab64100) | Negative | | | |
| CD45R(B220) | Pharmingen (01121D) | Negative | | | |
| CD68(FA-11) | Abcam (ab64100) | Negative | | | |
| CD79a(SP18) | Serotec (MCA1957) | Negative | | | |
| EMR1(F4/80) | Original | Negative | | | |
| Gr-1 | Southern Biotechnology (1900-01) | Negative | | | |
| **Antibody** | **Code No.** | **Results** | **Antigen retrieval and concentration** | **Epitope** | **Percent homology (%) to the epitope** |
| pH2AX | CST (#9718) | Positive | Antigen retrieval buffer pH9 (Nichirei), 1:500 | A synthetic phosphopeptide corresponding to residues surrounding Ser139 of human H2A.X | NMR 98.6%, Mouse 97.2% |
| 8-OHdG | Santa Cruz Biotechnology (sc-393871) | Positive | Antigen retrieval buffer pH9 (Nichirei), 1:5000 | | |
| Ki67 | Abcam (ab16667) | Positive | Antigen retrieval buffer pH9 (Nichirei), 1:200 | Human Ki67, aa 1200-1300 | NMR 47.5%, Mouse 44.6% |
| HMGB1 | Abcam (ab79823) | Positive | Antigen retrieval buffer pH9 (Nichirei), 1:500 | Human HMGB1, aa 150 to the C-terminus | NMR 100.0%, Mouse 97.0% |
| Cleaved Caspase-3 | CST (#9664) | Positive | none, 1:400 | Human Caspase-3, amino-terminal residues adjacent to Asp175 | NMR 100.0%, Mouse 100.0% |
| **Antibody** | **Code No.** | **Results** | **Concentration for WB** | **Epitope** | **Percent homology (%) to the epitope** |
| MLKL | Abcam (ab184718) | Positive | 1:1000 | | |
| pMLKL | Abcam (ab196436) | Positive | 1:1000 | | |
| CD45 | Abcam (ab10558) | Positive | 1:500 | Human CD45, aa 900-1000 | NMR 87.1%, Mouse 91.1% |
| Cleaved Caspase-3 | CST (#9664) | Positive | 1:1000 | Human Caspase-3, amino-terminal residues adjacent to Asp175 | NMR 100.0%, Mouse 100.0% |
| β-Actin | CST (#4970) | Positive | 1:2000 | Human β-actin, near the amino-terminus | NMR 100.0%, Mouse 100.0% |
| GAPDH | Invitrogen (MA5-15738) | Positive | 1:1000 | Rabbit GAPDH, Recombinant full length protein | NMR 98.2%, Mouse 94.9% |
| Vinculin | Sigma-Aldrich (V9131) | Positive | 1:1000 | Human Vinculin | NMR 99.5%, Mouse 99.1% |

**Dataset S2: Changes in expression of cell-to-cell communication genes after each stimulation.**

**NMR**

| Gene names | LPS.logFC | LPS.FDR | MC.logFC | MC.FDR | UV.logFC | UV.FDR |
|---|---|---|---|---|---|---|
| A2M | 3.08708171 | 7.41E-05 | 0.67542408 | 0.76276404 | 1.12563624 | 0.44888903 |
| AANAT | 0.06566789 | 0.99439059 | -0.1017118 | 1 | -2.7128991 | 0.48898714 |
| ACE | -0.8968818 | 0.04988499 | -1.9221723 | 0.00011397 | -0.3546998 | 0.74224926 |
| ADAM10 | 0.99461418 | 5.94E-06 | 0.02675559 | 0.9996892 | 0.36295387 | 0.39399963 |
| ADAM12 | 1.3668188 | 0.15267725 | -0.1094461 | 0.99980152 | 1.45918283 | 0.34346071 |
| ADAM15 | -1.4895282 | 3.77E-06 | -0.1546465 | 0.91254213 | -0.2809328 | 0.72977117 |
| ADAM17 | 0.67097088 | 0.00571364 | -0.096057 | 0.93311149 | -0.0634886 | 0.94394032 |
| ADAM2 | -1.7601143 | 0.04800046 | -0.9950735 | 0.6098233 | -0.8240104 | 0.6503075 |
| ADAM23 | -1.0715959 | 0.08180737 | -0.4830141 | 0.77725103 | -0.6132208 | 0.6284056 |
| ADAM9 | 1.00447164 | 3.08E-05 | 0.03984808 | 0.99094967 | 0.49097129 | 0.23173672 |
| ADIPOQ | -1.4661343 | 0.04195814 | -0.6803696 | 0.73311143 | -0.8954766 | 0.51091387 |
| ADM | 3.49628666 | 2.73E-27 | 0.90669383 | 0.05365064 | 0.64850085 | 0.24063047 |
| ADM2 | 1.40234442 | 0.16517665 | 0.63395871 | 0.83898555 | 0.46685713 | 0.87087488 |
| AGRN | 1.96795777 | 4.96E-07 | -0.175492 | 0.92362417 | 0.52896107 | 0.5092277 |
| AGT | 0.76413182 | 0.74796032 | -0.0936419 | 1 | 1.17215449 | 0.80787414 |
| AHSG | -0.8959797 | 0.15603667 | -0.2426995 | 0.92251912 | 1.5193156 | 0.03604614 |
| ALB | -3.2261223 | 0.15988644 | -3.2261223 | 0.45303874 | -3.2261223 | 0.38726928 |
| ALOX5AP | 0.27292209 | 0.67012495 | -0.6993615 | 0.56031601 | -0.2905266 | 0.84247594 |
| AMH | 0.66034972 | 0.59843013 | 1.27419923 | 0.57527293 | 0.49187216 | 0.87525745 |
| ANGPT1 | -0.7432155 | 0.16521037 | -1.0264413 | 0.21529448 | -0.2986268 | 0.8273972 |
| ANGPT2 | 1.62131625 | 2.53E-05 | 0.26917279 | 0.84537962 | 0.39282768 | 0.66947488 |
| ANGPTL1 | -4.6412769 | 2.01E-24 | -2.7657839 | 8.56E-10 | -0.7484787 | 0.31221773 |
| ANGPTL2 | -1.8687785 | 3.31E-13 | -1.5522908 | 2.63E-08 | -0.3754599 | 0.48302042 |
| ANGPTL4 | 2.09427699 | 2.16E-10 | 1.26520122 | 0.00212745 | 1.6708413 | 1.75E-05 |
| ANXA1 | 1.95048022 | 9.87E-09 | 0.52984804 | 0.49607993 | 0.47764542 | 0.50096044 |
| APLN | -2.7458547 | 3.58E-11 | -0.6745807 | 0.39949666 | -0.1140721 | 0.93489716 |
| APOB | -0.8419654 | 0.11217198 | -0.6457538 | 0.57209559 | -0.2181568 | 0.8853419 |
| APOD | -0.3146097 | 0.63094877 | 1.69619002 | 0.01225177 | 0.62761489 | 0.56383506 |
| APOE | 0.41618941 | 0.32364605 | 0.88390494 | 0.12182194 | 0.95595962 | 0.0723035 |
| APP | -0.2399975 | 0.44736088 | -0.4502057 | 0.42073187 | -0.0823955 | 0.93041823 |
| AREG | 0.95182178 | 0.41744975 | -1.1919651 | 0.72664945 | -0.4067678 | 0.91136055 |
| ARF1 | 0.41991152 | 0.07785584 | -0.1103066 | 0.90590608 | 0.01359608 | 0.9957431 |
| ARTN | -0.1709741 | 0.80278414 | 1.03205779 | 0.21618869 | -0.2919288 | 0.84334897 |
| AVP | -1.9117975 | 0.40648128 | -1.9117975 | 0.73311143 | 0.66772638 | 0.91486787 |
| AZGP1 | -2.7040807 | 0.24499161 | -2.7040807 | 0.56597942 | -2.7040807 | 0.50096044 |
| B2M | 2.59846357 | 2.44E-08 | 0.69904404 | 0.50599663 | 0.04523699 | 0.99089832 |
| BDNF | -1.2793371 | 0.27116514 | -0.9957997 | 0.73311143 | -0.8389787 | 0.7269224 |
| BGN | -1.9259457 | 9.83E-12 | -1.4530449 | 4.79E-06 | 0.62800935 | 0.17224373 |
| BMP15 | -1.9118761 | 0.40648128 | -0.0771879 | 1 | 0.70084218 | 0.91174243 |
| BMP2 | -2.0650944 | 4.83E-08 | 0.74737035 | 0.23163024 | 0.35080736 | 0.68165907 |
| BMP3 | -1.7528157 | 0.00113227 | -1.8436202 | 0.00612993 | -0.2297395 | 0.88706793 |
| BMP4 | -1.297827 | 0.00042374 | -0.2354523 | 0.85344141 | -1.213202 | 0.01150475 |
| BMP5 | -1.2162278 | 0.1210247 | -0.468348 | 0.83772395 | -0.3639504 | 0.85807029 |
| BMP6 | -0.2501015 | 0.83011663 | -2.1442477 | 0.16573061 | -0.6810443 | 0.75792521 |
| BMP7 | -0.7056843 | 0.02981459 | 0.27545907 | 0.76260222 | 0.03794812 | 0.98307305 |
| BMP8B | 2.52888355 | 0.19832529 | 0.69671916 | 0.93846792 | 2.48727279 | 0.47811861 |
| BST1 | 3.49146618 | 6.33E-07 | 0.5383325 | 0.81307512 | 1.45255231 | 0.20352319 |
| BTC | -6.1334566 | 5.02E-05 | -0.7082684 | 0.85782376 | -1.2261049 | 0.62521203 |
| BTLA | -0.4973405 | 0.62994299 | -0.1084303 | 0.99659947 | 0.39592624 | 0.86062633 |
| C1QA | 0.22042592 | 0.73364556 | 0.08702544 | 0.98726945 | 0.42631938 | 0.72379338 |
| C1QB | 0.22832042 | 0.66126253 | -0.2492846 | 0.86769392 | 0.593583 | 0.46573956 |
| C1QTNF1 | -1.080569 | 0.00025118 | -1.8785197 | 1.69E-09 | -0.3470644 | 0.58504778 |
| C3 | 3.17233522 | 1.17E-12 | 1.32197395 | 0.02929067 | 0.50319844 | 0.60588391 |
| C4A | 0.50632243 | 0.1996525 | 0.66825281 | 0.31610863 | 0.97276835 | 0.04913834 |
| C4B | 2.13300834 | 0.12389155 | 1.53350077 | 0.6588408 | 0.75457429 | 0.85921805 |
| CALCA | 6.80518497 | 5.50E-06 | 0 | 1 | 2.60724199 | 0.5023289 |
| CALCB | 6.18661543 | 0.00010179 | 1.83314186 | 0.74335844 | 3.48024467 | 0.2930509 |
| CALM1 | 0.0635663 | 0.85346055 | -0.2248455 | 0.75543605 | -0.1086683 | 0.88930828 |
| CALM2 | -0.9040332 | 0.00180782 | -0.1677322 | 0.87974443 | -0.4568159 | 0.40226368 |
| CALM3 | -0.3707762 | 0.12250833 | -0.1855013 | 0.7832134 | 0.06329617 | 0.9358964 |
| CALR | 1.40699714 | 2.33E-06 | 0.18355015 | 0.87146765 | 0.61347036 | 0.22777903 |
| CAMP | 4.3877506 | 0.01766467 | 3.48830707 | 0.33312994 | 3.77132064 | 0.21303867 |
| CCK | -0.0983627 | 0.97468065 | 0.1799251 | 0.99871815 | -0.9007015 | 0.82801506 |
| CCL1 | 1.30907293 | 0.51240248 | -1.9191873 | 0.73311143 | 0.67885522 | 0.90599008 |
| CCL11 | 4.24339803 | 2.24E-08 | 0.07575421 | 1 | 0.48595332 | 0.81039742 |
| CCL17 | 4.87757618 | 3.45E-12 | 0.57167053 | 0.85736166 | 0.93584087 | 0.6287061 |
| CCL2 | 4.95530911 | 1.01E-13 | 2.32268011 | 0.00243014 | 0.90109126 | 0.46332665 |
| CCL20 | 4.32215024 | 2.90E-08 | 0.70390382 | 0.74490884 | 0.1990562 | 0.94144058 |
| CCL22 | 5.4656819 | 1.62E-13 | -0.235236 | 0.94233544 | -1.182084 | 0.34075763 |
| CCL24 | -0.2671481 | 0.7132904 | 0.94670162 | 0.4038403 | 0.19958648 | 0.91473673 |
| CCL25 | 1.17099865 | 0.5252299 | 0.41089411 | 0.96903112 | 0.78529703 | 0.87074472 |
| CCL26 | -1.3973728 | 0.18989851 | -0.6075234 | 0.83501748 | -0.2228845 | 0.94383316 |
| CCL28 | 0.07307573 | 0.98699098 | -1.7621454 | 0.70712649 | -1.7635748 | 0.63432122 |
| CCL4 | 3.76548375 | 0.00140035 | 4.95392222 | 0.00033232 | -1.2994095 | 0.71156913 |
| CCL5 | 6.9997681 | 4.50E-09 | 0.98292126 | 0.7500911 | 0.78023964 | 0.78392921 |
| CCL8 | 5.39108153 | 9.25E-05 | 4.73891845 | 0.00523708 | 1.7321154 | 0.50096044 |
| CD14 | 2.59250332 | 4.01E-12 | 0.7428112 | 0.29634659 | 0.71234778 | 0.28714486 |
| CD34 | -1.3923405 | 3.49E-07 | -0.9217171 | 0.01072979 | 0.11216789 | 0.90439053 |
| CD40LG | -2.1813111 | 0.1575836 | -1.9164677 | 0.54313178 | 0.28182129 | 0.95274953 |
| CD55 | -1.0428207 | 1.63E-05 | -0.1150769 | 0.91571843 | -0.3529253 | 0.47926157 |
| CD5L | -3.9742385 | 2.66E-06 | -3.8820863 | 4.27E-05 | 0.90428979 | 0.50769049 |
| CDH1 | -0.3520998 | 0.27117266 | -0.1863119 | 0.85184156 | -0.1628531 | 0.84428032 |

**Mouse**

| Gene names | LPS.logFC | LPS.FDR | MC.logFC | MC.FDR | UV.logFC | UV.FDR |
|---|---|---|---|---|---|---|
| A2m | 3.22117594 | 0.00738992 | 1.29491186 | 0.59859869 | -2.540836 | 0.29159903 |
| Aanat | -1.7663097 | 0.37740051 | 0.11928322 | 0.99544146 | -1.7663097 | 0.47190341 |
| Ace | -0.8449697 | 0.00192818 | -0.0175873 | 0.99544146 | -0.6373833 | 0.04703573 |
| Adam10 | 0.20435697 | 0.29246449 | 0.44560915 | 0.05013115 | 0.13998856 | 0.58316808 |
| Adam12 | 0.14378115 | 0.68677391 | 0.27907566 | 0.61242883 | -0.1575977 | 0.73812328 |
| Adam15 | -0.4929346 | 0.00203526 | 0.1946249 | 0.48259269 | -0.2446811 | 0.22972797 |
| Adam17 | 0.73226938 | 8.30E-06 | 0.21028213 | 0.46966079 | 0.07615819 | 0.78410504 |
| Adam2 | 2.73000778 | 0.16140809 | 0 | 1 | 1.90544007 | 0.4247637 |
| Adam23 | -0.8251217 | 8.09E-06 | 0.13699588 | 0.71986902 | -0.3834985 | 0.08996619 |
| Adam9 | 1.44104999 | 4.62E-19 | 0.55785224 | 0.00470856 | 0.08396387 | 0.75689724 |
| Adipoq | -1.9003013 | 5.50E-14 | -0.8718995 | 0.00374553 | -1.1702945 | 1.90E-05 |
| Adm | 4.75451696 | 2.23E-44 | 0.90337543 | 0.02624442 | 0.09136379 | 0.88133821 |
| Adm2 | 6.51757856 | 8.32E-15 | 1.00236609 | 0.69334016 | 4.99946898 | 7.61E-08 |
| Agrn | -0.1188669 | 0.58261292 | 0.0892049 | 0.84476367 | -0.1705959 | 0.50800503 |
| Agt | -0.9951734 | 0.00515041 | -1.6273645 | 2.12E-05 | -1.6998993 | 6.12E-06 |
| Ahsg | -0.4067692 | 0.62551133 | -1.2533262 | 0.28541266 | -2.2795685 | 0.02064688 |
| Alb | -1.351785 | 0.00134973 | -0.2528878 | 0.78267308 | -0.9039649 | 0.06869427 |
| Alox5ap | 1.18547073 | 8.50E-06 | 1.30152457 | 6.19E-06 | 0.3268907 | 0.36628156 |
| Amh | 1.07154555 | 0.44826508 | 2.39192365 | 0.10329246 | 1.60718857 | 0.29695933 |
| Angpt1 | 0.68721438 | 0.01234155 | -0.6423382 | 0.07701907 | -1.0520785 | 0.00041521 |
| Angpt2 | 3.02996764 | 5.11E-34 | 0.43657667 | 0.27092976 | -0.0770836 | 0.87128515 |
| Angptl1 | -6.3575717 | 3.38E-40 | -0.6852762 | 0.22684126 | -0.2186456 | 0.70943567 |
| Angptl2 | -3.7606063 | 2.46E-57 | -0.4758639 | 0.12986534 | -0.7055883 | 0.00515655 |
| Angptl4 | 3.66112757 | 8.60E-47 | 0.83754721 | 0.00398866 | -0.1821851 | 0.62171725 |
| Anxa1 | -0.2442167 | 0.35393949 | 0.43752287 | 0.213314 | 0.17503936 | 0.61465084 |
| Apln | 3.36249993 | 3.61E-13 | 1.37625373 | 0.01466014 | 0.80209099 | 0.16852921 |
| Apob | -4.6378027 | 0.01853112 | -2.7751169 | 0.25802119 | -2.7293317 | 0.17982234 |
| Apod | 2.24984697 | 1.22E-17 | 0.66728014 | 0.05498957 | -0.1117066 | 0.80245769 |
| Apoe | -1.4978398 | 6.80E-09 | -0.1360882 | 0.82898455 | -0.3285967 | 0.34645781 |
| App | -0.0601308 | 0.82425101 | 0.1989114 | 0.60851137 | -0.0613357 | 0.87125896 |
| Areg | 1.70978571 | 6.04E-05 | 0.2081172 | 0.85242218 | 2.26511334 | 3.79E-07 |
| Arf1 | 0.30417773 | 0.1506959 | -0.0940255 | 0.84664852 | 0.13234459 | 0.65541067 |
| Artn | 0.68662122 | 0.25654188 | 1.17713991 | 0.10619479 | 1.35981182 | 0.02661554 |
| Avp | -2.7156304 | 0.03761175 | -1.1022101 | 0.55125473 | -1.9522945 | 0.16691797 |
| Azgp1 | -3.0429672 | 0.1271334 | -1.1557549 | 0.72622359 | -0.3500765 | 0.90344094 |
| B2m | 1.65601349 | 5.11E-06 | 0.80107216 | 0.11103208 | 0.71843992 | 0.10710183 |
| Bdnf | 0.11961208 | 0.76442214 | 0.0781353 | 0.94025129 | 0.10020343 | 0.85941581 |
| Bgn | -0.5465849 | 0.0679359 | -0.0604962 | 0.95298831 | 0.04939365 | 0.93760895 |
| Bmp15 | -1.4775988 | 0.36019435 | 1.07859429 | 0.58491715 | 0.20977874 | 0.93628681 |
| Bmp2 | -0.1598935 | 0.70637815 | -0.315501 | 0.63368797 | -0.2696943 | 0.59616061 |
| Bmp3 | -1.3599672 | 1.56E-06 | -0.1765364 | 0.77746803 | -0.8141834 | 0.01206092 |
| Bmp4 | -2.1896719 | 4.35E-17 | -0.5101709 | 0.16672416 | -0.8067763 | 0.00538735 |
| Bmp5 | -4.2416957 | 1.14E-06 | -1.3817369 | 0.19797621 | -0.1234292 | 0.93677698 |
| Bmp6 | -1.5730799 | 3.64E-07 | -0.1736183 | 0.80016463 | -0.5046141 | 0.18709882 |
| Bmp7 | -0.3349894 | 0.15653398 | -0.3886395 | 0.24155194 | 0.22014577 | 0.46672491 |
| Bmp7 | 0.16047915 | 0.95283723 | 0.10191549 | 0.99544146 | -1.783512 | 0.47190341 |
| Bst1 | 3.88363098 | 2.55E-34 | 1.90427546 | 2.86E-09 | 0.93783167 | 0.00798756 |
| Btc | -3.6535411 | 3.81E-24 | -0.5890059 | 0.25762074 | -1.8326976 | 4.18E-07 |
| Btla | -0.6250014 | 0.16467763 | 1.46529432 | 0.0008229 | 0.34216573 | 0.54516761 |
| C1qa | -1.8521544 | 6.14E-12 | 0.94375896 | 0.00291666 | -0.2942999 | 0.42800577 |
| C1qb | -1.7355234 | 3.20E-10 | 0.98749434 | 0.00226071 | -0.2907526 | 0.44971394 |
| C1qtnf1 | -1.6383674 | 1.26E-17 | -0.2326344 | 0.50665221 | -0.7184468 | 0.00083998 |
| C3 | 2.1024299 | 5.85E-08 | 1.20741528 | 0.0098336 | 0.13937547 | 0.83164464 |
| C4b | 0.9971362 | 0.02062759 | 0.86327614 | 0.14960836 | 0.60941033 | 0.28160909 |
| C4b | 0.30916284 | 0.36064133 | 0.39244444 | 0.4471611 | -0.3855788 | 0.33523592 |
| Calca | 5.43353026 | 2.91E-18 | 0.68052401 | 0.75607076 | 1.43278934 | 0.22148227 |
| Calcb | 0.65204679 | 0.06278545 | 0.15557112 | 0.8529477 | -0.0759833 | 0.91123731 |
| Calm1 | 0.19984905 | 0.43207682 | 0.08142151 | 0.89185201 | 0.04317825 | 0.92897975 |
| Calm2 | -0.2690364 | 0.25983079 | 0.1838406 | 0.66671579 | -0.4014142 | 0.13762134 |
| Calm3 | -0.5578471 | 0.00256499 | -0.2263676 | 0.47939804 | -0.3213298 | 0.16158473 |
| Calr | 1.46366004 | 7.86E-09 | 0.18947947 | 0.72577658 | 0.65261222 | 0.03094537 |
| Camp | 4.98935361 | 3.59E-23 | 0.29928423 | 0.84686999 | 0.63006488 | 0.45356414 |
| Cck | -1.1302649 | 0.0244026 | -0.4661196 | 0.58532084 | -0.5478944 | 0.38395985 |
| Cckbr | -1.6976606 | 0.03216798 | 0.51634882 | 0.62724493 | 2.343704 | 3.31E-06 |
| Ccl11 | -0.7192836 | 0.0298155 | 1.41852949 | 3.51E-05 | 0.67097203 | 0.07873655 |
| Ccl17 | 2.38828929 | 0.00014024 | 2.33597418 | 0.00111162 | 2.70023359 | 6.64E-05 |
| Ccl2 | 5.95455689 | 7.90E-41 | 3.92922529 | 1.36E-20 | 1.37373942 | 0.00176145 |
| Ccl20 | 3.62744579 | 3.22E-17 | 0.90508038 | 0.2030531 | 3.49659802 | 3.75E-15 |
| Ccl22 | 0.20998981 | 0.53871906 | 1.50836103 | 7.85E-07 | 1.80704191 | 1.23E-09 |
| Ccl24 | -5.6307262 | 7.16E-28 | 0.44661278 | 0.55536437 | -1.2342965 | 0.00999617 |
| Ccl24 | 0.46307153 | 0.16854393 | 0.18218672 | 0.80318268 | 0.09981306 | 0.86187705 |
| Ccl24 | -2.5422227 | 0.19953569 | -2.5422227 | 0.39077643 | 1.06646539 | 0.56554841 |
| Ccl28 | 1.78437837 | 0.00329729 | 0.44969035 | 0.76671709 | -0.0434987 | 0.98409087 |
| Ccl4 | 8.84917164 | 3.04E-58 | 4.10011978 | 1.02E-15 | 1.76733729 | 0.00347179 |
| Ccl5 | 5.20873774 | 4.66E-34 | 2.26744827 | 9.66E-08 | 1.95402315 | 5.38E-06 |
| Ccl8 | -1.6030951 | 0.00064732 | 0.9283204 | 0.1643638 | 0.91379802 | 0.10764916 |
| Cd14 | 4.04779533 | 4.71E-43 | 0.90877466 | 0.00716098 | -0.4757467 | 0.19246378 |
| Cd34 | -1.9389045 | 3.97E-13 | -0.1176821 | 0.86453462 | -0.6332877 | 0.04937708 |
| Cd40lg | 0.20372539 | 0.9337657 | 1.97653004 | 0.40002829 | 2.29927458 | 0.19409323 |
| Cd55 | -0.601391 | 0.00365188 | 0.45778854 | 0.09928874 | -0.3047268 | 0.2462772 |
| Cd5l | -7.0832333 | 7.39E-10 | 1.69046021 | 0.00510486 | -2.7345093 | 0.00028711 |
| Cdh1 | 0.15689756 | 0.55846393 | -0.2404493 | 0.57250112 | 0.58482742 | 0.03052844 |

| | | | | | | |
|---|---|---|---|---|---|---|
| CFH | -0.3417222 | 0.38595361 | -0.868968 | 0.08443331 | -0.1745488 | 0.86620251 |
| CGN | -1.2973641 | 1.69E-05 | -0.4757514 | 0.47724803 | -1.2370504 | 0.00097375 |
| CHAD | -3.5008453 | 0.12341862 | -2.8692933 | 0.51911003 | -0.6098165 | 0.91842029 |
| CLCF1 | 3.98508032 | 5.75E-11 | 0.6100569 | 0.77331387 | 0.81562902 | 0.57929991 |
| CLEC11A | -1.5491133 | 0.06143912 | 0.54341948 | 0.79655754 | -0.0216755 | 0.9985772 |
| CNTF | 0.17230158 | 0.59420065 | -0.1313905 | 0.90465056 | 0.1246725 | 0.8787046 |
| CNTN2 | -3.0610118 | 7.49E-06 | -0.0185064 | 1 | -0.7806996 | 0.50106056 |
| CNTN4 | -2.2321858 | 0.00114063 | -1.787908 | 0.05743226 | -1.052401 | 0.35453737 |
| COL11A1 | -0.8683029 | 0.23047068 | -2.6846156 | 0.00088754 | -0.6532881 | 0.66004287 |
| COL14A1 | -1.0149072 | 3.73E-06 | -1.8392356 | 5.05E-16 | -0.5378534 | 0.11390127 |
| COL18A1 | -1.4717101 | 9.79E-10 | -0.4584193 | 0.34277961 | 0.23583689 | 0.68910491 |
| COL1A1 | -1.805009 | 4.07E-05 | -3.0096215 | 2.55E-10 | -0.0647491 | 0.97925622 |
| COL1A2 | -1.8188401 | 5.82E-06 | -3.1532385 | 3.13E-13 | -0.5508636 | 0.50096044 |
| COL2A1 | -2.4429945 | 0.00503479 | -0.6544393 | 0.75675966 | -0.9239154 | 0.53795572 |
| COL3A1 | -2.4393997 | 9.91E-09 | -2.731486 | 3.49E-09 | -0.5720139 | 0.50803421 |
| COL4A1 | 0.07423788 | 0.88437268 | -0.3774554 | 0.70818488 | 0.10720583 | 0.93090502 |
| COL4A2 | -0.3891412 | 0.3355177 | -0.3901529 | 0.68555999 | -0.0347031 | 0.99018003 |
| COL4A3 | -3.4300329 | 3.69E-15 | 0.22969346 | 0.8893779 | -0.757093 | 0.30333882 |
| COL4A4 | -2.5631483 | 1.26E-09 | 0.13988827 | 0.94693408 | -0.5053277 | 0.56374358 |
| COL4A5 | -1.9559571 | 1.92E-07 | -0.0354565 | 1 | -0.0248595 | 0.9957431 |
| COL4A6 | -2.7809342 | 3.35E-08 | -0.5324804 | 0.66337385 | -1.1010464 | 0.12345986 |
| COL5A1 | -1.9032757 | 1.50E-07 | -2.1071307 | 1.09E-07 | -0.3895848 | 0.6462757 |
| COL5A2 | -1.5610793 | 4.29E-09 | -1.6812421 | 4.77E-09 | -0.2490665 | 0.70543166 |
| COL5A3 | 0.03217451 | 0.95494119 | 0.14625819 | 0.91346746 | 0.18469733 | 0.83867509 |
| COL6A1 | -1.4848666 | 2.61E-07 | -1.0544527 | 0.00399871 | -0.0188821 | 0.9957431 |
| COL6A2 | -1.2786621 | 8.34E-05 | -1.1708165 | 0.00390613 | 0.09630426 | 0.93523192 |
| COL6A3 | -1.5151604 | 4.70E-07 | -1.4248924 | 3.66E-05 | 0.0518462 | 0.973251 |
| COL7A1 | -1.6487302 | 0.00010019 | 0.26407809 | 0.86190448 | 0.60150174 | 0.48210999 |
| COL8A1 | -0.8324957 | 0.03314989 | -1.3625292 | 0.00295934 | 0.22967674 | 0.82410815 |
| COL9A1 | -1.9215982 | 0.40648128 | -1.9215982 | 0.73311143 | 1.16968456 | 0.81716818 |
| COL9A2 | -0.8111696 | 0.09677405 | -0.2541102 | 0.88113091 | -0.0490574 | 0.98649627 |
| COL9A3 | -1.2518298 | 0.01380023 | -1.1329655 | 0.14890024 | -0.3596006 | 0.78121572 |
| COLQ | 0.44670417 | 0.1693314 | 0.57575038 | 0.27629773 | 0.11497728 | 0.90720537 |
| CORT | -1.0292076 | 0.62486006 | -1.4352766 | 0.78515344 | -1.4306849 | 0.73625357 |
| CP | 1.29638773 | 0.00294995 | 0.02701758 | 1 | 1.85623653 | 0.00031218 |
| CRP | 0 | 1 | 0 | 1 | 2.6131438 | 0.52730961 |
| CSF1 | 0.80522655 | 0.0389081 | 0.52856753 | 0.53970162 | 0.75814542 | 0.21255617 |
| CSF2 | 3.25023469 | 0.15285232 | 1.83468659 | 0.74490884 | 0 | 1 |
| CSF3 | 5.77431072 | 1.88E-33 | 1.40588786 | 0.03771107 | -0.1817106 | 0.91891717 |
| CTF1 | 0.27625349 | 0.64440614 | 0.43031969 | 0.75069414 | 0.40104659 | 0.72613814 |
| CTHRC1 | -2.7386424 | 3.66E-11 | -2.5328184 | 1.44E-08 | 0.31693289 | 0.76857508 |
| CUBN | -1.2020263 | 0.00112803 | -0.7087599 | 0.28883607 | -0.6148395 | 0.35099046 |
| CX3CL1 | 1.49552446 | 1.50E-06 | -0.0933345 | 0.96327045 | 0.28746629 | 0.71495239 |
| CXCL1 | 6.40052754 | 1.56E-19 | 4.74461887 | 6.45E-11 | 1.85588247 | 0.06356017 |
| CXCL10 | 8.82850004 | 7.15E-09 | 6.23708245 | 0.00012039 | 1.52643136 | 0.53929764 |
| CXCL11 | 8.01290002 | 1.72E-09 | 3.99604131 | 0.01246231 | 1.63241033 | 0.5172217 |
| CXCL12 | -0.918071 | 0.01345642 | -0.6711869 | 0.32835023 | 0.04025366 | 0.98649108 |
| CXCL13 | 5.24667533 | 0.0069052 | 0 | 1 | 2.59694005 | 0.52411461 |
| CXCL16 | 1.95800159 | 2.61E-08 | 0.21348446 | 0.87847766 | 0.61003223 | 0.35635705 |
| DCN | 0.8272341 | 0.03938057 | -0.7045976 | 0.33746382 | 0.17799289 | 0.88128631 |
| DEFB1 | 1.52469294 | 0.23791452 | 1.87318045 | 0.4153436 | 3.25450662 | 0.02197921 |
| DHH | -0.5526349 | 0.66216495 | -0.4987494 | 0.90307342 | -0.6370951 | 0.81665904 |
| DKK1 | 0.46961262 | 0.6386128 | 1.64656996 | 0.15802131 | 1.8221353 | 0.08133475 |
| DKK2 | -2.5185478 | 2.53E-17 | -1.5043482 | 3.34E-06 | -0.6735734 | 0.13961016 |
| DLK1 | -1.5783433 | 0.33053501 | -4.3503871 | 0.12368013 | -0.1334143 | 0.98649627 |
| DLK2 | -1.9962763 | 0.00268005 | 0.05057148 | 1 | -1.2504477 | 0.24790662 |
| DLL1 | 0.73599321 | 0.06072136 | 0.30524694 | 0.79095801 | -0.281263 | 0.7719279 |
| DLL3 | -0.1526615 | 0.76052343 | 0.20449305 | 0.89048216 | -0.3582718 | 0.68910491 |
| DLL4 | 1.93796287 | 8.19E-14 | -0.7494293 | 0.08085031 | -0.3047419 | 0.63852087 |
| DSCAM | 0.74986351 | 0.35997274 | 0.8012779 | 0.67096517 | 0.06961105 | 0.99089832 |
| DUSP18 | 0.62055301 | 0.01570377 | 0.13586281 | 0.89328189 | 0.05686244 | 0.95572439 |
| EBI3 | 5.69885303 | 1.77E-24 | 1.10284979 | 0.28243536 | -0.2728197 | 0.8853419 |
| ECM1 | -1.2936716 | 0.00032442 | -0.4380306 | 0.63103768 | -0.4257266 | 0.58043085 |
| EDA | -0.9579576 | 0.11217198 | -0.6425735 | 0.64217925 | -0.7166585 | 0.51058362 |
| EDIL3 | -2.1490888 | 0.00095249 | -1.6580993 | 0.07752557 | -1.321513 | 0.19194033 |
| EDN1 | 0.00679207 | 1 | 0.70108235 | 0.20123791 | 0.09466482 | 0.93620623 |
| EDN2 | 0.44623507 | 0.80278414 | 2.10410983 | 0.43911323 | 1.12098661 | 0.72247913 |
| EDN3 | -4.4507753 | 3.58E-22 | -1.5360676 | 0.00213567 | -0.622319 | 0.43291913 |
| EFEMP1 | -1.9321181 | 1.71E-07 | -0.9410006 | 0.09974072 | -0.34155 | 0.70267512 |
| EFEMP2 | -1.0104917 | 0.00013088 | -0.209578 | 0.80346611 | 0.17504768 | 0.81077388 |
| EFNA1 | -0.5716667 | 0.04552846 | -0.4829529 | 0.36330715 | -0.3199037 | 0.13639429 |
| EFNA2 | -1.3596668 | 0.19091414 | -1.1518572 | 0.59807345 | 0.50315364 | 0.82935648 |
| EFNA3 | -2.2961033 | 3.05E-14 | -0.5671429 | 0.33788402 | -0.3306222 | 0.63340401 |
| EFNA4 | -2.1227326 | 8.96E-09 | -0.2785464 | 0.81678951 | -0.420283 | 0.60501889 |
| EFNA5 | 0.76278878 | 0.00043694 | 0.34276752 | 0.47456627 | 0.50198664 | 0.1371001 |
| EFNB1 | 0.32659328 | 0.29812206 | 0.2087342 | 0.81517346 | 0.82001181 | 0.02728013 |
| EFNB2 | -1.3109382 | 1.47E-07 | -0.0025134 | 1 | 0.25005025 | 0.67266999 |
| EFNB3 | 0.72089746 | 0.45718501 | 0.26193776 | 0.95468624 | -1.1886988 | 0.54981837 |
| EGF | -3.6624282 | 6.02E-13 | -0.492941 | 0.73311143 | -1.1958083 | 0.10580957 |
| EGFL8 | -1.2981955 | 0.0097299 | -0.8405121 | 0.3872266 | -1.078555 | 0.15551979 |
| EPGN | 3.91040695 | 2.95E-06 | 2.0208712 | 0.11407328 | 1.53292047 | 0.28608717 |
| EREG | -2.2187358 | 1.34E-13 | -0.4867502 | 0.44553685 | -0.811183 | 0.05866048 |
| F10 | 0.4027275 | 0.82987578 | 0.80712864 | 0.86140235 | 2.38660653 | 0.20932498 |
| F12 | -0.8058749 | 0.18672695 | -0.171571 | 0.95177909 | -0.5360839 | 0.66584691 |
| F13A1 | -1.0566048 | 0.00588622 | -0.2471552 | 0.85208129 | 0.23997248 | 0.82047914 |

| | | | | | | |
|---|---|---|---|---|---|---|
| Cfdp1 | -2.0085922 | 1.14E-13 | -0.3548829 | 0.44592885 | -1.0975287 | 0.00021785 |
| Cgn | -1.2015224 | 3.90E-06 | -0.6953872 | 0.0381079 | -1.0694084 | 0.00017069 |
| Chad | -4.6348244 | 0.01861972 | -1.0735135 | 0.70782932 | -4.6348244 | 0.03914805 |
| Clcf1 | 1.1990406 | 0.00051987 | 0.63166244 | 0.22893715 | 1.38304546 | 0.00019927 |
| Clec11a | -3.0606198 | 2.89E-16 | -1.2305301 | 0.00397035 | -1.0714637 | 0.0098908 |
| Cntf | 0.19842295 | 0.93856401 | 0.67095983 | 0.86980064 | 1.80876702 | 0.37339734 |
| Cntn2 | 0.86053675 | 0.20721655 | -0.8132704 | 0.55536437 | 1.25695659 | 0.08511373 |
| Cntn4 | 1.93419486 | 0.32264291 | 0 | 1 | 2.72648907 | 0.2425829 |
| Col11a1 | -2.8526058 | 4.77E-15 | -1.2285076 | 0.00097893 | -0.5609853 | 0.15851408 |
| Col14a1 | -3.2741479 | 5.70E-18 | 0.75101802 | 0.149556 | -0.6269897 | 0.17291195 |
| Col18a1 | -0.1319992 | 0.5397299 | 0.34190533 | 0.21862797 | 0.16126932 | 0.54140213 |
| Col1a1 | -2.8330983 | 6.96E-10 | -0.7877512 | 0.24778301 | -0.860194 | 0.12032441 |
| Col1a2 | -2.2682563 | 4.98E-07 | -0.8098315 | 0.2266736 | -0.9964134 | 0.06313021 |
| Col2a1 | -0.9752249 | 0.46857473 | -0.5284475 | 0.84512436 | 3.59165181 | 5.12E-07 |
| Col3a1 | -2.0804726 | 6.55E-06 | -0.2597278 | 0.80369787 | -1.0702626 | 0.04926393 |
| Col4a1 | 1.14319367 | 0.00016176 | -0.2846576 | 0.61691976 | -0.6562783 | 0.07279295 |
| Col4a2 | 1.02320019 | 0.00020602 | -0.2384979 | 0.65629743 | -0.6218139 | 0.06038418 |
| Col4a3 | -0.2805565 | 0.60112744 | -0.0837073 | 0.96394619 | -0.535419 | 0.39125918 |
| Col4a4 | -1.3203947 | 0.00060917 | -0.1957709 | 0.82835795 | -0.7366266 | 0.11444398 |
| Col4a5 | -2.085306 | 4.92E-21 | -0.2967991 | 0.41289114 | -1.3799078 | 1.64E-09 |
| Col4a6 | -2.4652717 | 5.29E-13 | -0.0622981 | 0.96202233 | -1.6188693 | 6.64E-06 |
| Col5a1 | -3.4087039 | 3.91E-24 | 0.28344368 | 0.65156716 | -0.8450992 | 0.02627152 |
| Col5a2 | -1.2973955 | 1.36E-05 | -0.069967 | 0.94521394 | -0.6271143 | 0.08506353 |
| Col5a3 | -1.1012118 | 4.35E-05 | 0.79315431 | 0.01656382 | -1.262774 | 1.19E-05 |
| Col6a1 | -1.7882344 | 4.84E-10 | -0.1760425 | 0.78956251 | -0.1655428 | 0.71558952 |
| Col6a2 | -1.8616358 | 6.01E-10 | -0.2030465 | 0.76033005 | -0.038151 | 0.95903799 |
| Col6a3 | -2.4554586 | 4.96E-13 | -0.1453075 | 0.86605517 | -0.5898997 | 0.16440633 |
| Col6a6 | -1.3407204 | 6.57E-09 | -0.1161694 | 0.83987862 | -0.2702693 | 0.39755586 |
| Col8a1 | -0.8557265 | 0.00872064 | 0.65191936 | 0.12839346 | 0.68154308 | 0.06329918 |
| Col9a1 | -2.0692015 | 0.11274582 | -4.8069716 | 0.01708224 | -2.8979558 | 0.07337709 |
| Col9a2 | -1.297133 | 0.19917957 | 0.03735933 | 0.99733604 | -0.691086 | 0.58163235 |
| Col9a2 | -2.3243382 | 1.40E-11 | 0.17908526 | 0.78186291 | -1.2772978 | 0.00020829 |
| Colq | -4.0509042 | 1.50E-12 | -0.6722478 | 0.40002238 | -1.458314 | 0.00972403 |
| Cort | -2.5627909 | 0.21074793 | 0.11797274 | 0.99539312 | -2.5627909 | 0.29536974 |
| Cp | 3.10157785 | 4.37E-32 | -0.2744078 | 0.56628986 | -0.7115858 | 0.01874276 |
| Crp | 6.69326979 | 3.99E-40 | 0.42033385 | 0.81304534 | -2.59862 | 0.06916683 |
| Csf1 | 1.32410764 | 7.82E-15 | 0.7971782 | 2.84E-05 | -0.6361307 | 0.00120394 |
| Csf2 | 7.48016749 | 1.07E-23 | 1.73146444 | 0.21383909 | 1.78237802 | 0.12204746 |
| Csf3 | 12.8421824 | 1.14E-133 | 0.13083512 | 0.9834477 | 2.45552581 | 0.00432191 |
| Ctf1 | -1.0686134 | 0.00285468 | -0.5652024 | 0.29642546 | 0.04818149 | 0.95214309 |
| Cthrc1 | -3.2199438 | 9.75E-17 | -0.3760117 | 0.58532084 | -0.3301442 | 0.5265169 |
| Cubn | -2.3618182 | 3.21E-11 | -1.094468 | 0.00391972 | -1.4131765 | 7.84E-05 |
| Cx3cl1 | 0.77578729 | 0.01630654 | -0.0205144 | 0.99544146 | 0.54000942 | 0.17274203 |
| Cxadr | 10.8890033 | 2.78E-78 | 3.65890492 | 6.74E-13 | 3.37014952 | 6.41E-11 |
| Cxcl10 | 4.85444039 | 2.25E-38 | 3.53404167 | 1.08E-21 | 2.37141 | 1.27E-10 |
| Cxcl11 | 3.4309695 | 1.68E-07 | 2.40323847 | 0.00312555 | 1.50182833 | 0.09949091 |
| Cxcl12 | 0.98250242 | 0.00071136 | -0.4106334 | 0.39166779 | -0.0250362 | 0.97455188 |
| Cxcl13 | 7.51444042 | 4.24E-39 | 0.40841588 | 0.65594607 | 0.54921886 | 0.39236335 |
| Cxcl16 | 1.8418852 | 2.07E-25 | 0.60884426 | 0.0049486 | 0.98604459 | 2.14E-07 |
| Dcn | 1.02528636 | 0.00700498 | 0.20542882 | 0.81153275 | -0.4560833 | 0.35845477 |
| Defa24 | 0.61076871 | 0.00132618 | -0.0250434 | 0.98133679 | 0.89542113 | 6.94E-06 |
| Dhh | -1.2715717 | 8.76E-05 | 0.28304283 | 0.59783872 | 0.02699266 | 0.97369844 |
| Dkk1 | 0 | 1 | 1.88806348 | 0.55536437 | 1.94038195 | 0.42292157 |
| Dkk2 | -1.0320336 | 1.88E-08 | -0.5641574 | 0.01258204 | -0.3386659 | 0.1437707 |
| Dlk1 | -1.7734413 | 0.02453275 | -0.0330548 | 0.99582803 | -0.4191528 | 0.67157124 |
| Dlk2 | 0.85001053 | 0.01143361 | -0.0668011 | 0.95751376 | 0.05311985 | 0.94345546 |
| Dll1 | 0.21215905 | 0.4789575 | -0.0689419 | 0.93422283 | 0.21397699 | 0.57454965 |
| Dll3 | 0.19151577 | 0.92816311 | -2.5421832 | 0.38389965 | 1.05381917 | 0.55160661 |
| Dll4 | 1.11788039 | 6.91E-06 | -0.9523213 | 0.00237022 | -0.3037348 | 0.39701444 |
| Dscam | -0.5611883 | 0.50449521 | 0.50477502 | 0.69073516 | -0.7686382 | 0.46235418 |
| Dusp18 | -1.6311628 | 3.94E-17 | -0.4442956 | 0.09023877 | 0.60107164 | 0.0049903 |
| Ebi3 | 3.44738949 | 3.05E-22 | 1.1884207 | 0.00519349 | -0.0265022 | 0.97830632 |
| Ecm1 | -0.2000365 | 0.47074319 | 0.70677075 | 0.01906064 | 0.19592231 | 0.58032573 |
| Eda | -0.5452916 | 0.04127832 | -0.6309139 | 0.06126421 | -1.0771298 | 0.00012541 |
| Edil3 | -0.8774855 | 0.04105638 | -0.5848548 | 0.3690654 | -1.0551112 | 0.0312116 |
| Edn1 | 2.64842899 | 1.70E-22 | -0.1626761 | 0.80304586 | 0.15610739 | 0.7237363 |
| Edn2 | 1.28533112 | 0.03660217 | -1.0396056 | 0.35785022 | -1.0936974 | 0.22356503 |
| Edn3 | -5.6638194 | 1.87E-13 | -1.4685663 | 0.02529377 | -2.1103432 | 0.00042781 |
| Efemp1 | 0.76969067 | 0.00025792 | 1.11610216 | 4.83E-07 | 0.39905217 | 0.12689372 |
| Efemp2 | 1.01316147 | 2.29E-07 | 0.85744138 | 8.99E-05 | 0.00788825 | 0.9947254 |
| Efna1 | 0.52330221 | 0.01754959 | -0.3294994 | 0.34536746 | 0.13281667 | 0.68727593 |
| Efna2 | -1.5031544 | 9.52E-06 | -0.3149577 | 0.58532084 | 0.23665457 | 0.59027811 |
| Efna3 | -1.2926587 | 0.0001617 | -0.6353044 | 0.20983389 | -0.0812517 | 0.90305687 |
| Efna4 | -1.1366452 | 9.24E-06 | -0.2821889 | 0.54726559 | 0.29254648 | 0.39285304 |
| Efna5 | 1.09197872 | 2.63E-07 | -0.3147799 | 0.38508561 | 0.09323345 | 0.79936719 |
| Efnb1 | -0.9619389 | 2.63E-05 | -0.4034003 | 0.24654507 | 0.29287459 | 0.3311395 |
| Efnb2 | -0.33861 | 0.16940718 | -0.5755504 | 0.05310739 | 0.40203723 | 0.15835405 |
| Efnb3 | -2.2111721 | 1.30E-07 | -0.5904674 | 0.30238044 | -0.4150413 | 0.39912795 |
| Egf | -2.5799589 | 3.18E-13 | -0.3039034 | 0.64153888 | -0.8152402 | 0.04558899 |
| Egfl8 | -1.9261556 | 4.12E-12 | -0.8404315 | 0.00721923 | -0.4346071 | 0.18253153 |
| Epgn | 4.77506337 | 1.01E-30 | 0.22957497 | 0.80016463 | 2.78600657 | 6.87E-12 |
| Ereg | 0.45800441 | 0.11163914 | 0.07480329 | 0.92750113 | 0.67681209 | 0.03227208 |
| F10 | 6.30287066 | 5.22E-37 | 4.26837018 | 5.65E-19 | 2.43928336 | 5.45E-07 |
| F12 | -0.3104692 | 0.8567931 | 0.50706927 | 0.85633626 | 1.3092972 | 0.32020276 |
| F13a1 | -3.0170801 | 4.02E-24 | 0.07563167 | 0.93478232 | -0.836004 | 0.01304955 |

| | | | | | | |
|---|---|---|---|---|---|---|
| F2 | -1.3189896 | 4.23E-05 | -1.0091218 | 0.01921659 | -0.2074016 | 0.81864375 |
| F7 | -0.4990375 | 0.83479961 | -0.1154573 | 1 | -0.6202841 | 0.91463052 |
| F8 | -1.7929008 | 0.00065343 | -0.1137088 | 0.97198012 | -0.6380446 | 0.52276098 |
| FARP2 | -0.4203173 | 0.07299161 | -0.2772179 | 0.59830916 | -0.2243153 | 0.65610492 |
| FAT4 | -1.3315371 | 0.00116377 | -1.0529697 | 0.08084313 | 0.04953791 | 0.98302732 |
| FBLN1 | -0.5530061 | 0.14208373 | -1.6789845 | 1.01E-05 | 0.73292762 | 0.17495326 |
| FBLN2 | -1.1876167 | 0.00094689 | -1.3616973 | 0.00152208 | -0.353613 | 0.67015744 |
| FBN1 | -0.5467914 | 0.08701497 | -1.1896958 | 0.00054112 | 0.21562599 | 0.78596806 |
| FGB | 0.05930591 | 0.99999306 | -0.0770479 | 1 | -1.9118006 | 0.66570245 |
| FGF1 | -1.1093161 | 0.01098065 | -0.1366638 | 0.9492978 | -0.4011449 | 0.68261003 |
| FGF13 | -2.072388 | 2.56E-11 | -0.0596784 | 0.98229072 | -0.6827864 | 0.16743728 |
| FGF14 | 1.97187925 | 0.40068849 | 0 | 1 | 1.82851402 | 0.68910491 |
| FGF16 | -0.6646751 | 0.69755327 | -1.279981 | 0.75069414 | 0.12983478 | 0.98712614 |
| FGF18 | -2.2776022 | 0.00629458 | -0.7547683 | 0.70711703 | -0.1411431 | 0.96036141 |
| FGF5 | -2.7066495 | 0.24440075 | -0.8727371 | 0.9129743 | -2.7066495 | 0.50096044 |
| FGF6 | -3.5941816 | 0.08650753 | -0.4721717 | 0.94739696 | 0.18492485 | 0.97937658 |
| FGG | 0.73138885 | 0.56065514 | 0.8039016 | 0.80639562 | 0.50975999 | 0.8729139 |
| FGL1 | 3.67066482 | 0.09509304 | 6.94946232 | 0.00267906 | 7.13674293 | 0.00290054 |
| FN1 | -0.7657843 | 0.11202017 | -1.8121619 | 0.00048473 | -0.4138696 | 0.68910491 |
| FST | -0.2001073 | 0.73099675 | -0.6122461 | 0.56250984 | -0.0602696 | 0.97985069 |
| GAD1 | 3.1711017 | 0.162717 | 0 | 1 | 0 | 1 |
| GAL | 0.88015973 | 0.18337552 | -0.115735 | 0.98347814 | 0.16216072 | 0.94250341 |
| GAS6 | -1.5374408 | 3.96E-10 | -0.7866987 | 0.02002169 | -0.031497 | 0.98301498 |
| GAST | 0 | 1 | 3.49701282 | 0.40277235 | 0 | 1 |
| GCG | 0.12355057 | 0.97743637 | -3.2127942 | 0.49067051 | -0.1419466 | 0.99521624 |
| GDF5 | 0.06619334 | 0.99571923 | -0.0853871 | 1 | 1.52798594 | 0.69161614 |
| GDF6 | 1.49623773 | 0.44573659 | -2.7051697 | 0.56824858 | -0.0916829 | 0.9957431 |
| GDNF | 0.20937221 | 0.76612924 | -0.0885335 | 0.98905631 | 0.3335089 | 0.82318617 |
| GNAI2 | 0.98845775 | 0.00014764 | -0.0835282 | 0.95392522 | 0.32967368 | 0.54659022 |
| GNAS | -0.7758591 | 0.0038908 | -0.2795069 | 0.7184522 | -0.2068701 | 0.76440757 |
| GNB3 | -0.437367 | 0.40648128 | -0.0862543 | 0.98118499 | 0.28296687 | 0.81716818 |
| GNRH1 | 2.76925403 | 0.23475763 | 0 | 1 | 1.82835006 | 0.68910491 |
| GPHA2 | -0.6103176 | 0.61663295 | -1.7176393 | 0.4396556 | -1.9612253 | 0.29668485 |
| GREM1 | 1.31449568 | 0.00108701 | -0.93778 | 0.1463576 | 0.03848988 | 0.9910417 |
| GRP | -1.684634 | 0.36422335 | -0.1140835 | 1 | 1.2543115 | 0.64724419 |
| GSTP1 | -2.0826425 | 7.35E-13 | -0.1212799 | 0.93129415 | -0.1844608 | 0.82974395 |
| GUCA2A | 0.051955 | 1 | -1.9222603 | 0.73357605 | 2.1073557 | 0.58260398 |
| HAS2 | 3.09286456 | 1.69E-06 | 0.30151925 | 0.9129743 | 1.25923004 | 0.23851472 |
| HDC | 0.585769 | 0.23865899 | -0.780195 | 0.40536828 | 0.7971778 | 0.29840792 |
| HEBP1 | -0.1881223 | 0.49016883 | -0.0833386 | 0.9387498 | 0.22189375 | 0.67587268 |
| HGF | -0.520827 | 0.5828039 | -1.0841503 | 0.56597942 | 1.04205398 | 0.4358163 |
| HMGB1 | 0.00651655 | 1 | 0.06033411 | 0.9783863 | -0.1083102 | 0.90894216 |
| HP | 7.38687961 | 1.06E-32 | 4.44219274 | 2.27E-14 | 3.70362788 | 4.23E-10 |
| HRAS | -0.6487339 | 0.00779198 | 0.03303521 | 0.99684597 | -0.1410899 | 0.83867509 |
| HRG | 1.97187951 | 0.40068849 | 1.83482169 | 0.74490884 | 0 | 1 |
| HSP90AA1 | 0.8852143 | 0.00248079 | -0.0317338 | 1 | 0.25745649 | 0.71407529 |
| HSP90B1 | 1.20593585 | 5.47E-05 | 0.10786672 | 0.94076946 | 0.45663534 | 0.44307435 |
| HSPA1A | -0.3993169 | 0.16702695 | 0.4842017 | 0.32545425 | -0.4685944 | 0.30714297 |
| HSPG2 | -0.6374188 | 0.08385343 | -0.6519341 | 0.30994288 | -0.2393586 | 0.79613204 |
| ICAM1 | 3.07291039 | 4.64E-08 | 1.38783955 | 0.10749701 | 0.90701418 | 0.38471758 |
| ICAM2 | 1.48740178 | 0.00033266 | -0.0628051 | 0.99494542 | 0.46287774 | 0.62521203 |
| ICAM4 | 2.87348144 | 9.89E-09 | 1.57990202 | 0.01895232 | 0.95616048 | 0.26990645 |
| ICAM5 | 0.72314636 | 0.70644613 | -0.6083895 | 0.93648476 | -0.0970101 | 0.9957431 |
| IFNG | 4.65546606 | 0.02380002 | 1.83449262 | 0.74490884 | 0 | 1 |
| IGF1 | -2.4310279 | 2.29E-07 | -2.0126862 | 0.00019805 | -0.0288873 | 0.9957431 |
| IGFBP4 | 0.58805058 | 0.09698358 | 0.15037271 | 0.91716619 | 0.85460537 | 0.07065446 |
| IHH | 1.34858545 | 0.5039064 | -1.9141029 | 0.73311143 | 0.69510222 | 0.90445257 |
| IL10 | 4.79175404 | 8.53E-12 | 2.4870101 | 0.00962939 | -0.2817965 | 0.93041823 |
| IL11 | -1.4096241 | 0.0022001 | 0.08946297 | 0.98033916 | -0.4623068 | 0.64764113 |
| IL12A | 0.00722308 | 1 | -0.1000568 | 1 | -2.7043041 | 0.50096044 |
| IL12B | -0.0671026 | 0.93996154 | -0.8522053 | 0.47599616 | -0.263725 | 0.87304015 |
| IL13 | 1.33322119 | 0.48484713 | -0.0811762 | 1 | 2.94596405 | 0.17449022 |
| IL15 | 0.76333026 | 0.3044611 | -0.5689208 | 0.78621169 | -0.703581 | 0.65571104 |
| IL16 | -1.1356228 | 0.00233195 | -0.440375 | 0.62977048 | -0.2604882 | 0.78796148 |
| IL17A | 0.0216447 | 1 | -0.0869695 | 1 | -1.9217227 | 0.66570245 |
| IL17B | 0.96014419 | 0.27769987 | -0.980883 | 0.65254017 | -0.4531845 | 0.8500279 |
| IL17F | -1.335873 | 0.51559679 | -0.635841 | 0.93343447 | -4.1345781 | 0.22061747 |
| IL18 | 0.52590777 | 0.10175532 | 0.50452604 | 0.40536828 | 0.58331419 | 0.2360175 |
| IL19 | 4.36363891 | 0.0189623 | 0 | 1 | 0 | 1 |
| IL1A | 2.86764329 | 3.98E-21 | 0.51311432 | 0.45302067 | -0.4651926 | 0.47926157 |
| IL1B | 7.32694461 | 8.17E-15 | 5.43055394 | 1.44E-08 | 2.11674612 | 0.0793414 |
| IL1RN | 1.7228196 | 1.08E-07 | 0.22103582 | 0.85381287 | 0.67505476 | 0.22777903 |
| IL2 | 2.34960194 | 0.0046175 | 1.79012057 | 0.20349076 | 1.816442 | 0.16894025 |
| IL20 | 7.30886096 | 6.69E-08 | 0 | 1 | 0 | 1 |
| IL23A | 0.61267483 | 0.74616929 | -0.4126146 | 0.96464689 | 1.50372948 | 0.63133711 |
| IL24 | 7.5045622 | 1.54E-07 | 1.8329991 | 0.74310601 | 0 | 1 |
| IL25 | -2.5616688 | 0.00118116 | 0.04003033 | 1 | -0.511099 | 0.7801868 |
| IL27 | 4.69331578 | 1.02E-05 | 1.51270715 | 0.61401092 | 1.00663752 | 0.75668426 |
| IL34 | -0.7674726 | 0.06486168 | -0.5810711 | 0.50366908 | -1.1830217 | 0.02490954 |
| IL4 | -0.9266847 | 0.47473903 | 0.37293103 | 0.93796486 | -0.9029489 | 0.73625357 |
| IL5 | -0.46178 | 0.86176972 | -1.3755792 | 0.83424825 | -1.4062062 | 0.79023255 |
| IL6 | 12.7613859 | 6.45E-20 | 9.67646401 | 6.52E-11 | 6.16573695 | 0.00196733 |
| IL7 | 2.08059148 | 0.00023773 | 0.67340103 | 0.65869975 | 0.18475515 | 0.92742131 |
| INHA | -2.0631116 | 0.00641694 | 0.11280379 | 0.98395433 | -0.0407666 | 0.9957431 |
| INHBA | -1.239342 | 0.05185775 | -1.8927034 | 0.02122926 | 0.18652371 | 0.92323398 |

| | | | | | | |
|---|---|---|---|---|---|---|
| F2 | -3.954513 | 0.01934967 | 0.96922964 | 0.52201207 | -2.038206 | 0.21529396 |
| F7 | 3.71961202 | 4.41E-12 | 6.75070808 | 3.25E-34 | 3.06057818 | 1.64E-07 |
| F8 | -1.241466 | 0.00389598 | -0.1417594 | 0.88815714 | -0.2103019 | 0.73040049 |
| Farp2 | 0.20724121 | 0.41043724 | -0.1902456 | 0.67377478 | -0.2235705 | 0.477119 |
| Fat4 | -3.1534947 | 4.32E-21 | -0.4733003 | 0.35415198 | -0.7153772 | 0.05757123 |
| Fbln1 | -1.017108 | 1.00E-06 | -0.2567975 | 0.48805049 | -0.0172142 | 0.97556911 |
| Fbln2 | -0.9948116 | 0.00010891 | 0.71849337 | 0.02563792 | 0.07662538 | 0.86955203 |
| Fbn1 | -0.4061566 | 0.34506317 | 1.04101515 | 0.03481631 | -0.4765874 | 0.35513189 |
| Fgb | -4.7762684 | 0.01419955 | -4.7762684 | 0.04870979 | -4.7762684 | 0.03067485 |
| Fgf1 | -1.2404941 | 3.57E-09 | -0.3277741 | 0.33069408 | 0.05953501 | 0.87700148 |
| Fgf12 | -1.7025563 | 9.60E-08 | -0.6792787 | 0.12217249 | -0.397428 | 0.33833553 |
| Fgf14 | -0.6043494 | 0.74999388 | 0.63766672 | 0.83804134 | 1.35670975 | 0.39351727 |
| Fgf16 | 0.22648659 | 0.78841405 | 0.15403059 | 0.94635562 | 0.88595584 | 0.27675662 |
| Fgf18 | -2.1937299 | 4.25E-06 | -0.4150025 | 0.63762424 | -0.6044696 | 0.32119294 |
| Fgf5 | -2.2163586 | 0.00130066 | -1.1025348 | 0.22083546 | -1.1038086 | 0.14386407 |
| Fgf6 | -5.3968951 | 0.00034965 | -1.5575737 | 0.28541266 | -1.0609328 | 0.37960167 |
| Fgg | 2.58669835 | 0.00072127 | -2.3763221 | 0.08740009 | -5.5355532 | 0.00082647 |
| Fgl1 | 2.72688165 | 3.26E-10 | 0.03016422 | 0.99567548 | -0.5122237 | 0.53024707 |
| Fn1 | -1.7471913 | 1.41E-06 | 1.22896935 | 0.00401675 | -0.7487566 | 0.08996619 |
| Fst | 1.15583813 | 3.54E-10 | 0.11406929 | 0.79377007 | -0.3616285 | 0.12679655 |
| Gad1 | 2.39281177 | 0.25112584 | 0.92787354 | 0.8460817 | 2.70911958 | 0.27378284 |
| Gal | -3.731002 | 1.56E-08 | -1.096574 | 0.20675733 | -3.8451884 | 5.02E-08 |
| Gas6 | -0.9642938 | 4.24E-05 | 0.80579082 | 0.00366563 | 0.07744151 | 0.851812 |
| Gast | -3.384327 | 0.00439589 | -1.7592575 | 0.20575049 | -2.0946388 | 0.07896579 |
| Gcg | -3.0356103 | 0.14844676 | 0.48721268 | 0.9096341 | -3.0356103 | 0.22252608 |
| Gdf5 | 3.81982036 | 1.02E-17 | 0.58935252 | 0.59087265 | 0.10382973 | 0.94249568 |
| Gdf6 | -4.3353927 | 0.01237466 | 0.3324997 | 0.90324043 | -1.1081477 | 0.48664208 |
| Gdnf | 1.48504751 | 1.82E-05 | -0.3667876 | 0.59022459 | -0.5203724 | 0.28805547 |
| Gnai2 | 0.48358465 | 0.01633325 | 0.4268137 | 0.11662535 | -0.0064902 | 0.9981106 |
| Gnas | -0.4536178 | 0.04653025 | -0.3482918 | 0.31310838 | -0.4299549 | 0.10772923 |
| Gnb3 | -0.3292438 | 0.65755388 | 0.03284882 | 0.99577013 | -0.7249946 | 0.41457407 |
| Gnrh1 | -0.2661454 | 0.61037786 | 0.67701714 | 0.31230023 | 0.59889774 | 0.28117632 |
| Gpha2 | -3.2333875 | 3.66E-12 | -0.7222225 | 0.25865386 | -3.2341045 | 3.17E-11 |
| Grem1 | 2.40689495 | 9.43E-18 | 0.04328008 | 0.97588194 | -0.3026096 | 0.45033933 |
| Grp | 0.57239853 | 0.72646976 | 0.10843174 | 0.9944048 | 1.7456315 | 0.2670167 |
| Gstp2 | 0.1046736 | 0.6490239 | 0.10728295 | 0.81083639 | 0.05995383 | 0.86180276 |
| Guca2a | -1.5015752 | 0.02026929 | 0.8475281 | 0.36901564 | -1.9902713 | 0.00668542 |
| Has2 | -1.6820967 | 3.00E-07 | 0.37119059 | 0.51495436 | -1.193778 | 0.00090591 |
| Hdc | 4.96651577 | 1.60E-46 | 1.1183273 | 0.00421249 | 1.71470963 | 7.60E-07 |
| Hebp1 | -1.2698701 | 1.24E-10 | 0.08962557 | 0.85435742 | -0.2828886 | 0.26936225 |
| Hgf | -0.0453194 | 0.94065698 | 0.810339 | 0.12466011 | 0.46143986 | 0.36767273 |
| Hmgb1 | -0.1301564 | 0.50346291 | 0.00114478 | 1 | 0.21528804 | 0.33846913 |
| Hp | 2.55634382 | 1.21E-15 | -1.2148609 | 0.00080306 | -1.0210382 | 0.00452034 |
| Hras | -0.1866651 | 0.41716951 | -0.3522398 | 0.26423388 | 0.30356365 | 0.25157489 |
| Hrg | 1.93407793 | 0.32264291 | 1.86757186 | 0.55536437 | 1.91115846 | 0.4247637 |
| Hsp90aa1 | 1.53826808 | 3.29E-13 | 0.17356129 | 0.69245559 | 0.5656047 | 0.02565793 |
| Hsp90b1 | 1.2554165 | 9.99E-07 | 0.12107574 | 0.85221677 | 0.42647004 | 0.19746562 |
| Hspa1a | 1.27499648 | 6.54E-11 | -0.1890846 | 0.62455371 | 0.38453522 | 0.12515914 |
| Hspg2 | 0.18866088 | 0.56885616 | 0.59635607 | 0.1380048 | -0.4226972 | 0.24912783 |
| Icam1 | 2.71628532 | 5.72E-51 | 1.32899773 | 7.79E-13 | 0.49984537 | 0.02067907 |
| Icam2 | 0.93973917 | 0.00128215 | 0.2446744 | 0.67426348 | -0.3650972 | 0.36312032 |
| Icam4 | -1.26188 | 0.24308308 | 0.85517049 | 0.51610708 | 0.69335075 | 0.50644305 |
| Icam5 | -0.9416288 | 0.08201947 | 0.99735815 | 0.07219475 | 1.12964877 | 0.01845364 |
| Ifng | 1.82219772 | 0.00987717 | 0.77015011 | 0.56546331 | 1.2938299 | 0.14136799 |
| Igf1 | -3.1263412 | 1.84E-36 | -0.5199221 | 0.12467044 | -0.9054037 | 0.0006452 |
| Igfbp4 | -1.0967482 | 1.07E-08 | -0.4285148 | 0.11496874 | -0.2203331 | 0.41201625 |
| Ihh | 0.68158469 | 0.5532796 | 0.15304433 | 0.97617458 | 0.19304913 | 0.93012376 |
| Il10 | 3.59372132 | 4.10E-26 | 2.89969217 | 4.53E-16 | 0.12462907 | 0.87700148 |
| Il11 | 1.08528237 | 0.03344321 | 0.35931165 | 0.74489687 | 0.22542472 | 0.79596564 |
| Il12a | 3.31866294 | 4.33E-07 | -0.1989995 | 0.92274918 | -0.2316371 | 0.84643131 |
| Il12b | -0.6536658 | 0.3738381 | 1.94669916 | 0.00205001 | 2.28971022 | 8.60E-05 |
| Il13 | -1.0875257 | 0.28643156 | -0.9155182 | 0.58269753 | 1.96739081 | 0.01620653 |
| Il15 | -1.2722251 | 5.05E-06 | 0.73351515 | 0.02893177 | -0.408521 | 0.24648179 |
| Il16 | 0.13918551 | 0.59084793 | -0.0854216 | 0.88242213 | -0.4075713 | 0.14102605 |
| Il17a | 5.52470224 | 0.00378364 | 0 | 1 | 1.89875497 | 0.43784215 |
| Il17b | 2.29117786 | 0.01309562 | -1.1732889 | 0.67876539 | 3.59781025 | 1.09E-05 |
| Il17f | 4.1495656 | 0.00031034 | 0.62518496 | 0.8617971 | 0.1590075 | 0.9665584 |
| Il18 | -1.4006065 | 1.98E-11 | 0.36601692 | 0.25364274 | 0.87237999 | 0.00010541 |
| Il19 | 8.01876947 | 1.58E-23 | -0.4210004 | 0.86438446 | 6.2634722 | 2.76E-15 |
| Il1a | 2.98875571 | 6.54E-30 | 0.3957661 | 0.36508557 | 1.05161574 | 0.00026853 |
| Il1b | 8.56387882 | 4.12E-59 | 3.05402381 | 2.94E-11 | 2.59713204 | 2.09E-08 |
| Il1rn | 4.60174312 | 9.55E-40 | 1.452899 | 4.08E-05 | 0.52696435 | 0.20083498 |
| Il2 | -4.1574313 | 0.01618845 | -0.9814436 | 0.65438762 | 0.35298846 | 0.83202312 |
| Il20 | -2.6169676 | 0.06406327 | -2.0121195 | 0.3213087 | -0.1756209 | 0.94759759 |
| Il23a | 4.62287817 | 2.91E-07 | 0.40724958 | 0.88803958 | 1.59106765 | 0.21112909 |
| Il24 | 6.87558398 | 5.57E-08 | 0 | 1 | 9.69680393 | 9.28E-17 |
| Il25 | -2.4185158 | 0.09459479 | 0.46155061 | 0.85213596 | 0.02839596 | 1 |
| Il27 | 3.89323241 | 7.06E-12 | 2.30043711 | 0.00407716 | 2.11832803 | 0.00790259 |
| Il34 | -1.1529325 | 3.80E-08 | -0.4057876 | 0.19239355 | -0.4886018 | 0.05360471 |
| Il4 | 0.97586549 | 0.27952616 | -0.1948933 | 0.94903928 | -0.0768205 | 0.97451936 |
| Il5 | -3.0508649 | 0.1117063 | 0.11881961 | 0.99169192 | 1.47004826 | 0.30235557 |
| Il6 | 9.41870408 | 2.35E-84 | 1.41085787 | 0.03616722 | 0.78929793 | 0.28369975 |
| Il7 | -0.8520154 | 0.00568783 | 0.06839601 | 0.94307628 | -0.0534749 | 0.93231959 |
| Inha | -0.3587144 | 0.29185511 | -0.338874 | 0.55031859 | -0.3739079 | 0.36518067 |
| Inhba | 1.94540788 | 4.15E-07 | 0.61144519 | 0.33767893 | 1.38634146 | 0.00131397 |

| | | | | | | |
|---|---|---|---|---|---|---|
| INHBB | 0.16565864 | 0.86024409 | -1.5170829 | 0.19628076 | 0.08245654 | 0.98308557 |
| ITGB3BP | -0.2633151 | 0.55656885 | 0.02469394 | 1 | -0.0149654 | 0.99660832 |
| ITIH2 | -1.9118748 | 0.40648128 | -0.0771854 | 1 | 0.66802022 | 0.91473673 |
| JAG1 | 0.76993542 | 0.00539081 | 0.33361993 | 0.62464949 | 0.34998059 | 0.52730961 |
| JAG2 | -1.1923023 | 3.78E-05 | -0.0639361 | 0.97579779 | 0.32064119 | 0.61470223 |
| KNG1 | -0.7631024 | 0.7285708 | 2.15514721 | 0.36559406 | 1.06592288 | 0.75690224 |
| L1CAM | -1.2744951 | 0.00704071 | -0.7012036 | 0.47352212 | 0.13986189 | 0.92671223 |
| LAMA1 | 0.53664051 | 0.42438236 | -1.3769595 | 0.17193406 | -0.1734109 | 0.93348628 |
| LAMA2 | -1.0587273 | 1.09E-05 | -1.1294424 | 3.67E-05 | -0.1141704 | 0.88269732 |
| LAMA3 | 0.31974756 | 0.25342552 | 0.03349029 | 0.99780159 | 0.51001659 | 0.20962365 |
| LAMA5 | 0.53538774 | 0.08916334 | 0.10424642 | 0.94000914 | 0.32271166 | 0.62152733 |
| LAMB1 | 0.30296138 | 0.27504008 | -0.2583964 | 0.7184522 | 0.3695766 | 0.43809825 |
| LAMB2 | -0.9919002 | 0.00069117 | -0.5834046 | 0.26888675 | -0.2082617 | 0.79344318 |
| LAMB3 | -0.2212543 | 0.49457831 | 0.13591333 | 0.90435318 | 0.24725056 | 0.70142174 |
| LAMC1 | -0.3198971 | 0.34910699 | -0.3619252 | 0.62218113 | 0.38416269 | 0.52075162 |
| LAMC2 | 2.17347542 | 1.54E-16 | -0.1117299 | 0.93129415 | 0.68637886 | 0.08630669 |
| LCN2 | 6.63126155 | 2.09E-29 | 1.18462644 | 0.15929103 | -0.4567462 | 0.72420458 |
| LEFTY2 | -0.1285848 | 0.942682 | -1.2153644 | 0.71837057 | 0.1669999 | 0.97531837 |
| LEP | 0.48434526 | 0.66165435 | -0.2179901 | 0.96933584 | -0.3724063 | 0.89279509 |
| LHB | 2.26626726 | 7.25E-05 | -0.2255128 | 0.94537662 | -0.0261133 | 0.99673641 |
| LIF | 2.81040657 | 6.33E-07 | 1.28886147 | 0.15466213 | 0.32601355 | 0.84233017 |
| LIN7C | -0.0094792 | 0.99086783 | -0.1860972 | 0.77805421 | -0.1517613 | 0.79613204 |
| LIPC | -0.8490802 | 0.47283888 | 0.0655356 | 1 | 0.14968479 | 0.97531837 |
| LIPH | -2.6224798 | 5.15E-09 | 0.11178647 | 0.96729422 | -0.4512638 | 0.64055165 |
| LPL | -1.1126219 | 0.04009931 | 0.27551307 | 0.89472774 | -0.6781019 | 0.51013907 |
| LRP1B | -0.6698013 | 0.56310613 | -2.7887837 | 0.08340282 | -0.8588032 | 0.70345792 |
| LRPAP1 | -0.2064615 | 0.4801918 | -0.3506891 | 0.53971973 | 0.05147575 | 0.9600714 |
| LTA | -1.8726494 | 0.22250722 | -1.0707194 | 0.78181537 | -2.521463 | 0.31596675 |
| LTB | 0.95454856 | 0.07391867 | -0.158194 | 0.95441387 | 0.53253034 | 0.63833511 |
| LTBP1 | 0.97850729 | 1.50E-05 | -0.6796593 | 0.03652869 | 0.42252589 | 0.30321739 |
| LTBP3 | -0.9634295 | 6.15E-06 | -0.8733041 | 0.00054861 | -0.2680441 | 0.55085254 |
| LTF | 5.96370151 | 0.00035397 | 2.61640449 | 0.57088754 | 3.08664726 | 0.39455617 |
| LY86 | -1.9584168 | 0.00079018 | -0.544951 | 0.73067286 | 0.26765846 | 0.86444159 |
| LYPD3 | -0.9854075 | 0.00319236 | -0.010234 | 1 | 0.16569482 | 0.87196454 |
| MATN1 | 2.95799168 | 0.00902952 | 2.68275538 | 0.12686134 | 3.82481908 | 0.00332036 |
| MDK | -0.4189988 | 0.32859703 | 0.55893022 | 0.48595958 | 0.21968089 | 0.83540995 |
| MFAP2 | -2.1113589 | 1.55E-06 | -0.8674315 | 0.26717676 | -0.5244496 | 0.56322641 |
| MFAP5 | -2.0491015 | 2.58E-17 | -2.1669855 | 1.35E-17 | -0.4474903 | 0.31318921 |
| MFGE8 | -0.9493591 | 0.00104721 | -0.2643343 | 0.7550292 | 0.06318124 | 0.95835937 |
| MFNG | -0.8547633 | 0.03690686 | -0.0770627 | 0.98044576 | 0.17219121 | 0.88444734 |
| MIA | 0.87067594 | 0.32510148 | 0.13186184 | 0.98907703 | 0.27370652 | 0.91842029 |
| MMP12 | 0.90787549 | 0.45571101 | -1.7091825 | 0.4692093 | 0.61221349 | 0.83768926 |
| MMP2 | 0.23615647 | 0.4955135 | -1.6246984 | 1.25E-07 | 0.30006433 | 0.65557668 |
| MMP7 | 6.52857656 | 7.49E-06 | 1.83279334 | 0.74273065 | 2.60637378 | 0.50097384 |
| MMP9 | 2.61332925 | 0.001796 | 2.64793111 | 0.01496652 | -0.3094887 | 0.91319386 |
| MST1 | 2.17190107 | 1.85E-06 | 1.36744873 | 0.03458105 | 1.31122622 | 0.04575868 |
| MSTN | -5.2156697 | 2.30E-09 | -0.3159579 | 0.93125047 | -1.4886182 | 0.26291967 |
| MYOC | -1.2676784 | 0.05007574 | -1.8064682 | 0.03279888 | -0.9309006 | 0.42641002 |
| NAMPT | 1.2421331 | 4.22E-06 | 0.26852613 | 0.74490884 | -0.1901724 | 0.80579731 |
| NCAM1 | -1.2030702 | 0.01671302 | -0.9972247 | 0.23163024 | -0.6895888 | 0.47266132 |
| NCAN | -0.1493739 | 0.81538579 | 0.67809052 | 0.50644887 | 0.84362688 | 0.28038594 |
| NDP | -0.2894785 | 0.82792074 | -2.7029062 | 0.16610857 | 0.27983534 | 0.93084587 |
| NGF | -0.5986346 | 0.38069651 | 0.78477423 | 0.53031176 | 1.39982945 | 0.07080961 |
| NID1 | -2.0125753 | 8.28E-13 | -1.1443033 | 0.00076443 | -0.3592796 | 0.54990261 |
| NID2 | -0.0117938 | 0.99733677 | -0.791077 | 0.21119794 | -0.0907223 | 0.94896137 |
| NLGN1 | 0.47134864 | 0.8354922 | -2.7059028 | 0.56597942 | 1.05407572 | 0.81039742 |
| NLGN2 | 0.77747706 | 0.00533707 | 0.46240586 | 0.41939318 | 0.20357641 | 0.78460068 |
| NLGN3 | -1.4117382 | 0.03438003 | -1.8212332 | 0.04506645 | -0.8804667 | 0.46681385 |
| NMB | -1.8552834 | 0.04186671 | 0.05459344 | 1 | 1.19401901 | 0.31299022 |
| NMU | -1.8657944 | 0.14672292 | -1.9906106 | 0.4038403 | -1.1485583 | 0.65084013 |
| NPB | -1.5375488 | 0.01124187 | -0.3158523 | 0.87018116 | 0.77697303 | 0.40837368 |
| NPFF | 0.19488999 | 0.68088141 | 0.36017923 | 0.73797649 | -0.0030663 | 1 |
| NPNT | 0.6799459 | 0.05130992 | -0.2423923 | 0.82531276 | 0.2208197 | 0.80937845 |
| NPPB | -2.7124142 | 0.24364335 | -0.0940151 | 1 | -2.7124142 | 0.50096044 |
| NPPC | -1.4511038 | 0.4095671 | -0.2444345 | 0.98836041 | 0.41942435 | 0.93185823 |
| NPY | -1.5872874 | 0.14264978 | -1.2632215 | 0.56597942 | 0.04422186 | 0.9957431 |
| NRG1 | 0.16281601 | 0.88539825 | 0.10144947 | 0.99869197 | 1.98779914 | 0.07262 |
| NRG2 | -0.9054308 | 0.18941279 | 0.35596199 | 0.86743651 | 0.81421569 | 0.4834697 |
| NRG4 | 2.13336106 | 0.00370234 | 0.27228656 | 0.93850514 | -0.012264 | 1 |
| NRTN | -1.3134682 | 0.03396916 | 0.64577518 | 0.63081279 | 1.59801572 | 0.02360856 |
| NTF3 | 0.54914096 | 0.56680065 | 0.18347166 | 0.97415643 | -0.0109428 | 1 |
| NTN1 | -1.2532894 | 0.0225391 | -1.8050371 | 0.00780747 | -0.3167206 | 0.83005277 |
| NTN3 | -4.3524896 | 0.03208323 | -1.7356546 | 0.65115476 | -2.5411449 | 0.39535 |
| NTN4 | -0.2147051 | 0.73433106 | -0.6005086 | 0.61292638 | 0.21153891 | 0.88827951 |
| NTNG1 | -0.2388907 | 0.74291586 | -0.2730326 | 0.90528264 | 0.24224312 | 0.88706793 |
| NTNG2 | -2.661052 | 3.35E-15 | -1.1447819 | 0.0043777 | -0.2175656 | 0.80383019 |
| NTS | 2.55997294 | 6.88E-05 | -0.7055234 | 0.74490884 | 0.82450763 | 0.55495488 |
| NUCB2 | 0.58018314 | 0.05143055 | 0.07648548 | 0.9655615 | -0.0629578 | 0.95675099 |
| NXPH1 | -3.2192229 | 0.14483498 | -1.3859121 | 0.79683968 | -1.3926398 | 0.74592291 |
| NXPH3 | -1.212238 | 0.13794032 | -0.3086679 | 0.9215809 | -0.7826656 | 0.62429602 |
| OBP2A | -0.9317857 | 0.6991076 | 0.44542027 | 0.96918347 | -0.4715878 | 0.94394032 |
| OLFM2 | -1.7114641 | 0.00264781 | -0.7509692 | 0.5487371 | -0.3818938 | 0.78628628 |
| OMG | 1.18761071 | 0.14473975 | 0.05573421 | 1 | -0.1884946 | 0.95031176 |
| ORM1 | 5.30086771 | 8.34E-07 | 0.84780214 | 0.81517346 | 1.89425687 | 0.32509611 |
| OSM | 5.34578119 | 3.18E-10 | 3.13036441 | 0.001719 | 1.38959819 | 0.3585718 |

| | | | | | | |
|---|---|---|---|---|---|---|
| Inhbb | 1.64807198 | 3.91E-15 | 0.73812914 | 0.00370698 | 0.9591184 | 3.33E-05 |
| Itgb3bp | -0.7943546 | 0.00431962 | -0.0295814 | 0.98696581 | -0.2580439 | 0.48338595 |
| Itih2 | -0.8573867 | 0.08997043 | 1.06839386 | 0.05141084 | -1.3028262 | 0.02416036 |
| Jag1 | 0.65908275 | 0.00035405 | 0.2293655 | 0.47932477 | 0.35835023 | 0.11612741 |
| Jag2 | -0.3164912 | 0.1870656 | -0.2490826 | 0.53937227 | -0.0674596 | 0.86477827 |
| Kng1 | 1.1980104 | 0.37396344 | -3.6913162 | 0.15457903 | -1.7801847 | 0.3944408 |
| L1cam | -1.5254999 | 2.90E-16 | 0.10820337 | 0.79831133 | 0.22099891 | 0.37058932 |
| Lama1 | 1.99208673 | 1.43E-07 | 0.09574037 | 0.94141539 | -2.9784335 | 2.84E-09 |
| Lama2 | -2.4713635 | 2.61E-23 | -0.028472 | 0.98631066 | -0.5430449 | 0.06823873 |
| Lama3 | -0.128452 | 0.60525984 | -0.10745 | 0.83378143 | 0.35671765 | 0.17985348 |
| Lama5 | 0.91453568 | 3.06E-07 | -0.0179528 | 0.9906216 | -0.255439 | 0.28971398 |
| Lamb1 | 0.43969268 | 0.08782922 | 0.08675943 | 0.892986 | -0.185717 | 0.60403286 |
| Lamb2 | -0.533891 | 0.02625801 | 0.34562708 | 0.36157967 | -0.3146663 | 0.29790738 |
| Lamb3 | -0.4455085 | 0.093383 | 0.01168894 | 0.99733604 | 0.214412 | 0.54558167 |
| Lamc1 | -0.0582874 | 0.87627552 | 0.30679768 | 0.53629826 | -0.5567832 | 0.09184445 |
| Lamc2 | 0.63594551 | 0.00659235 | -0.0588487 | 0.93605865 | 1.03515074 | 2.22E-05 |
| Lcn2 | 10.8643318 | 3.59E-39 | 0.20762557 | 0.89682583 | 3.93887168 | 4.12E-09 |
| Lefty1 | 0.15068098 | 0.95468461 | 0.10191507 | 0.99544146 | -1.7835129 | 0.47190341 |
| Lep | -1.0075678 | 0.01046151 | -1.3713136 | 0.00212971 | -1.5631518 | 0.00019527 |
| Lgsn | -0.4246382 | 0.70051827 | 0.30815143 | 0.88507139 | 0.1974611 | 0.90360368 |
| Lif | 3.71498221 | 3.00E-19 | 1.14536268 | 0.02276186 | -0.5511679 | 0.30701024 |
| Lin7c | 0.23858467 | 0.16624306 | 0.14556863 | 0.63090497 | 0.19196263 | 0.37237841 |
| Lipc | 1.26618434 | 0.0608572 | 1.12207897 | 0.25561117 | 0.29991149 | 0.80119276 |
| Liph | 0.76187711 | 2.57E-05 | -0.5696551 | 0.01260028 | 0.26790397 | 0.26413494 |
| Lpl | -2.8394322 | 1.06E-29 | 0.02861154 | 0.98641101 | -0.7154347 | 0.01246251 |
| Lrp1b | 0.15068376 | 0.95468461 | -1.7835108 | 0.58532084 | 0.1562095 | 0.97451936 |
| Lrpap1 | 0.10790231 | 0.57360605 | 0.18936758 | 0.52460249 | -0.2108544 | 0.33380229 |
| Lta | -0.5267023 | 0.51495494 | 0.27021656 | 0.87195523 | 0.5784599 | 0.51605688 |
| Ltb | 1.67567918 | 5.43E-07 | 0.72781434 | 0.13940225 | 1.19661605 | 0.00164582 |
| Ltbp1 | -0.3628969 | 0.02752738 | -0.1765986 | 0.5468037 | -0.101725 | 0.67462669 |
| Ltbp3 | -0.6905159 | 6.77E-06 | -0.306144 | 0.17423869 | -0.2156398 | 0.28926297 |
| Ltf | 8.73289735 | 7.71E-33 | 1.76120533 | 0.32748705 | 3.75061665 | 0.00011877 |
| Ly86 | -3.3022397 | 1.13E-23 | 0.92072463 | 0.01527246 | -0.0206373 | 0.98072672 |
| Lypd3 | -0.0118776 | 0.98305935 | -0.1951071 | 0.72749775 | 1.37040841 | 9.64E-07 |
| Matn1 | 0 | 1 | 7.31737093 | 2.06E-13 | 4.35726513 | 0.01064376 |
| Mdk | -1.6068864 | 0.00077876 | -1.2099547 | 0.04666975 | -1.4600969 | 0.00655822 |
| Mfap2 | -4.7264777 | 7.42E-60 | -0.8873596 | 0.00312745 | -1.2054286 | 1.10E-05 |
| Mfap5 | -1.0262489 | 0.00096704 | 1.15248551 | 0.00096486 | -0.9782565 | 0.00504252 |
| Mfge8 | 1.4693835 | 4.15E-14 | -0.0646127 | 0.90919529 | -0.0381726 | 0.92939564 |
| Mfng | -2.4425872 | 5.31E-26 | -0.330049 | 0.36904757 | -0.6318229 | 0.01532824 |
| Mia | -0.3499844 | 0.7102216 | -0.2561176 | 0.89956869 | -0.0593784 | 0.97556911 |
| Mmp12 | -0.719703 | 0.15000314 | 5.31928736 | 4.37E-27 | 2.44433682 | 1.97E-07 |
| Mmp2 | -0.4166143 | 0.15419211 | -0.0974675 | 0.89185201 | -0.6012527 | 0.0650034 |
| Mmp7 | 4.51295122 | 0.00408622 | -1.7691096 | 0.59159425 | -1.7691096 | 0.47724089 |
| Mmp9 | 2.24793005 | 6.47E-11 | -0.018471 | 0.99586104 | -2.1221496 | 1.71E-08 |
| Mst1 | 0.68431404 | 0.01587763 | 0.14611479 | 0.82860696 | 1.15234184 | 7.46E-05 |
| Mstn | -5.0339288 | 1.08E-23 | -1.3162069 | 0.01763662 | -1.3935205 | 0.00639849 |
| Myoc | -5.6342798 | 1.45E-49 | -0.701473 | 0.10706586 | -0.167747 | 0.73701191 |
| Nampt | 1.92728874 | 6.83E-22 | 0.20739265 | 0.58532084 | 0.27967494 | 0.29829897 |
| Ncam1 | -1.0730796 | 2.36E-05 | -0.3507909 | 0.39229436 | -0.58789 | 0.04818024 |
| Ncan | -1.3076532 | 0.11488589 | -0.3225355 | 0.8529302 | -1.5385288 | 0.11483873 |
| Ndp | -2.6407893 | 2.37E-07 | -0.6874123 | 0.27765176 | 0.08385766 | 0.91350326 |
| Ngf | 2.73360489 | 5.48E-15 | 1.51155129 | 0.0001971 | 0.83377499 | 0.06599908 |
| Nid1 | -0.4383772 | 0.11514377 | 0.69143603 | 0.0378586 | -0.6632913 | 0.02978577 |
| Nid2 | -0.5599529 | 0.05741885 | -0.3211546 | 0.52812164 | -1.0533675 | 0.00065132 |
| Nlgn1 | -0.6361659 | 0.51437461 | -1.1542396 | 0.45457281 | -0.483344 | 0.70989105 |
| Nlgn2 | 0.27678074 | 0.16079812 | 0.04109573 | 0.9490614 | 0.33121111 | 0.14563617 |
| Nlgn3 | -1.9863279 | 5.27E-05 | -0.8489026 | 0.16940663 | -0.5192518 | 0.34645781 |
| Nmb | -0.8095536 | 0.11393402 | -0.5111536 | 0.55567033 | -0.7191344 | 0.24421398 |
| Nmu | 0.16380846 | 0.95426817 | 0.89231969 | 0.83158224 | 2.85464283 | 0.15837367 |
| Npb | -1.8770738 | 0.03547756 | -0.3268224 | 0.84006281 | -0.768231 | 0.4313296 |
| Npff | -0.5445032 | 0.26267621 | -0.0330793 | 0.99544146 | 0.36704218 | 0.5416713 |
| Npnt | -1.2002375 | 2.42E-10 | -0.5758347 | 0.01421109 | -0.5492669 | 0.01295895 |
| Nppb | 3.59078087 | 1.02E-06 | -0.4109136 | 0.85491692 | 4.85345927 | 3.95E-10 |
| Nppc | -0.7173692 | 0.1257765 | -0.9970534 | 0.10471446 | -0.8378537 | 0.12539019 |
| Npy | 3.92762988 | 2.15E-13 | 2.01292904 | 0.00311067 | 0.77576921 | 0.37102732 |
| Nrg1 | 4.20553352 | 1.01E-10 | 2.46560225 | 0.00117262 | 4.42194294 | 9.69E-11 |
| Nrg2 | -2.9493579 | 7.20E-09 | -0.4632746 | 0.52896738 | 1.20495022 | 0.00393174 |
| Nrg4 | -1.5663591 | 1.79E-08 | -0.1450802 | 0.82038519 | 0.87329452 | 0.00341218 |
| Nrtn | 0.42427183 | 0.31009884 | -0.2836042 | 0.71674206 | 0.10054545 | 0.88775634 |
| Ntf3 | -1.3463562 | 4.42E-06 | -0.2094392 | 0.71388829 | -0.4174331 | 0.24742227 |
| Ntn1 | -0.9600045 | 2.75E-05 | -0.0209804 | 0.99229957 | -0.3938187 | 0.17393659 |
| Ntn3 | -2.4379904 | 1.75E-07 | -0.6265114 | 0.30130454 | -0.4996351 | 0.32170269 |
| Ntn4 | 0.6062383 | 0.02232329 | 0.88999916 | 0.00265616 | 0.00379505 | 1 |
| Ntng1 | -7.5250549 | 2.79E-15 | -0.7679081 | 0.21864779 | -0.7537902 | 0.14976181 |
| Ntng2 | 2.39156459 | 4.87E-16 | -0.0217827 | 0.99544146 | -0.2136948 | 0.64707744 |
| Nts | 1.8022969 | 0.00016921 | -0.0334675 | 0.99544146 | -0.0931315 | 0.93931405 |
| Nucb2 | 0.39325167 | 0.16936085 | -0.1142118 | 0.86605517 | 0.48515688 | 0.14098279 |
| Nxph1 | -0.3267613 | 0.60225924 | 0.81410855 | 0.26423388 | -2.477913 | 0.00139019 |
| Nxph3 | -1.3803736 | 0.00298953 | 0.25747431 | 0.7878521 | -0.162281 | 0.82300075 |
| Obp2b | 4.86511576 | 0.0020174 | 0 | 1 | 1.91017207 | 0.42103928 |
| Olfm2 | -0.697434 | 0.02816993 | -0.8158716 | 0.03733122 | -0.4234673 | 0.2855275 |
| Omg | -4.657005 | 0.00300324 | 0.5138738 | 0.77117588 | -1.439946 | 0.27488683 |
| Orm2 | 2.15626315 | 3.20E-16 | 0.10161372 | 0.88543177 | -0.4015536 | 0.24546368 |
| Osm | 6.75669668 | 2.43E-30 | 2.77871378 | 1.86E-05 | 2.35295006 | 0.00043316 |

| | | | | | | |
|---|---|---|---|---|---|---|
| OSTN | -1.4469601 | 0.20689998 | -2.8461331 | 0.08016415 | -0.3468715 | 0.91635875 |
| P4HB | 0.79146799 | 0.00467467 | 0.06785334 | 0.96997365 | 0.31029764 | 0.6117946 |
| PCSK9 | 2.12924048 | 0.00065417 | 0.08095972 | 0.99891223 | 1.01455518 | 0.37625281 |
| PDAP1 | 0.51076487 | 0.03055785 | 0.27227823 | 0.63698496 | 0.25242623 | 0.61533152 |
| PDGFA | -1.1548971 | 0.00054744 | -0.8025867 | 0.11880081 | 0.06135511 | 0.96826012 |
| PDGFB | -0.9520122 | 0.01628591 | -0.1788696 | 0.91227651 | -0.1297403 | 0.91855522 |
| PDGFC | 0.57199174 | 0.02432033 | -0.0039893 | 1 | 0.05964796 | 0.94962188 |
| PDGFD | -2.4635072 | 5.33E-08 | -2.4688103 | 5.36E-07 | -0.2692685 | 0.81716818 |
| PENK | -5.8168566 | 0.00018675 | 0.07691988 | 1 | -0.632213 | 0.7925427 |
| PGF | 1.5983842 | 9.42E-05 | -0.5664818 | 0.5855254 | 0.11006309 | 0.9446751 |
| PIGF | 0.46839531 | 0.19015858 | -0.010656 | 1 | 0.28582876 | 0.71075154 |
| PKM | -0.0055317 | 1 | -0.0711086 | 0.96965257 | -0.0894673 | 0.9291406 |
| PLAT | -1.722657 | 3.01E-08 | -0.6636017 | 0.21395675 | -0.0317943 | 0.98896679 |
| PLAU | 0.07379892 | 0.85261283 | -0.4928527 | 0.3943517 | -0.2781085 | 0.67538028 |
| PLG | 5.60688502 | 0.00111736 | 1.82839968 | 0.74490884 | 3.0863939 | 0.39562847 |
| PLTP | -1.0548046 | 5.32E-05 | -0.2252741 | 0.77805421 | 1.31680484 | 4.69E-06 |
| PODXL | -1.2690794 | 0.00277544 | -1.0622475 | 0.08856296 | -0.1123558 | 0.94046937 |
| PODXL2 | -1.4534304 | 0.0258215 | -0.7216552 | 0.61625631 | -0.6525448 | 0.60955314 |
| POMC | -0.5872919 | 0.31941384 | 0.86634406 | 0.32835023 | 0.07846681 | 0.97531837 |
| PON2 | -1.3281305 | 1.14E-08 | -0.2021072 | 0.78870992 | -0.2895995 | 0.57063479 |
| PPBP | 3.44128497 | 6.29E-05 | 2.56593346 | 0.03531809 | 3.34143574 | 0.00219131 |
| PROC | 2.81118342 | 0.25060249 | 3.12362623 | 0.53326002 | 1.80310094 | 0.70190127 |
| PROK1 | 1.09787009 | 0.37020699 | -2.307901 | 0.46505918 | -0.6640482 | 0.8544098 |
| PROK2 | 7.89841514 | 5.13E-13 | 0.69826477 | 0.93011556 | 0.69246437 | 0.89956595 |
| PROS1 | -0.4785446 | 0.17893736 | -0.0893991 | 0.96418547 | 0.07955308 | 0.94738316 |
| PRSS23 | 1.09977837 | 0.00205919 | -1.4732399 | 0.00059169 | 0.74257524 | 0.19789519 |
| PSAP | -0.8632439 | 0.00441693 | -0.3629759 | 0.63081279 | -0.171564 | 0.84490039 |
| PSEN1 | 1.01908968 | 1.72E-07 | 0.30989252 | 0.49874973 | 0.0857476 | 0.89853149 |
| PSPN | -0.568528 | 0.45607134 | 0.23673361 | 0.93612602 | -0.2280597 | 0.91376876 |
| PTDSS1 | 0.25659451 | 0.37391612 | 0.17220504 | 0.83623455 | 0.25748848 | 0.65342693 |
| PTGS2 | 4.19143791 | 4.99E-08 | 3.31595516 | 0.0001702 | 0.54084133 | 0.78752822 |
| PTHLH | 1.10550195 | 0.13377971 | -1.1836634 | 0.4692093 | -0.3008862 | 0.89492419 |
| PTMA | 0.26372599 | 0.45869866 | 0.22931612 | 0.8144201 | -0.0786907 | 0.9446751 |
| PTN | -1.7438105 | 2.19E-11 | -0.4356746 | 0.4516969 | -1.1523066 | 0.00032277 |
| PYY | -1.3422926 | 0.37592851 | 0.19311237 | 0.98856326 | -0.5052936 | 0.89484908 |
| QDPR | -0.4546814 | 0.07373487 | 0.02407446 | 1 | -0.1458232 | 0.82924692 |
| QRFP | 1.97111137 | 0.40047138 | 1.8347532 | 0.74490884 | 1.82836542 | 0.68910491 |
| RARRES2 | -1.1366451 | 0.00060006 | -0.5006983 | 0.49899561 | -0.5501348 | 0.36594299 |
| RBP4 | -1.7409457 | 0.00102995 | -0.52349 | 0.73311143 | -0.4215495 | 0.74873666 |
| RELN | -0.2238974 | 0.72727529 | -1.8135364 | 0.00375349 | -0.6069066 | 0.54880714 |
| RGMA | -1.3128883 | 2.26E-06 | 0.07231154 | 0.96893425 | -0.0019491 | 1 |
| RGMB | -1.0404752 | 6.01E-05 | -0.3524314 | 0.56639032 | -0.2246098 | 0.7269224 |
| RIMS2 | -2.5457466 | 0.03859629 | -0.3621835 | 0.9401472 | -0.697601 | 0.80595757 |
| RLN3 | 0.04938453 | 1 | -0.0956239 | 1 | -1.9217218 | 0.66570245 |
| RPH3A | -6.0786998 | 8.79E-05 | -1.6593193 | 0.39382375 | 0.25369624 | 0.93248233 |
| RPS19 | -0.3322741 | 0.36104864 | 0.29670216 | 0.74490884 | -0.0220433 | 0.9957431 |
| RSPO1 | -0.260537 | 0.68222954 | -0.1750602 | 0.94054271 | -0.5962404 | 0.56813596 |
| RSPO2 | -2.7065822 | 0.24499161 | -2.7065822 | 0.56597942 | -2.7065822 | 0.50096044 |
| RSPO3 | -1.2438664 | 0.00403933 | -1.0251219 | 0.116624 | -0.3134642 | 0.77529206 |
| RSPO4 | 0.75053342 | 0.58724445 | 0.22744857 | 0.9805664 | 4.84682914 | 0.00012155 |
| RTN4 | -0.8743373 | 0.0046681 | 0.04806196 | 0.99169467 | -0.3825197 | 0.54043498 |
| S100A8 | 7.61138633 | 5.88E-14 | 2.25752623 | 0.07108215 | 0.68568342 | 0.73984448 |
| S100A9 | 8.20049697 | 1.83E-16 | 2.38838964 | 0.03524048 | -0.0709613 | 1 |
| SEMA3A | 1.47228909 | 0.02225943 | -1.3652082 | 0.25186494 | 0.13686556 | 0.95883608 |
| SEMA3B | -1.4324317 | 2.25E-07 | -0.7955095 | 0.04676654 | -0.4016536 | 0.48296191 |
| SEMA3C | -0.9310113 | 7.69E-05 | -0.88572 | 0.00208778 | -0.3423183 | 0.47811861 |
| SEMA3D | -2.07675 | 5.84E-05 | -1.7137887 | 0.00880735 | -1.183885 | 0.1234339 |
| SEMA3E | -2.5938169 | 0.10539972 | -1.283948 | 0.73921296 | -0.3108715 | 0.94726368 |
| SEMA3F | -0.3990523 | 0.11245654 | 0.25150539 | 0.69701072 | 0.47941191 | 0.19506146 |
| SEMA3G | -1.6187663 | 3.84E-05 | -0.1427427 | 0.93850514 | 0.14424943 | 0.91270252 |
| SEMA4A | -0.9203079 | 0.00014948 | 0.13709526 | 0.88825225 | -0.5460793 | 0.1564064 |
| SEMA4B | 2.06350685 | 1.58E-13 | -0.0459311 | 0.991792 | -0.1962077 | 0.80787414 |
| SEMA4C | 0.82640558 | 0.00140462 | 0.25118333 | 0.74353945 | 0.33185585 | 0.52888779 |
| SEMA4D | 0.72114908 | 0.00798241 | -0.072472 | 0.96594578 | 0.03030044 | 0.98649108 |
| SEMA4F | -0.2516921 | 0.46728088 | 0.04290747 | 0.99465564 | 0.01845191 | 0.9957431 |
| SEMA4G | -0.8120098 | 0.02638041 | 0.3388705 | 0.73482109 | 0.77270268 | 0.1426701 |
| SEMA5A | -0.862958 | 0.01009873 | -0.8602111 | 0.07243687 | -1.2095567 | 0.00343332 |
| SEMA6A | 0.49824888 | 0.12704722 | -0.5074174 | 0.42052079 | -0.2070371 | 0.80195123 |
| SEMA6D | 0.02682405 | 0.96931928 | -0.5668784 | 0.42673891 | -0.2729062 | 0.75503655 |
| SEMA7A | 2.00620347 | 1.23E-15 | -0.6508828 | 0.11407328 | 0.08987345 | 0.9197356 |
| SERPINC1 | 0.87273893 | 0.11249448 | -0.3732244 | 0.83241796 | 0.14035144 | 0.94057664 |
| SERPINE1 | 5.17341705 | 3.11E-21 | 0.88467205 | 0.39188252 | 2.30535919 | 0.0001606 |
| SERPINE2 | 0.47636128 | 0.2909601 | -0.4552506 | 0.67028691 | -0.1544606 | 0.90783205 |
| SERPING1 | 1.25168667 | 3.35E-05 | 0.18267279 | 0.87508947 | 0.43336369 | 0.48612459 |
| SFRP1 | 0.13699904 | 0.82241527 | 0.07493304 | 0.98905631 | 0.00567361 | 1 |
| SHANK1 | -3.0431595 | 0.00025825 | -0.1490219 | 0.97441817 | -0.621336 | 0.69161614 |
| SLIT1 | 2.29090281 | 0.18876015 | 1.20024524 | 0.82867669 | 1.19196154 | 0.7925427 |
| SLIT2 | 0.01390332 | 0.99254926 | -0.9518584 | 0.06046592 | -0.3783943 | 0.62888021 |
| SLIT3 | -1.0784762 | 0.00019039 | -1.6356682 | 1.28E-07 | -0.3911704 | 0.50303651 |
| SLPI | 3.25887619 | 4.30E-06 | 0.43228175 | 0.85894295 | -1.9129664 | 0.05179379 |
| SORBS1 | -0.758979 | 0.15417962 | -0.5187457 | 0.70674542 | -0.6538167 | 0.50096044 |
| SOST | 0 | 1 | 0 | 1 | 2.59612631 | 0.52961287 |
| SPINT1 | 0.32033718 | 0.23360025 | 0.0002778 | 1 | 0.11457353 | 0.87842011 |
| SPP1 | -1.666244 | 0.09098091 | -3.4959453 | 0.00417401 | -0.657793 | 0.78161676 |
| SYTL3 | -0.6679841 | 0.17973626 | -0.4189257 | 0.74490884 | 0.25476195 | 0.83768926 |

| | | | | | | |
|---|---|---|---|---|---|---|
| Ostn | -3.2175345 | 2.06E-07 | -1.1125389 | 0.08288818 | -0.7447455 | 0.20396123 |
| P4hb | 1.04152633 | 1.32E-05 | -0.0215234 | 0.99258138 | 0.33369283 | 0.29218553 |
| Pcsk9 | 0.84101317 | 0.05863919 | -0.8152418 | 0.22222 | 0.55789914 | 0.31807338 |
| Pdap1 | 0.25788907 | 0.19785745 | -0.1431867 | 0.70450248 | 0.41434427 | 0.05985609 |
| Pdgfa | 1.67483046 | 1.99E-22 | -0.1015587 | 0.8121687 | -0.1776432 | 0.48471914 |
| Pdgfb | 0.23032515 | 0.31756719 | -0.1445427 | 0.74680516 | -0.5128193 | 0.03818631 |
| Pdgfc | 0.49165648 | 0.05225719 | 0.76724789 | 0.00755157 | -0.1657463 | 0.65172764 |
| Pdgfd | -1.1948042 | 1.17E-06 | -0.0161041 | 0.99544146 | -0.799171 | 0.00399688 |
| Penk | -3.0863873 | 1.14E-32 | -0.5914788 | 0.07221522 | 0.13556398 | 0.72538322 |
| Pgf | 5.88200623 | 6.39E-38 | 1.09381445 | 0.06142003 | 1.08935197 | 0.03799239 |
| Pigf | 0.54996213 | 0.03496325 | 0.40403227 | 0.30863733 | 1.25125118 | 1.40E-06 |
| Pkm | 1.60659725 | 1.77E-08 | 0.12133483 | 0.86980064 | 0.52229353 | 0.14686491 |
| Plat | -0.0332771 | 0.94404149 | 0.9636327 | 0.00633387 | -0.8101981 | 0.02009288 |
| Plau | -1.2563422 | 7.02E-06 | 1.54491206 | 1.22E-07 | 0.02882199 | 0.96887164 |
| Plg | -4.6272978 | 0.01471821 | -2.7407231 | 0.23559247 | -2.717525 | 0.15729019 |
| Pltp | -2.189415 | 1.70E-19 | 0.74892801 | 0.01119201 | -0.6942141 | 0.01359436 |
| Podxl | -1.2365911 | 9.77E-10 | -0.4320168 | 0.13539326 | -0.2149666 | 0.44538915 |
| Podxl2 | -1.9257947 | 2.56E-13 | -0.7369489 | 0.02275393 | -0.9317257 | 0.00123214 |
| Pomc | 0.39818204 | 0.6694827 | -0.7271231 | 0.65987228 | -1.4904553 | 0.20603069 |
| Pon2 | 0.16831452 | 0.42280165 | 0.0952393 | 0.83381799 | 0.11157976 | 0.70180302 |
| Ppbp | 2.21039289 | 0.00064174 | 7.56472825 | 2.17E-25 | -0.7008331 | 0.50008642 |
| Prob1 | -2.533323 | 0.21364123 | -0.6454888 | 0.87958999 | -2.533323 | 0.29777044 |
| Prok1 | 0.21567138 | 0.91177511 | -3.4051731 | 0.20614675 | 0.50620029 | 0.81562102 |
| Prok2 | 8.95009962 | 1.86E-52 | 1.55623731 | 0.40436554 | 0.16501828 | 0.96073648 |
| Pros1 | -0.1588312 | 0.43002375 | 0.55632149 | 0.00958214 | 0.20816971 | 0.39272949 |
| Prss23 | -0.4893502 | 0.03142519 | 0.34682991 | 0.31840941 | -0.5238985 | 0.04353039 |
| Psap | -0.1796811 | 0.5532497 | 1.18098092 | 3.22E-05 | 0.08699934 | 0.84870302 |
| Psen1 | 0.54611419 | 0.00041048 | 0.04483461 | 0.92393334 | 0.10665077 | 0.647784 |
| Pspn | -1.1048051 | 0.51977655 | -1.1716077 | 0.69676331 | 1.07912014 | 0.49641277 |
| Ptdss1 | -0.0442302 | 0.82879421 | -0.1135215 | 0.72975291 | 0.0486646 | 0.86088213 |
| Ptgs2 | 6.81026609 | 1.23E-34 | 2.16566877 | 4.87E-05 | 0.72475273 | 0.25165574 |
| Pthlh | -0.4257039 | 0.35611887 | 0.64445102 | 0.30838781 | -0.1400425 | 0.84261753 |
| Ptma | 0.28739919 | 0.21288249 | -0.0032662 | 1 | 0.33944882 | 0.20827321 |
| Ptn | -4.3335848 | 5.29E-82 | -0.173891 | 0.66425667 | -0.4962466 | 0.04044363 |
| Pyy | -0.034451 | 0.98455217 | 0.56167375 | 0.73259755 | 4.73930545 | 1.30E-09 |
| Qdpr | -0.5727024 | 0.00040778 | -0.159092 | 0.59303677 | 0.12638228 | 0.58913888 |
| Qrfp | -1.5435816 | 0.19068854 | 0.25616695 | 0.924065 | -0.4682258 | 0.76222336 |
| Rarres2 | 2.31949602 | 7.21E-16 | -0.0916988 | 0.91017438 | -0.1837154 | 0.67852527 |
| Rbp4 | -0.5078324 | 0.08276289 | -0.3913962 | 0.39257065 | -0.5568231 | 0.09998929 |
| Reln | 0.01659769 | 0.96507696 | -0.1218786 | 0.82495405 | -0.9244676 | 0.00027772 |
| Rgma | -0.7387566 | 0.00016882 | -0.4536773 | 0.08812673 | 0.00487106 | 1 |
| Rgmb | -1.2537868 | 7.75E-09 | -0.2859768 | 0.44535116 | -0.1460184 | 0.66002219 |
| Rims2 | 0.22013613 | 0.81765729 | 0.14494791 | 0.95943841 | 1.46026467 | 0.05256427 |
| Rln3 | -3.9452508 | 0.02615822 | -1.2694767 | 0.5790536 | -0.0529257 | 0.98938345 |
| Rph3a | 1.0689138 | 0.48174417 | 2.21642035 | 0.21601476 | -2.5576104 | 0.28537072 |
| Rps19 | 0.72923633 | 0.00060622 | 0.32435541 | 0.34681421 | 0.00876485 | 0.99362448 |
| Rspo1 | -1.2637728 | 1.12E-06 | -0.2860284 | 0.54425579 | -1.2668699 | 5.34E-06 |
| Rspo2 | -3.1824497 | 5.34E-05 | -0.1657423 | 0.92949934 | -1.4967114 | 0.06234169 |
| Rspo3 | -0.6944534 | 0.05459616 | 0.40910489 | 0.47210337 | -0.9019352 | 0.02654321 |
| Rspo4 | -2.2891208 | 3.04E-07 | -1.185146 | 0.01776743 | -0.0571456 | 0.9473847 |
| Rtn4 | -0.3896316 | 0.09997271 | -0.0003951 | 1 | -0.2661783 | 0.37539989 |
| S100a8 | 10.7741561 | 1.03E-27 | 3.30771665 | 7.50E-05 | 8.16852529 | 1.06E-18 |
| Gm5849 | 10.3333156 | 5.82E-27 | 2.84118278 | 0.00061702 | 7.42245024 | 7.09E-17 |
| Sema3a | 1.68434079 | 1.72E-23 | -0.2016486 | 0.55286318 | 0.10225772 | 0.72033276 |
| Sema3b | -2.7018227 | 4.38E-24 | -0.7360416 | 0.02631042 | -1.2208238 | 1.49E-05 |
| Sema3c | -0.9204515 | 1.28E-06 | 0.03284747 | 0.9681616 | -0.7256927 | 0.00058367 |
| Sema3d | -3.069011 | 1.70E-17 | -0.0638669 | 0.96412032 | -1.4826177 | 9.86E-05 |
| Sema3e | -0.884244 | 0.00570174 | -0.2580079 | 0.67741092 | 0.31021346 | 0.47190341 |
| Sema3f | 0.90658907 | 3.44E-07 | -0.3165447 | 0.2608965 | 0.21780078 | 0.37769291 |
| Sema3g | -0.497256 | 0.07654294 | -0.7842134 | 0.01685746 | -0.1903 | 0.62687034 |
| Sema4a | 0.08952851 | 0.63748989 | 0.04862864 | 0.9162928 | 0.33314741 | 0.08553166 |
| Sema4b | 0.12083884 | 0.5570081 | -0.0837406 | 0.84938784 | 0.1716335 | 0.48170859 |
| Sema4c | 1.25115993 | 4.91E-11 | 0.00237881 | 1 | 0.39248816 | 0.10672302 |
| Sema4d | 1.94432062 | 2.01E-13 | 0.75854468 | 0.02533639 | 0.62328333 | 0.05459588 |
| Sema4f | -1.2466556 | 0.00067862 | 0.15239423 | 0.85213596 | 0.13699883 | 0.80379337 |
| Sema4g | -2.3551981 | 6.59E-18 | -0.3860162 | 0.36655073 | 0.17950062 | 0.64487515 |
| Sema5a | 0.37015585 | 0.05806288 | -0.5980808 | 0.00754325 | -0.1533588 | 0.56988456 |
| Sema6a | -0.3825234 | 0.11260065 | -0.8638292 | 0.00082432 | -0.2983806 | 0.31297702 |
| Sema6d | 1.39849535 | 1.18E-13 | 0.12013444 | 0.78925908 | -0.2670157 | 0.30570362 |
| Sema7a | 1.60189221 | 4.09E-22 | -0.3360786 | 0.2056405 | -0.0364409 | 0.92105544 |
| Serpinc1 | -1.6471159 | 0.21496253 | -0.8228959 | 0.72681158 | -0.5000239 | 0.7851265 |
| Serpine1 | 6.74694397 | 2.64E-29 | 2.06282079 | 0.00034967 | 0.1933521 | 0.82114015 |
| Serpine2 | -0.2594154 | 0.15911893 | 1.00418105 | 7.18E-09 | -0.2451343 | 0.269252 |
| Serping1 | 1.11822586 | 0.00043356 | 0.51452818 | 0.30076672 | -0.1073317 | 0.84479632 |
| Sfrp1 | -0.7921096 | 0.00080113 | -0.4265043 | 0.22684126 | 0.82567969 | 0.00135844 |
| Shank1 | -1.4361904 | 2.31E-05 | -0.3181277 | 0.58532084 | -0.6757621 | 0.08604053 |
| Slit1 | 1.347228 | 0.00040568 | -0.4949621 | 0.49372885 | 0.65464859 | 0.18277152 |
| Slit2 | -1.4562118 | 4.34E-09 | 0.12967658 | 0.82673128 | -0.2403735 | 0.48253147 |
| Slit3 | -0.1414731 | 0.55649178 | 0.60844193 | 0.01632163 | -0.3585075 | 0.1698296 |
| Slpi | 8.99130112 | 7.33E-48 | 2.68705357 | 4.39E-06 | 5.98776889 | 1.52E-25 |
| Sorbs1 | -0.8790844 | 9.49E-06 | -0.3990583 | 0.16985677 | -0.3741072 | 0.13294042 |
| Sost | 0 | 1 | 2.66751158 | 0.3415101 | 4.37299125 | 0.01534773 |
| Spint1 | 0.55348248 | 0.02927421 | 0.35906512 | 0.36904757 | 0.83507379 | 0.00208972 |
| Spp1 | 3.16502849 | 1.86E-18 | 2.90572207 | 5.28E-15 | -0.034667 | 0.97253456 |
| Sytl3 | 1.27530526 | 0.00025461 | 0.0849929 | 0.9490614 | 0.33256964 | 0.52615939 |

| | | | | | | |
|---|---|---|---|---|---|---|
| TAC4 | -0.7447793 | 0.10617887 | -0.3755422 | 0.76500859 | -0.5848189 | 0.49657272 |
| TCN2 | -0.8296532 | 0.00038802 | -0.541173 | 0.14890024 | 0.09506233 | 0.90167883 |
| TCTN1 | -1.0791047 | 0.00553612 | -0.0641996 | 0.98792538 | -0.209775 | 0.84546682 |
| TFPI | 0.802441 | 0.02953483 | -0.5503201 | 0.48726357 | -0.0435171 | 0.98344503 |
| TG | -0.0636086 | 0.97894377 | 0.57439293 | 0.88067092 | -1.4931056 | 0.50824948 |
| TGFA | 0.41158014 | 0.29620682 | -0.3508365 | 0.73311143 | 0.51199197 | 0.45082966 |
| TGFB1 | -0.2262334 | 0.60500434 | 0.08643785 | 0.96976028 | 0.46317326 | 0.50917262 |
| TGFB2 | -2.1541767 | 7.46E-14 | -0.1830891 | 0.86026061 | 0.30195961 | 0.65089806 |
| TGFB3 | -0.4878208 | 0.23566229 | -0.2378876 | 0.85263649 | 0.34029489 | 0.68910491 |
| TGM2 | 1.47383724 | 2.11E-07 | 0.18311901 | 0.86335005 | -0.3835658 | 0.52520099 |
| THBS1 | 3.71689495 | 2.18E-09 | 0.66845028 | 0.68357677 | 0.63310246 | 0.64282561 |
| THBS2 | -0.5072194 | 0.17579424 | -0.4682313 | 0.56077739 | 0.46183257 | 0.50096044 |
| TIMP1 | 6.17012922 | 3.83E-24 | 2.36088536 | 0.00020143 | 2.06000986 | 0.00305346 |
| TIMP2 | -0.7093635 | 0.02424589 | -1.6237532 | 4.96E-07 | 0.19783748 | 0.81340003 |
| TIMP3 | -1.3807838 | 5.13E-06 | -0.6165211 | 0.26711857 | -0.0798544 | 0.94480091 |
| TNC | 2.13119399 | 1.98E-08 | -1.6115044 | 0.00036429 | -0.0181234 | 0.9957431 |
| TNF | 2.62163778 | 0.00038237 | 1.57345426 | 0.19820825 | -0.2714109 | 0.91333037 |
| TNFSF13 | 0.73934311 | 0.20856416 | 0.50216524 | 0.74490884 | -0.3444907 | 0.82357824 |
| TNFSF13B | 1.31948548 | 0.08106863 | -0.7439694 | 0.74490884 | 0.92241708 | 0.52276098 |
| TNFSF14 | 0.56362828 | 0.38618505 | 0.00340957 | 1 | -0.4496267 | 0.76105796 |
| TNFSF15 | 0.32717007 | 0.78420256 | -0.0870245 | 1 | -0.6927849 | 0.7681879 |
| TNFSF18 | -1.6517349 | 0.02664902 | 0.17747417 | 0.96610337 | 3.48098065 | 4.54E-06 |
| TPH1 | 0.61356803 | 0.43198943 | 1.124392 | 0.37284917 | -0.2854901 | 0.90439053 |
| TSLP | -1.0775421 | 0.36603642 | 0.40412987 | 0.91109076 | 0.68167963 | 0.73984448 |
| UBA52 | -0.5338954 | 0.07999201 | -0.1147259 | 0.93158309 | -0.2129602 | 0.77639921 |
| UCN2 | -0.8940143 | 0.26970984 | -0.4380992 | 0.8562534 | 0.25408401 | 0.90599008 |
| VASP | 0.91442509 | 0.00181771 | 0.14007955 | 0.91163333 | 0.19360218 | 0.8122439 |
| VCAM1 | -0.1835833 | 0.74796032 | -1.8272071 | 0.00036429 | -0.5251094 | 0.56275408 |
| VCAN | 2.08617681 | 4.71E-05 | 0.2062373 | 0.93169409 | 0.52798184 | 0.65528066 |
| VIM | -1.4008349 | 0.00012913 | -0.5514491 | 0.50534743 | -0.0851932 | 0.95402473 |
| VIP | 0.78946849 | 0.60387674 | 2.17133255 | 0.28968877 | 0.96249462 | 0.75184315 |
| VTN | 0.31886903 | 0.75417904 | 1.24620912 | 0.37426581 | 1.22549469 | 0.33257522 |
| VWF | -1.6314676 | 1.40E-09 | 0.10853395 | 0.93576433 | -0.0865456 | 0.93022216 |
| WNT1 | 3.59571348 | 0.08614027 | 3.12137621 | 0.45922981 | 0 | 1 |
| WNT11 | -1.763195 | 0.00033093 | 0.21708653 | 0.91335755 | 0.57222478 | 0.54733415 |
| WNT16 | -0.418102 | 0.26978892 | 0.19989456 | 0.87008106 | 0.05245668 | 0.97705763 |
| WNT2 | -2.8401148 | 7.10E-05 | -0.8547628 | 0.54535998 | 0.25643944 | 0.88706793 |
| WNT3 | -0.5350639 | 0.19944738 | -0.006541 | 1 | 0.61914468 | 0.35477649 |
| WNT3A | -1.0611232 | 0.10821919 | 0.38808352 | 0.82867669 | 0.39831454 | 0.78242832 |
| WNT4 | -3.4095104 | 3.58E-09 | -0.3372933 | 0.85304366 | -0.1014982 | 0.96499376 |
| WNT5A | -1.1841166 | 0.02091827 | -0.718827 | 0.51257591 | -0.4266444 | 0.7149693 |
| WNT7A | -1.91328 | 0.40648128 | -1.91328 | 0.73311143 | 0.69964724 | 0.91261588 |
| WNT7B | -1.1258864 | 0.00846695 | -0.8667871 | 0.2239536 | -0.0456952 | 0.98649627 |
| YARS | 0.9866768 | 4.95E-05 | 0.15740432 | 0.8612652 | 0.23833268 | 0.68910491 |
| ZP3 | -3.2150463 | 0.16232413 | -0.6045096 | 0.94076946 | -1.3865263 | 0.7702463 |

| | | | | | | |
|---|---|---|---|---|---|---|
| Tac4 | -3.0791316 | 8.68E-05 | -0.9147009 | 0.36331378 | 1.52242025 | 0.01385684 |
| Tcn2 | -0.8081696 | 1.57E-05 | 0.18014563 | 0.60851553 | -0.276138 | 0.25814352 |
| Tctn1 | -0.2022462 | 0.38909217 | 0.29774244 | 0.38503983 | -0.0204124 | 0.97143499 |
| Tfpi | 0.59599083 | 0.01259913 | 0.15257011 | 0.76977094 | -0.6106401 | 0.02718553 |
| Tg | 2.30194454 | 0.00261557 | -0.3012229 | 0.90294968 | -1.1999174 | 0.37482347 |
| Tgfa | 1.32039937 | 8.25E-10 | -0.2545205 | 0.53640249 | 1.11635092 | 1.38E-06 |
| Tgfb1 | 0.54556029 | 0.00609975 | 1.06885211 | 1.16E-07 | 0.45886004 | 0.04884594 |
| Tgfb2 | -1.9650697 | 7.65E-21 | -0.7667124 | 0.00152732 | -1.2682799 | 6.93E-09 |
| Tgfb3 | -1.2938383 | 1.97E-06 | -0.3781114 | 0.40436554 | -0.1077703 | 0.8141279 |
| Tgm2 | 1.89507048 | 1.09E-20 | -0.205449 | 0.5947293 | -0.7889476 | 0.00055354 |
| Thbs1 | 5.01144199 | 1.11E-29 | 1.60540221 | 0.00041601 | 1.0733905 | 0.0220746 |
| Thbs2 | -0.3074585 | 0.27779521 | 0.47603047 | 0.21416698 | 0.06851204 | 0.8879909 |
| Timp1 | 5.56849991 | 1.03E-26 | 3.32855041 | 2.65E-11 | 3.60819296 | 6.65E-13 |
| Timp2 | -2.0457076 | 6.60E-16 | -0.3365977 | 0.4392093 | -0.7550758 | 0.01011207 |
| Timp3 | -1.6256148 | 9.90E-14 | -0.284637 | 0.46043286 | -0.5573681 | 0.03371376 |
| Tnc | 3.64881185 | 3.64E-18 | 1.03817429 | 0.04847114 | 3.28708379 | 2.10E-14 |
| Tnf | 3.90480202 | 3.14E-18 | 1.79616994 | 0.00028643 | -0.1657201 | 0.83372508 |
| Tnfsf13 | -0.4802465 | 0.35897013 | -0.2239351 | 0.84040829 | 0.18445989 | 0.80245769 |
| Tnfsf13b | 0.00848113 | 0.9961278 | 0.23907277 | 0.72493781 | 0.0427015 | 0.95725913 |
| Tnfsf14 | 2.06602372 | 6.29E-05 | 0.70055253 | 0.42971122 | 0.88013 | 0.18612452 |
| Tnfsf15 | -0.7069122 | 0.093383 | -0.1231368 | 0.90824105 | -0.4766354 | 0.3628712 |
| Tnfsf18 | -0.9227788 | 0.05061718 | 1.11956463 | 0.01895183 | -3.3413123 | 6.90E-08 |
| Tph1 | 4.03523676 | 1.87E-28 | 1.05098326 | 0.05686194 | 0.87357149 | 0.09574038 |
| Tslp | -0.5111817 | 0.27466946 | -0.1951587 | 0.8529302 | -0.7000933 | 0.19744917 |
| Gm11808 | 0.33426421 | 0.10807152 | 0.07666526 | 0.87958999 | 0.05417929 | 0.88133821 |
| Ucn2 | -1.9476039 | 0.00029984 | -0.0185526 | 0.99973987 | -0.0251724 | 0.98703844 |
| Vasp | 2.03737662 | 6.37E-22 | 0.3831648 | 0.24616854 | 0.6860826 | 0.00510806 |
| Vcam1 | 1.28868635 | 4.05E-07 | 1.31578754 | 1.49E-06 | 1.23346834 | 6.63E-06 |
| Vcan | 1.52731439 | 6.47E-10 | 1.41970114 | 7.40E-08 | 0.16902426 | 0.65802689 |
| Vim | -0.0719293 | 0.84821184 | 0.58528393 | 0.13874208 | 0.14015801 | 0.75400219 |
| Vip | -3.0503762 | 0.11052211 | -0.3838527 | 0.92393334 | -3.0503762 | 0.17352435 |
| Vtn | -1.385505 | 0.00121097 | -0.1142789 | 0.92743303 | -1.0876155 | 0.02710722 |
| Vwf | -1.7202088 | 1.11E-11 | -0.0352058 | 0.97883568 | -0.135987 | 0.73580964 |
| Wnt1 | 2.73852671 | 0.16006983 | 0 | 1 | 2.70163642 | 0.24433395 |
| Wnt11 | -1.2967539 | 1.03E-05 | 0.33095562 | 0.50343892 | -0.4128888 | 0.26413105 |
| Wnt16 | 2.65239749 | 1.02E-09 | -0.1569633 | 0.89473342 | -0.4364769 | 0.48170859 |
| Wnt2 | -2.0549521 | 2.18E-05 | 1.94103569 | 0.00012686 | -1.976171 | 0.00016864 |
| Wnt3 | -1.2453995 | 4.50E-05 | -0.0821018 | 0.92849824 | 0.0965756 | 0.85449316 |
| Wnt3a | -2.7101037 | 3.48E-09 | 0.14654427 | 0.88823644 | 0.5795374 | 0.25469632 |
| Wnt4 | 0.7338918 | 0.00580024 | -0.416738 | 0.32022962 | 0.86642182 | 0.00304286 |
| Wnt5a | -1.1267144 | 3.42E-08 | -0.0656751 | 0.90704158 | -0.2327015 | 0.4016115 |
| Wnt7a | -1.4102306 | 0.19319663 | -0.16987 | 0.95679834 | 2.09339338 | 0.00534042 |
| Wnt7b | -0.9438562 | 0.00282019 | -0.3673897 | 0.50793122 | -0.2653668 | 0.54796943 |
| Yap1 | 0.88658507 | 4.32E-07 | -0.1644462 | 0.63424309 | 0.226353 | 0.34594987 |
| Zp3 | 0.70375861 | 0.65637155 | -2.5427215 | 0.38739077 | -2.5427215 | 0.27888372 |

**Dataset S3: Summary of enrichment scores determined by xCell.**

| | NMR_CT1 | NMR_CT2 | NMR_CT3 | NMR_LPS1 | NMR_LPS2 | NMR_LPS3 | NMR_3MC1 | NMR_3MC2 | NMR_3MC3 | NMR_UV1 | NMR_UV2 | NMR_UV3 |
|---|---|---|---|---|---|---|---|---|---|---|---|---|
| Adipocytes | 0 | 0.0025 | 0 | 0 | 0 | 0 | 0 | 0 | 0 | 0.0025 | 0 | 0 |
| Astrocytes | 0 | 0 | 0 | 0 | 0 | 0 | 0 | 0 | 0 | 0 | 0 | 0 |
| B-cells | 0 | 0 | 0 | 0 | 0 | 0 | 0 | 0 | 0.0021 | 0 | 0 | 0 |
| Basophils | 0.014 | 0 | 0 | 0.0751 | 0.1505 | 0.0806 | 0.0303 | 0 | 0.005 | 0 | 0 | 0 |
| CD4+ T-cells | 0 | 0 | 0 | 0 | 0 | 0 | 0 | 0 | 0 | 0 | 0 | 0 |
| CD4+ Tcm | 0 | 0 | 0 | 0 | 0 | 0 | 0 | 1.00E-04 | 0 | 0 | 0 | 0 |
| CD4+ Tem | 0 | 0 | 0 | 0.0048 | 0 | 0 | 0 | 0 | 0.0047 | 0 | 0 | 0 |
| CD4+ memory T-cells | 0 | 0 | 0 | 0 | 0 | 0 | 0 | 0 | 0 | 0 | 0 | 0 |
| CD4+ naive T-cells | 0 | 0 | 0 | 0 | 0 | 0 | 0 | 0 | 0 | 0 | 0 | 0 |
| CD8+ T-cells | 0 | 0 | 0 | 0 | 0 | 0 | 0 | 0 | 0 | 0 | 0 | 0 |
| CD8+ Tcm | 0 | 0 | 0 | 0 | 0 | 0 | 0 | 0 | 8.00E-04 | 0 | 0 | 0 |
| CD8+ Tem | 0 | 0 | 0 | 0 | 0 | 0 | 0 | 0 | 0 | 0 | 0 | 0 |
| CD8+ naive T-cells | 0 | 0 | 0 | 0 | 0 | 0 | 0 | 0 | 4.00E-04 | 0 | 0 | 0 |
| CLP | 0.0015 | 0.0028 | 0 | 0 | 0 | 0.0054 | 0.003 | 0.0012 | 0.0015 | 0 | 0 | 0.0012 |
| CMP | 0 | 0.0019 | 0.0018 | 0 | 0 | 0 | 0 | 0 | 0 | 0.0024 | 0.0034 | 0.0016 |
| Chondrocytes | 0 | 0 | 0 | 0 | 0 | 0 | 0 | 0 | 0 | 0 | 0 | 0 |
| Class-switched memory B-cells | 0 | 0 | 0 | 1.00E-04 | 0 | 0 | 0 | 0 | 0.0031 | 0 | 0 | 0 |
| DC | 0 | 0 | 0 | 0.0227 | 0.0108 | 0.0152 | 0 | 0 | 0.0018 | 0 | 0 | 0 |
| Endothelial cells | 0 | 0 | 0.001 | 0 | 0.0061 | 0.005 | 0 | 0 | 0 | 0 | 0.0043 | 0 |
| Eosinophils | 0.0014 | 0 | 0 | 0 | 0 | 0 | 0 | 0 | 6.00E-04 | 0 | 0 | 0 |
| Epithelial cells | 4.00E-04 | 0 | 0 | 0 | 0 | 0 | 0 | 0 | 3.00E-04 | 0 | 0 | 5.00E-04 |
| Erythrocytes | 0 | 0 | 0 | 0 | 0 | 0 | 0 | 0 | 0 | 0 | 0 | 0 |
| Fibroblasts | 0 | 0.0058 | 0.0021 | 0.0123 | 0.0718 | 0 | 0 | 0 | 0 | 0.0149 | 0.035 | 2.00E-04 |
| GMP | 0 | 0.0026 | 0.0137 | 0 | 0 | 0 | 0 | 0 | 0 | 0.0121 | 0.0123 | 0 |
| HSC | 0.0516 | 0.1384 | 0.147 | 0 | 0.0103 | 0.0086 | 0.0065 | 0.0176 | 0.0091 | 0.0886 | 0.1131 | 0.0388 |
| Hepatocytes | 0 | 0 | 0 | 0.0021 | 0.0041 | 0.0023 | 0 | 0 | 0 | 0 | 0 | 0 |
| Keratinocytes | 0 | 0 | 0 | 0 | 0 | 0 | 0 | 0 | 0 | 0 | 1.00E-04 | 5.00E-04 |
| MEP | 0.0046 | 0.0012 | 0 | 0 | 0 | 0 | 0.0025 | 0.0068 | 0 | 0.0017 | 0.006 | 0.012 |
| MPP | 0 | 0 | 0 | 0 | 0 | 0 | 0 | 0 | 0 | 0 | 0 | 0 |
| MSC | 0.0785 | 0.1532 | 0.083 | 0 | 0.0016 | 0 | 0.0045 | 0.077 | 0.0084 | 0.0676 | 0.0633 | 0.0569 |
| Macrophages | 1.00E-04 | 0 | 0 | 1.00E-04 | 0 | 0 | 0 | 0 | 0.0079 | 0.0049 | 0.0072 | 3.00E-04 |
| Macrophages M1 | 0 | 0 | 0 | 0.0175 | 0.0135 | 0.0079 | 0 | 0 | 0.0102 | 5.00E-04 | 0.0054 | 0 |
| Macrophages M2 | 0.0109 | 0.0042 | 0.0063 | 0 | 0 | 0 | 0.0062 | 0.0088 | 5.00E-04 | 0.0145 | 0.0146 | 0.0052 |
| Mast cells | 0 | 0 | 0 | 0.0038 | 0.0046 | 0 | 0 | 0 | 0 | 0 | 0 | 0 |
| Megakaryocytes | 0 | 0 | 0 | 0 | 0 | 0 | 0 | 0 | 0 | 1.00E-04 | 0.0015 | 0 |
| Melanocytes | 4.00E-04 | 9.00E-04 | 0.0018 | 0 | 7.00E-04 | 0 | 0 | 0 | 0 | 0.0033 | 0 | 1.00E-04 |
| Memory B-cells | 0 | 0 | 0 | 0 | 0 | 0 | 0 | 0 | 0 | 0 | 0 | 0 |
| Mesangial cells | 0 | 0 | 0 | 0 | 0.0037 | 0 | 0 | 0 | 0 | 0.0038 | 0.0026 | 0 |
| Monocytes | 0 | 0 | 0 | 0.0263 | 0.0815 | 0 | 0 | 0 | 0.0099 | 0 | 0 | 0 |
| Myocytes | 0.0067 | 0.0096 | 0.0124 | 0 | 4.00E-04 | 0.0126 | 0.0107 | 0.011 | 0.0156 | 0.0068 | 0 | 0.0082 |
| NK cells | 0 | 0 | 0 | 0 | 0 | 0 | 0 | 0 | 0 | 0 | 0 | 0 |
| NKT | 0.001 | 0 | 0 | 0 | 0 | 0 | 0 | 0.0011 | 0 | 0 | 0 | 0 |
| Neurons | 9.00E-04 | 0.002 | 0.0016 | 0 | 0 | 2.00E-04 | 0.0011 | 0.0013 | 3.00E-04 | 6.00E-04 | 7.00E-04 | 4.00E-04 |
| Neutrophils | 5.00E-04 | 0 | 0 | 0.0194 | 0.0491 | 0.0062 | 0 | 0 | 0.0011 | 0 | 0 | 0 |
| Osteoblast | 0 | 0 | 0 | 0.0094 | 0 | 0.0048 | 0 | 0 | 0 | 0 | 0 | 0 |
| Pericytes | 0 | 0 | 0.0057 | 0.0637 | 0.0674 | 0.0259 | 0 | 0 | 0 | 0.0194 | 0.0058 | 0 |
| Plasma cells | 0.0024 | 0.001 | 0 | 0.0078 | 0 | 0.0021 | 0.0081 | 0.0059 | 8.00E-04 | 0.003 | 0 | 0 |
| Platelets | 0.0022 | 0 | 0 | 0 | 0 | 0 | 0.0069 | 0.0083 | 3.00E-04 | 0 | 0 | 0.0015 |
| Preadipocytes | 0 | 0 | 0 | 0.018 | 0 | 0 | 0 | 0 | 0 | 0 | 0 | 0 |
| Sebocytes | 0 | 0 | 0 | 0 | 0 | 0 | 0 | 0 | 0 | 0 | 0 | 0 |
| Skeletal muscle | 0.0148 | 0.0163 | 0.0152 | 0 | 0.0089 | 0.0063 | 0.0085 | 0.0182 | 0.0213 | 0.0115 | 0.0055 | 0.0052 |
| Smooth muscle | 0.0281 | 0.0608 | 0.078 | 0 | 0.0043 | 0.0399 | 0.0828 | 0.0305 | 0.0447 | 0.033 | 0.0481 | 0.0474 |
| Tgd cells | 0 | 0 | 0 | 0 | 0 | 0 | 0 | 0 | 0 | 0 | 0 | 0 |
| Th1 cells | 0.0032 | 0 | 0 | 0.0021 | 0 | 0 | 0 | 0.0075 | 0 | 0 | 0 | 0.0079 |
| Th2 cells | 0.0049 | 0 | 0 | 0 | 0 | 0 | 0.0257 | 0.013 | 0.0178 | 0.0014 | 0.0212 | 0.043 |
| Tregs | 0 | 0 | 0 | 0.0734 | 0.0082 | 0.0439 | 0.0017 | 0 | 0.0069 | 0.0031 | 0 | 0 |
| aDC | 0.0084 | 0 | 0 | 0.3088 | 0.1938 | 0.1709 | 0 | 0 | 0.0701 | 0.0044 | 0.0045 | 0 |
| cDC | 0.0028 | 0 | 0.0045 | 5.00E-04 | 0 | 0 | 0 | 0 | 0.0067 | 0.014 | 0.0087 | 0 |
| iDC | 0.0104 | 0 | 0.0087 | 0 | 0 | 0 | 0.0072 | 0.0251 | 0.0258 | 0.0162 | 0.0143 | 0 |
| ly Endothelial cells | 0 | 8.00E-04 | 5.00E-04 | 0 | 0.0024 | 0.0012 | 0 | 6.00E-04 | 7.00E-04 | 0 | 0.0033 | 0 |
| mv Endothelial cells | 0 | 3.00E-04 | 8.00E-04 | 4.00E-04 | 0.0071 | 0.0033 | 0 | 0 | 0 | 7.00E-04 | 0.0123 | 0.0023 |
| naive B-cells | 0 | 0 | 0 | 0 | 0 | 0 | 0 | 0 | 0 | 0 | 0 | 0 |
| pDC | 0 | 0 | 0 | 0.0115 | 0.0024 | 0.006 | 0 | 0 | 0.0028 | 0 | 0 | 0 |
| pro B-cells | 0.0019 | 0 | 0 | 0 | 0 | 0 | 0.0121 | 0.0103 | 0.0081 | 0 | 0 | 0.0011 |
| ImmuneScore | 0.0013 | 0 | 0 | 0.0482 | 0.0973 | 0.0143 | 0 | 0 | 0.0156 | 0.0033 | 0.0048 | 2.00E-04 |
| StromaScore | 0 | 0.0042 | 0.0016 | 0.0062 | 0.039 | 0.0025 | 0 | 0 | 0 | 0.0087 | 0.0196 | 1.00E-04 |
| MicroenvironmentScore | 0.0013 | 0.0042 | 0.0016 | 0.0544 | 0.1363 | 0.0168 | 0 | 0 | 0.0156 | 0.012 | 0.0244 | 3.00E-04 |

| | Mouse_Cont | Mouse_CT2 | Mouse_CT3 | Mouse_LPS1 | Mouse_LPS2 | Mouse_LPS3 | Mouse_3MC | Mouse_3MC | Mouse_3MC | Mouse_UV1 | Mouse_UV2 | Mouse_UV3 |
|---|---|---|---|---|---|---|---|---|---|---|---|---|
| Adipocytes | 0.002 | 0 | 0.0028 | 0 | 0 | 0 | 6.00E-04 | 0.0066 | 0.0062 | 1.00E-04 | 0 | 0 |
| Astrocytes | 0 | 0 | 0 | 0 | 0 | 0 | 0 | 0 | 0 | 0 | 0 | 0 |
| B-cells | 0 | 0 | 0 | 0 | 0 | 0 | 0.0085 | 0.0084 | 0.0035 | 0 | 0 | 0 |
| Basophils | 0 | 0 | 0 | 0.0816 | 0.0424 | 0.0677 | 0.0115 | 0 | 0.0049 | 0.0164 | 0.0214 | 0 |
| CD4+ T-cells | 0 | 0 | 0 | 0 | 0 | 0 | 0 | 0 | 0 | 0 | 0 | 0 |
| CD4+ Tcm | 0 | 0 | 0 | 0 | 0 | 0 | 0 | 0 | 0 | 0 | 0 | 0 |
| CD4+ Tem | 3.00E-04 | 0 | 0.0023 | 0 | 0 | 0 | 0.0136 | 0.0164 | 0.0331 | 0.0107 | 0.0096 | 0 |
| CD4+ memory T-cells | 0 | 0 | 0 | 0 | 0 | 0 | 0 | 0 | 0 | 0 | 5.00E-04 | 0 |
| CD4+ naive T-cells | 0 | 0 | 0 | 0 | 0 | 0 | 0 | 0 | 0 | 0 | 0 | 0 |
| CD8+ T-cells | 0 | 0 | 0 | 0 | 0 | 0 | 0 | 0 | 0 | 0 | 0 | 0 |
| CD8+ Tcm | 0 | 0 | 0 | 0 | 0 | 0 | 0 | 0 | 0 | 0 | 0 | 0 |
| CD8+ Tem | 0 | 0 | 0 | 0 | 0 | 0 | 0 | 0 | 0 | 0 | 0 | 0 |
| CD8+ naive T-cells | 0 | 0 | 0 | 0 | 0 | 0 | 0 | 0 | 0 | 2.00E-04 | 0.0037 | 0 |
| CLP | 0 | 5.00E-04 | 0 | 0 | 4.00E-04 | 0 | 0.0015 | 0 | 0 | 8.00E-04 | 0.0035 | 0.0013 |
| CMP | 0.0012 | 0.001 | 2.00E-04 | 0 | 0 | 0 | 3.00E-04 | 0.0017 | 0.0042 | 0 | 0 | 0 |
| Chondrocytes | 0 | 0 | 0 | 0 | 0 | 0 | 7.00E-04 | 0 | 0 | 0 | 0 | 0 |
| Class-switched memory B-cells | 0 | 0 | 0 | 0.0141 | 0.0032 | 0.014 | 0.0136 | 0.0168 | 0.0139 | 0 | 0 | 0 |
| DC | 0 | 0 | 0 | 0 | 0 | 0 | 0.0063 | 0.0064 | 0.0032 | 0.0074 | 0.0015 | 0.0012 |
| Endothelial cells | 0.0027 | 0 | 0.0039 | 0.0059 | 0.0022 | 0.0134 | 0 | 6.00E-04 | 0.0135 | 0 | 0 | 0 |
| Eosinophils | 0 | 0 | 0 | 0 | 0 | 0 | 0.0045 | 0.0057 | 0.0048 | 0 | 0 | 0 |
| Epithelial cells | 0.0041 | 0.0055 | 5.00E-04 | 0.0084 | 0.0141 | 0.0096 | 0.0012 | 4.00E-04 | 0 | 0.0179 | 0.0219 | 0.0144 |
| Erythrocytes | 0 | 0 | 0 | 0 | 0 | 0 | 0 | 0 | 0 | 0 | 0 | 0 |
| Fibroblasts | 0.0114 | 0.0061 | 0.0331 | 0.0411 | 0 | 0.0316 | 0.0593 | 0.0738 | 0.0641 | 0.0097 | 0.0164 | 0.015 |
| GMP | 0 | 0 | 0 | 0 | 0 | 0 | 0 | 0 | 0.0085 | 0.0046 | 0 | 0.0034 |
| HSC | 0.08 | 0.067 | 0.0564 | 0.0057 | 0 | 0.0044 | 0.0392 | 0.0357 | 0.1136 | 0.0084 | 0.0033 | 0.026 |
| Hepatocytes | 0 | 0 | 0 | 0.0027 | 0.0015 | 0.0023 | 0 | 0 | 0 | 3.00E-04 | 3.00E-04 | 1.00E-04 |
| Keratinocytes | 3.00E-04 | 3.00E-04 | 0 | 0.0014 | 0.003 | 0.0013 | 0 | 0 | 0 | 0.0051 | 0.0054 | 0.0048 |
| MEP | 0.0104 | 0.0057 | 0.0088 | 0.0188 | 0.0172 | 0.0181 | 0 | 0 | 0 | 0.0104 | 0.0192 | 0.0044 |
| MPP | 0 | 0 | 0 | 0 | 0 | 0 | 0 | 0 | 0 | 0 | 0 | 0 |
| MSC | 0.122 | 0.1173 | 0.1322 | 0.0119 | 0 | 0.0241 | 0 | 0.027 | 0.0585 | 0.014 | 0 | 0.0276 |
| Macrophages | 0 | 0 | 0 | 0.0184 | 0 | 0.0161 | 0.025 | 0.0234 | 0.0266 | 0 | 0 | 0 |
| Macrophages M1 | 0.0074 | 0.0046 | 0.0068 | 6.00E-04 | 0 | 0.0017 | 0.0287 | 0.0334 | 0.0283 | 0.0132 | 0.008 | 0.0062 |
| Macrophages M2 | 0 | 0 | 0 | 0.0073 | 0 | 0.0072 | 0.0209 | 0.0173 | 0.0155 | 0 | 0 | 0 |
| Mast cells | 0 | 0 | 0 | 0.0012 | 0.0017 | 0.0018 | 0.0044 | 0.0035 | 0.001 | 0 | 0 | 0 |
| Megakaryocytes | 3.00E-04 | 1.00E-04 | 0 | 0.0057 | 0.0028 | 0.0064 | 0 | 0 | 1.00E-04 | 0 | 0 | 0 |
| Melanocytes | 0.0013 | 9.00E-04 | 0.0014 | 0 | 0 | 0 | 0 | 7.00E-04 | 0.0019 | 2.00E-04 | 0 | 4.00E-04 |
| Memory B-cells | 0 | 0 | 0 | 0 | 0 | 0 | 0 | 0 | 0 | 0 | 0 | 0 |
| Mesangial cells | 0 | 0 | 0 | 0 | 0 | 0.0011 | 0 | 0 | 0 | 0 | 0 | 0 |
| Monocytes | 0 | 0 | 0 | 0.0102 | 0 | 0.0023 | 0.0162 | 0.0194 | 0.0204 | 0 | 0 | 0 |
| Myocytes | 0 | 0.003 | 0.0039 | 0.0029 | 0.0038 | 0.002 | 0.0023 | 0 | 0 | 0.0018 | 0 | 0.0016 |
| NK cells | 0 | 0 | 0 | 0 | 0 | 0 | 0 | 0 | 0 | 0 | 0 | 0 |
| NKT | 0.0161 | 0.0243 | 0.0658 | 0 | 0 | 0 | 0.0382 | 0.0182 | 0.0199 | 0.0107 | 0 | 0.0116 |
| Neurons | 0.0016 | 0.0016 | 0.0019 | 0 | 0 | 0 | 6.00E-04 | 4.00E-04 | 7.00E-04 | 2.00E-04 | 0 | 5.00E-04 |
| Neutrophils | 0 | 0 | 0 | 0.0786 | 0.0426 | 0.0689 | 6.00E-04 | 0 | 0 | 3.00E-04 | 0 | 0 |
| Osteoblast | 0 | 0 | 0 | 5.00E-04 | 0.0137 | 0 | 0.0053 | 0 | 0 | 0.0206 | 0.0209 | 0.0109 |
| Pericytes | 0.0121 | 2.00E-04 | 0.042 | 0.033 | 0.0212 | 0.0283 | 0 | 0 | 0 | 0.0289 | 0.0397 | 0.0148 |
| Plasma cells | 7.00E-04 | 0 | 0.0022 | 0.0225 | 0.024 | 0.0264 | 0.0031 | 0.0026 | 0.0114 | 0.0051 | 0 | 0 |
| Platelets | 0 | 0 | 0 | 0 | 0 | 0 | 0 | 0 | 0 | 0 | 0 | 0 |
| Preadipocytes | 0.0094 | 0.0174 | 0.004 | 0.0073 | 0 | 0.0058 | 0.0086 | 0.033 | 0.0341 | 0.0023 | 0 | 8.00E-04 |
| Sebocytes | 0 | 0 | 0 | 1.00E-04 | 0 | 0 | 0 | 0 | 0 | 6.00E-04 | 0.0011 | 6.00E-04 |
| Skeletal muscle | 3.00E-04 | 7.00E-04 | 0.005 | 3.00E-04 | 0.0012 | 0 | 0 | 4.00E-04 | 0.0023 | 0.0043 | 0 | 0.002 |
| Smooth muscle | 0.0898 | 0.0923 | 0.092 | 0.0206 | 0 | 0 | 0.0619 | 0.0344 | 0.0684 | 0.0636 | 0.0544 | 0.0838 |
| Tgd cells | 0 | 0 | 0 | 0 | 0 | 0 | 0 | 0 | 0 | 0 | 0 | 0 |
| Th1 cells | 0 | 0 | 0 | 0.0118 | 0.0128 | 0.0123 | 0 | 0 | 0 | 0.0058 | 0.0077 | 0.0028 |
| Th2 cells | 0 | 0 | 0 | 0 | 0 | 0 | 0 | 0 | 0.0037 | 0.0286 | 0.0783 | 0.0104 |
| Tregs | 0 | 0 | 0 | 0 | 0 | 0 | 0 | 0 | 0 | 0 | 0 | 0 |
| aDC | 0 | 0 | 0 | 0.0132 | 0 | 0.0344 | 0.0013 | 0 | 0.0334 | 0 | 0 | 0 |
| cDC | 0.0434 | 0.0286 | 0.034 | 0 | 0 | 0.001 | 0.0993 | 0.1296 | 0.0684 | 0.0365 | 0.0117 | 0.0221 |
| iDC | 0.1695 | 0.1345 | 0.1286 | 0.0048 | 0 | 0.0189 | 0.224 | 0.2812 | 0.1284 | 0.1806 | 0.0911 | 0.0843 |
| ly Endothelial cells | 0 | 0 | 0 | 0 | 0 | 0 | 0 | 0 | 0.0015 | 0 | 0 | 0 |
| mv Endothelial cells | 0 | 0 | 0 | 6.00E-04 | 3.00E-04 | 0.0033 | 0 | 0 | 7.00E-04 | 0 | 0 | 0 |
| naive B-cells | 0 | 0 | 0 | 0 | 0 | 0 | 0 | 0 | 0 | 0 | 0 | 0 |
| pDC | 0 | 0 | 0 | 0 | 0 | 1.00E-04 | 0 | 0 | 0 | 0 | 0 | 0 |
| pro B-cells | 0 | 0 | 0 | 0 | 0.0015 | 0 | 0 | 0 | 0 | 0 | 0.0041 | 0 |
| ImmuneScore | 0 | 0 | 0 | 0.0723 | 0.0295 | 0.0594 | 0.0437 | 0.0445 | 0.0397 | 0.0051 | 0.001 | 8.00E-04 |
| StromaScore | 0.008 | 0.0031 | 0.0199 | 0.0235 | 0.0011 | 0.0225 | 0.03 | 0.0405 | 0.0419 | 0.0049 | 0.0082 | 0.0075 |
| MicroenvironmentScore | 0.008 | 0.0031 | 0.0199 | 0.0958 | 0.0306 | 0.0819 | 0.0736 | 0.085 | 0.0816 | 0.01 | 0.0092 | 0.0083 |

**Dataset S4: Summary of enriched GO terms and pathways determined by Metascape.**

**Fig. 4A**

| GroupID | Category | Term | Description | -LogP | Symbols |
|---|---|---|---|---|---|
| 1_Summary | GO Biological Processes | GO:0031341 | regulation of cell killing | 6.066852715 | IL12B,IL12RB1,PGLYRP1,RIPK3,CLEC7A,SLAMF6,ITGAL,VCAM1,TNFSF9,TREML2,CCDC88B,SLAMF8,ALOX15,MMP12,FCRLB,SOX15,CD177,GPR55, CTSS,GPR171,IL18BP,BRCA1,IL2RG,NRG1 |
| 2_Summary | KEGG Pathway | hsa04060 | Cytokine-cytokine receptor interaction | 5.624808977 | CCR3,IL2RG,IL12B,IL12RB1,TNFRSF9,CCL22,TNFSF9,TNFRSF10B,ALOX15,GPR35,MMP12,VCAM1,IL18BP,BRCA1 |
| 3_Summary | GO Biological Processes | GO:0050900 | leukocyte migration | 5.36324153 | BDKRB1,F7,ITGAL,CCL22,VCAM1,TNFRSF10B,RIPK3,SLC7A11,SLAMF8,CD177 |
| 4_Summary | Canonical Pathways | M5885 | NABA MATRISOME ASSOCIATED | 5.354099137 | CTSS,F7,F10,NRG1,IL12B,MMP12,PLOD2,CCL22,TNFSF9,INSL6,CLEC7A,P4HA3 |
| 5_Summary | Reactome Gene Sets | R-HSA-5213460 | RIPK1-mediated regulated necrosis | 4.466772045 | TNFRSF10B,RIPK3,MLKL,NRG1,IL12B,CEMIP,PSRC1,IL2RG |
| 6_Summary | Reactome Gene Sets | R-HSA-1474244 | Extracellular matrix organization | 4.304360619 | CTSS,ITGAL,MATN1,MMP12,PLOD2,VCAM1,P4HA3,F7,F10 |
| 7_Summary | GO Biological Processes | GO:0046579 | positive regulation of Ras protein signal transduction | 4.280391268 | GPR35,NRG1,GPR65,GPR55,BDKRB1,CCR3,CEMIP,TMC8,SLAMF8 |
| 8_Summary | Canonical Pathways | M169 | PID INTEGRIN2 PATHWAY | 3.969876849 | F10,ITGAL,VCAM1,IGSF9B,CD177,ROBO3,CDHR1,TREML2,SLAMF6 |
| 9_Summary | Reactome Gene Sets | R-HSA-140877 | Formation of Fibrin Clot (Clotting Cascade) | 3.581700021 | F7,F10,CD177,ALOX15,NRG1,MMP12,SOX15,SLC7A11,ITGAL,TNFRSF10B,BDKRB1,KIFC1 |
| 10_Summary | GO Biological Processes | GO:0009268 | response to pH | 3.548836403 | CTSS,INSRR,GPR65 |

**Fig. S13A**

| GroupID | Category | Term | Description | -LogP | Symbols |
|---|---|---|---|---|---|
| 1_Summary | KEGG Pathway | hsa04115 | p53 signaling pathway | 4.554952749 | DDB2,MDM2,SESN2,SGK1,CPT1A |
| 2_Summary | GO Biological Processes | GO:0043255 | regulation of carbohydrate biosynthetic process | 4.112356404 | MST1,CD244,SESN2,CPT1A,PGGHG |

**Dataset S5: Naked mole-rats used in this study.**

| | ID | Sex | Treatment | Birth | Day of experiments | Use |
|---|---|---|---|---|---|---|
| *Chemical carcinogenesis by 3MC muscular injection* | | | | | | |
| *Naked mole-rat* | L40 | M | 3MC | 16.9.18 | 17.10.31 | Pathological study |
| *Naked mole-rat* | L41 | M | 3MC | 16.9.18 | 17.10.31 | |
| *Naked mole-rat* | L43 | M | 3MC | 16.9.18 | 17.10.31 | |
| *Naked mole-rat* | 2B4 | M | 3MC | 17.1.13 | 18.01.10 | Pathological study |
| *Naked mole-rat* | 2B7 | M | 3MC | 17.1.13 | 18.01.10 | |
| *Naked mole-rat* | 2B9 | M | 3MC | 17.1.13 | 18.01.10 | |
| *Naked mole-rat* | 2E11 | M | 3MC | 17.06.16 | 18.04.29 | Pathological study |
| *Naked mole-rat* | *2E14* | M | 3MC | 17.06.16 | 18.04.29 | |
| *Naked mole-rat* | 2E21 | M | 3MC | 17.06.16 | 18.04.29 | |
| *Chemical carcinogenesis by 3MC (41.6 mg/g of body weight) muscular injection* | | | | | | |
| *Naked mole-rat* | L77 | F | 3MC | 19.3.11 | 21.1.12 | |
| *Naked mole-rat* | L78 | F | 3MC | 19.3.11 | 21.1.12 | |
| *Naked mole-rat* | GH3 | M | 3MC | 19.1.10 | 21.1.12 | |
| *Naked mole-rat* | GH4 | M | 3MC | 19.1.10 | 21.1.12 | |
| *Naked mole-rat* | 2E25 | M | 3MC | 18.11.2 | 21.1.12 | |
| *Naked mole-rat* | 2E26 | M | 3MC | 18.11.2 | 21.1.12 | |
| *Naked mole-rat* | GH24 | M | 3MC | 19.12.11 | 21.1.13 | |
| *Naked mole-rat* | MG23 | M | 3MC | 19.10.11 | 21.1.13 | |
| *Naked mole-rat* | L100 | M | 3MC | 19.8.23 | 21.1.13 | |
| *Chemical carcinogenesis by 3MC subcutaneous injection* | | | | | | |
| *Naked mole-rat* | 2E8 | F | 3MC | 17.06.16 | 19.03.13 | |
| *Naked mole-rat* | 2E9 | F | 3MC | 17.06.16 | 19.03.13 | |
| *Naked mole-rat* | 2E12 | F | 3MC | 17.06.16 | 19.03.13 | |
| *Naked mole-rat* | C51 | M | 3MC | 18.07.08 | 19.03.13 | |
| *Naked mole-rat* | C52 | M | 3MC | 18.07.09 | 19.03.13 | |
| *Chemical carcinogensis by DMBA/TPA* | | | | | | |
| *Naked mole-rat* | 2D15 | M | DMBA/TPA | 18.9.14 | 19.06.28 | Pathological study |
| *Naked mole-rat* | 2D17 | M | DMBA/TPA | 18.9.14 | 19.06.28 | Pathological study |
| *Naked mole-rat* | L66 | M | DMBA/TPA | 18.9.24 | 19.06.28 | Pathological study |
| *Naked mole-rat* | L67 | M | DMBA/TPA | 18.9.24 | 19.06.28 | Pathological study |
| *Naked mole-rat* | C54 | M | DMBA/TPA | 18.7.8 | 19.06.28 | Pathological study |
| *Naked mole-rat* | C58 | M | DMBA/TPA | 18.7.8 | 19.06.28 | |
| *3MC injection 1 week* | | | | | | |
| *Naked mole-rat* | Y29 | F | 3MC | 17.12.25 | 18.11.27 | Pathologcal study and RNAsequencing |
| *Naked mole-rat* | C53 | M | 3MC | 18.7.8 | 19.04.05 | RNA-sequencing |
| *Naked mole-rat* | C32 | F | 3MC | 16.6.27 | 18.11.27 | Pathologcal study |
| *Naked mole-rat* | 2B13 | F | 3MC | 17.4.25 | 18.11.27 | Pathologcal study and RNA sequencing |
| *3MC injection 3 weeks* | | | | | | |
| *Naked mole-rat* | GH28 | M | 3MC 3 wk | 19.12.11 | 21.1.12 | Pathologcal study |
| *Naked mole-rat* | MG21 | F | 3MC 3 wk | 19.10.11 | 21.1.12 | Pathologcal study |
| *Naked mole-rat* | L102 | M | 3MC 3 wk | 19.8.23 | 21.1.12 | Pathologcal study |
| *DMBA treatment 24 hr* | | | | | | |
| *Naked mole-rat* | MG3 | M | DMBA 24 h | 19.2.3 | 21.5.25 | Pathologcal study |
| *Naked mole-rat* | BC11 | M | DMBA 24 h | 20.4.8 | 21.5.25 | Pathologcal study |
| *Naked mole-rat* | 2E31 | F | DMBA 24 h | 18.11.2 | 21.5.25 | Pathologcal study |
| *DMBA/TPA treatment 2 weeks* | | | | | | |
| *Naked mole-rat* | GH29 | M | DMBA/TPA 2 wk | 19.12.11 | 21.1.12 | Pathologcal study |
| *Naked mole-rat* | MG24 | M | DMBA/TPA 2 wk | 19.10.11 | 21.1.12 | Pathologcal study |
| *Naked mole-rat* | L101 | F | DMBA/TPA 2 wk | 19.8.23 | 21.1.12 | Pathologcal study |
| *LPS subcutaneous injection* | | | | | | |
| *Naked mole-rat* | C30 | M | LPS | 16.6.27 | 19.02.07 | Pathologcal study and RNA sequencing |
| *Naked mole-rat* | Y31 | N.D | LPS | 17.12.25 | 19.02.07 | Pathologcal study and RNA sequencing |
| *Naked mole-rat* | 2B8 | F | LPS | 17.1.13 | 19.02.07 | Pathologcal study and RNA sequencing |
| *UV Irradiation* | | | | | | |
| *Naked mole-rat* | Y30 | F | UV | 17.12.25 | 19.02.21 | Pathologcal study and RNA sequencing |
| *Naked mole-rat* | C46 | M | UV | 17.1.5 | 19.01.08 | Pathologcal study and RNA sequencing |
| *Naked mole-rat* | 2B18 | F | UV | 17.4.25 | 19.01.08 | Pathologcal study and RNA sequencing |
| *Healthy Sample* | | | | | | |
| *Naked mole-rat* | 2B15 | F | Control | 17.4.25 | 19.04.12 | Pathological study |
| *Naked mole-rat* | C33 | F | Control | 16.6.27 | 17.08.04 | Pathological study |
| *Naked mole-rat* | C34 | M | Control | 16.6.27 | 17.08.10 | Pathological study |
| *Naked mole-rat* | C31 | F | Control | 16.6.27 | 19.02.08 | Pathological study and RNA sequencing |
| *Naked mole-rat* | 2B10 | F | Control | 17.1.23 | 19.02.08 | Pathological study and RNA sequencing |
| *Naked mole-rat* | Y32 | M | Control | 19.12.25 | 19.02.08 | Pathological study and RNA sequencing |
| *LPS injection (intraperitoneal)* | | | | | | |
| *Naked mole-rat* | D30 | M | LPS | 16.8.16 | 18.09.20 | Pathologcal study |
| *Naked mole-rat* | 2E16 | M | LPS | 17.06.16 | 18.07.30 | Pathologcal study |
| *Naked mole-rat* | 2B23 | F | LPS | 17.4.25 | 18.09.20 | Pathologcal study |

#### Dataset S6: Primers used in this study.

| Gene/primer name | Primer | Sequcence (5'-3') |
|---|---|---|
| *Ripk3* (Mouse) | Forward | GAAATGGATTGCCCGAGGGA |
| | Reverse | GTGCTTGCCTCTCAGGACAT |
| *RIPK3* (NMR_CDS) | Forward | CTCATCCCGAGCACCACTTC |
| | Reverse | CCGCTTCTTCCCAGGTGACAA |
| *RIPK3* (3'UTR) | Forward | GCGGCCTGTAGGTGTTGAA |
| | Reverse | GTCAGTTGGGGCATAGCAGG |
| *Mlkl* (Mouse) | Forward | AGGAGGCTAACCAGCAGATAGA |
| | Reverse | GAGCATTGCTTCAGGGTTTTGT |
| *MLKL* (NMR) | Forward | ACCTGGGGACTAACTCTGCT |
| | Reverse | CCACATTCATGCAAACAGCCC |
| *Il6* (Mouse) | Forward | TCTATACCACTTCACAAGTCGGA |
| | Reverse | GAATTGCCATTGCACAACTCTTT |
| *IL6* (NMR) | Forward | GCTAGTCCTCCACGATGTCC |
| | Reverse | TTGCCTTTTCCTCCTCTAGGC |
| *ACTB/Actb* | Forward | AGACCTTCAACACCCCAGCCATGT |
| | Reverse | GGCCAGCCAGGTCCAGACGCAG |
| *GAPDH/Gapdh* | Forward | CTCCTGCGACTTCAACAGCAA |
| | Reverse | TACCAGGAAATGAGCTTGACA |
| *Ripk3* KO F | Forward | AGCGACACCTTGTGATCTCC |
| *Ripk3* KO R | Reverse | CTGGCCCAAGACAACCCTTA |
| *Ripk3* Wild F | Forward | GGAAAAGTCAGCCAATCCCG |
| *Ripk3* Wild R | Reverse | GCAAGACTAGAGCACACCCTC |
